# Structural insights into cobalamin loading and reactivation of human methionine synthase

**DOI:** 10.1101/2025.11.10.687659

**Authors:** Douglas S. M. Ferreira, Katie McLennan, Calum Diamond, Melanie Vollmar, Wasim Kiyani, D. Sean Froese, Jola Kopec, Henry J. Bailey, Rod Chalk, Arnaud Baslé, Jonathan M. Elkins, Jesse A Coker, Wyatt W. Yue, Thomas J. McCorvie

## Abstract

Human methionine synthase (MTR) is an essential enzyme of one carbon metabolism. Consisting of a catalytic N-half and a cobalamin binding C-half, MTR utilises this intricate organometallic cofactor in the methyl transfer from methyltetrahydrofolate to homocysteine producing methionine. Cobalamin loading into MTR, and its subsequent activation, requires methylmalonic aciduria and homocystinuria Type D (MMADHC) protein and methionine synthase reductase (MTRR), respectively. However, the molecular basis of cobalamin binding and activation of human MTR aided by MMADHC and MTRR remains unknown. Here, using cryo-electron microscopy, we determined structures of human MTR in its apo, and cobalamin bound states. Apo MTR adopts a conformation where the two halves of the enzyme act independently with the C-half posed to bind cobalamin. Binding of cobalamin and its activation causes conformational changes in MTR that result in a flexible catalytically active state. AlphaFold predictions, validated by interaction studies, show that MMADHC interacts with the C-half of apo MTR to facilitate cobalamin loading. Unexpectedly we found that MTRR interacts at two distinct sites within the C-half of MTR which may aid in activation. Collectively these findings lay the groundwork to uncover the mechanisms through how MMADHC and MTRR coordinate cobalamin loading and activation of human MTR.

## Introduction

Methionine synthase (EC 2.1.1.13; 5-methyltetrahydrofolate-homocysteine methyltransferase, MTR) forms a crucial node connecting the methionine and folate cycles in one-carbon metabolism. MTR catalyses the methylation of homocysteine (Hcy) using 5-methyltetrahydrofolate (MeTHF) as the methyl group donor, to generate methionine (Met) and tetrahydrofolate (THF)^1^. A key function of MTR is to ensure a supply of Met precursor to synthesise the global methyl donor *S*-adenosyl-L-methionine (SAM) involved in a myriad of epigenetic methylation reactions^2^. MTR also serves as an important regulator of the folate cycle which carries one-carbon units for purine and pyrimidine biosynthesis^3,4^. In humans, the primary circulating folate species is MeTHF^5^, and MTR is the only enzyme that can metabolise MeTHF to regenerate THF, the active form required for subsequent one-carbon transfer reactions and DNA synthesis^6^.

MTR is a ∼140 kDa monomeric protein comprising a catalytic N-half and a cofactor binding C-half (Fig. 1a). The N-half consists of two rigidly linked (Linker 1) TIM barrels, known as a homocysteine (Hcy) domain and a folate (Fol) domain, which exclusively bind the substrates Hcy and MeTHF, respectively. Connected to the N-half by a flexible linker (Linker 2), the C-half consists of a four-helical bundle Cap domain, a cobalamin binding domain (Cob), and a SAM binding activation domain (Act)^7^. Notably, MTR is one of only two human enzymes that uses cobalamin (Cbl) as a cofactor^7^.

**Fig. 1.**
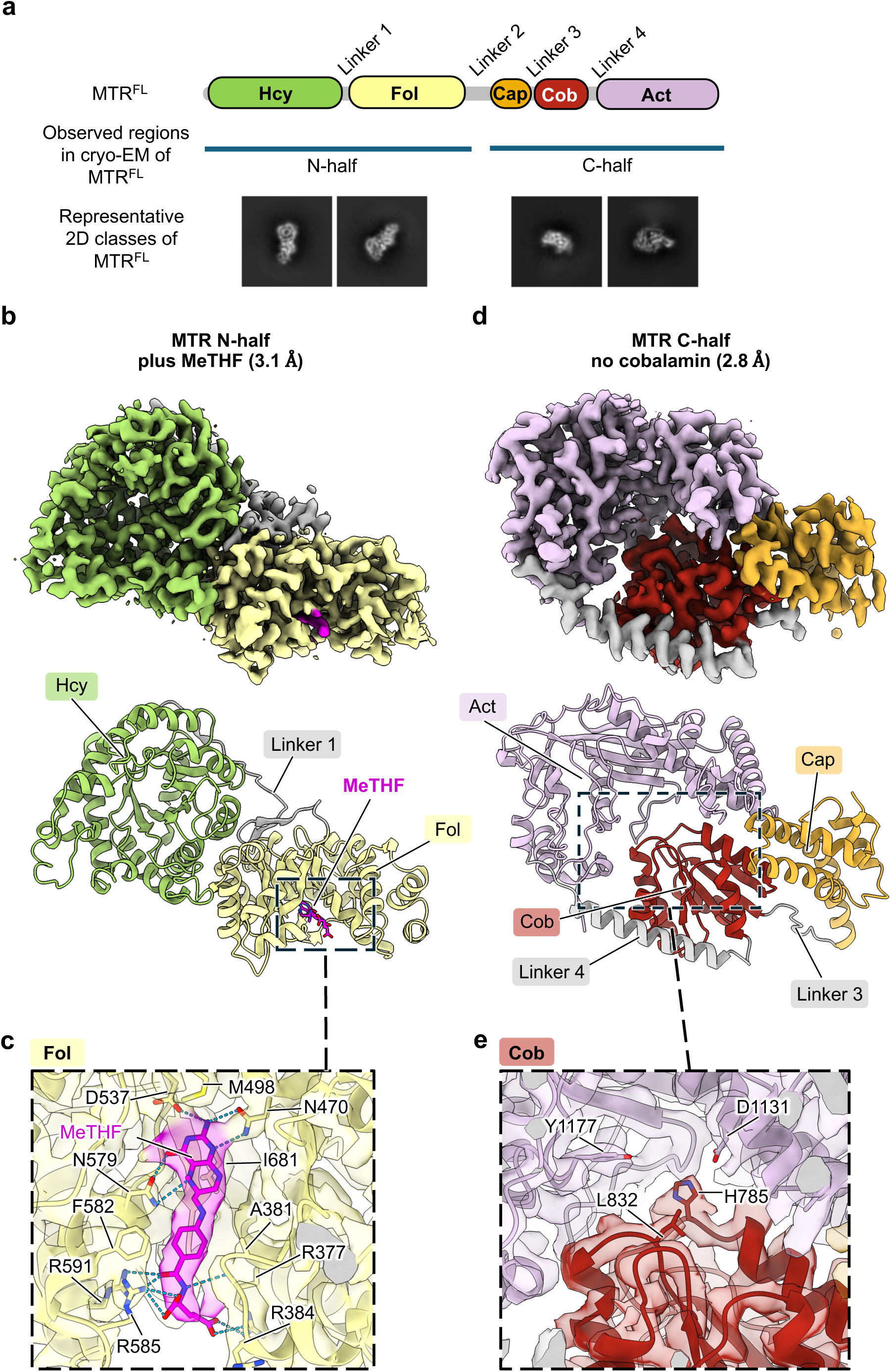
Cryo-EM analysis of MTR^FL^ bound to MeTHF. **a**, Cartoon schematic of the domain organization of MTR. Regions resolved of MTR^FL^ by cryo-EM are indicated. Representative 2D classes for each MTR map show the presence of just one half of the full-length enzyme. **b**, The 3.1 Å resolution map and model of N-half bound to MeTHF. Domains are coloured as in Fig. 1a and MeTHF is coloured magenta. **c**, Overlay of the map and model of the MeTHF binding pocket in the N-half of MTR^FL^, highlighting key residues and the ligand. **d**, The 2.8 Å resolution map and model of the C-half with no bound Cbl cofactor. Domains are coloured as in Fig. 1a. **e**, Overlay of the map and model of the cobalamin binding pocket in the C-half of MTR^FL^, highlighting key residues in the region and showing no detectable density corresponding to Cbl.

Cbl, also known as vitamin B_12_, is a large 1.3 kDa molecule consisting of a centrally bound hexacoordinated cobalt ion within a corrin ring, a ribose group tethered 5,6-dimethyl-benzimidazole (DMB) tail that can coordinate as a lower *α*-axial ligand (‘base-on’) and an upper β-axial ligand. This upper ligand can vary with the most common being a cyano or hydroxy group, whereas adenosyl or methyl group are the biological active forms in humans^7^. Within MTR the Cbl cofactor binds with the DMB group disengaged (‘base-off’) where in some instances a conserved histidine residue can bind as the lower axial ligand (His-off or His-on). During MTR’s catalytic cycle, the bound Cbl undergoes different cobalt oxidation and coordination states, such as alternating between the oxidised hexacoordinated methylcob(III)alamin form and the reduced tetracoordinated cob(I)alamin form. Cob(I)alamin is a strong nucleophile capable of extracting the methyl group from MeTHF, however it is also highly susceptible to oxidative inactivation, resulting in a cob(II)alamin form that renders MTR inactive. This necessitates the enzyme to enter a reactivation cycle where a reductive methylation process regenerates the functional methyl(III)cobalamin form from the inactive cob(II)alamin^8^. Previous structural studies using isolated domains of the bacterial orthologue, MetH, have suggested that for MTR to function large conformational changes are required in order for the Cob domain to interact with the Hcy and Fol domains within the rigid N-half during catalysis but also with the Act domain in the C-half during reactivation^9–11^.

Differentiating from its bacterial counterpart, human MTR has been shown to interact with three eukaryote-only proteins, namely methylmalonic aciduria and homocystinuria type C protein (MMACHC), methylmalonic aciduria and homocystinuria type D protein (MMADHC), and methionine synthase reductase (MTRR)^12^. MMACHC and MMADHC are known Cbl processing enzymes where MMACHC has Cbl reductase and dealkylase activities, to process various Cbl forms by removing the upper axial ligand to eventually produce cob(II)alamin^13^. Recent studies have shown that MMADHC can form a complex with MMACHC^14,15^, and can load Cbl onto MTR through forming a rare cobalt-sulfur bond donating a cysteine residue as one of the axial ligands^16,17^. MTRR is a multi-domain diflavin oxidoreductase that contains at its N-terminus a flavin mononucleotide (FMN) binding domain, flexibly linked to a C-terminal module that contains of a connecting (Conn) domain, a flavin adenine dinucleotide (FAD) binding domain, and a nicotinamide adenine dinucleotide phosphate (NADPH) binding domain^18^. MTRR plays a key role in reactivating MTR for catalysis, working in concert with the SAM-bound Act domain of MTR. MTRR donates electrons from NADPH to reduce and methylate the inactive cob(II)alamin bound to MTR, to produce functional methylcob(III)alamin^19,20^. This is a multistep sequential process where electrons from NADPH are initially transferred to FAD and then FMN before ultimately being transferred to the bound Cbl^7^. In bacteria this function is carried out by two separate proteins, flavodoxin and flavodoxin reductase^21^. Significantly, the essential roles of MTR and its interaction partners in human health are stressed by reported pathogenic mutations in their corresponding genes, that are associated with a collection of rare inherited metabolic disorders^22^. Altered Cbl and folate metabolism is also known to be linked to an increased risk of neural tube defects^23^.

Little is known on how human MTR binds Cbl and if it shares the same conformational flexibility as its bacterial orthologues. Even less is known on how human MTR interacts with its eukaryote specific partner proteins that support its functionality. To address these questions, we have used cryo-electron microscopy (cryo-EM) together with AlphaFold 3 (AF3) structure predictions, to investigate how human MTR binds Cbl and how it interacts with its functional protein partners MMACHC, MMADHC, and MTRR.

## Results

### Human MTR can be purified without bound cobalamin

We expressed and purified apo full-length human MTR (MTR^FL^) with a C-terminal His_6_ and FLAG tag from insect cells. Though milligrams of recombinant protein could be obtained, size-exclusion gel filtration demonstrated that the majority of MTR^FL^ formed aggregates with some monomeric protein (Supplementary Fig.1a). Negative stain electron microscopy (NS-EM) confirmed the presence of homogenous and monodisperse particles from the monomeric peak (Supplementary Fig. 1b). The molecular weight was determined to be 142.69 kDa by native mass-spectrometry, agreeing with the calculated molecular weight of 143.85 kDa (with tag) and confirming the purified monomeric protein was in the apo form without any detectable bound Cbl ligands (Supplementary Fig.1c) as previously reported^24,25^.

Using an absorbance-based activity assay^26^ monomeric MTR^FL^ was determined to be catalytically active (Supplementary Fig. 1d). To further assess functionality, we turned to thermal shift experiments testing MTR^FL^ along with purified MTR^N-Half^ (Hcy and Fol domains) and MTR^C-Half^ (Cap, Cob, and Act domains) constructs. Here we determined the melting temperature (T*_m_*) of these constructs as purified along with being in the presence of relevant substrates and different cobalamins. In these experiments, MTR^FL^ was stabilised by MeTHF with a 3.3 ± 0.3 °C melting temperature increase (ΔT*_m_*) (Supplementary Fig. 1e, g). MeTHF also induced a ΔT*_m_* increase of 4.8 ± 0.4 °C for MTR^N-Half^, whereas no statistically significant change in T*_m_* was observed for MTR^C-Half^ as expected (Supplementary Fig.1e, g). In contrast, both Hcy and SAM showed no T*_m_* changes for any of the three constructs (Supplementary Fig. 1g). We then tested hydroxycobalamin (HOCbl), cyanocobalamin (CNCbl), and methylcobalamin (MeCbl) against the three constructs. As expected, no T*_m_* changes were observed for MTR^N-Half^, whereas all three Cbl ligands stabilised both MTR^FL^ and MTR^C-Half^ with MeCbl eliciting the largest T*_m_* shifts (Supplementary Fig. 1f, g). Overall, these experiments demonstrate that we can successfully purify functional apo full-length and constructs corresponding to each half of human MTR.

### Apo MTR is in a reactivation conformation for cobalamin loading

Having successfully purified apo MTR^FL^ and encouraged by our NS-EM results, we attempted to determine a high-resolution structure of this as-purified, Cbl free state by cryo-EM. Initial screening resulted in 2D classes representing only the C-half of MTR likely in the reactivation conformation where the Act domain is engaged with the Cob domain (Supplementary Fig. 2a). Unfortunately, after many attempts we could not produce conditions that yielded reliable good 2D classes representing the entirety of full-length MTR, likely due to instability and air-water denaturation. To overcome this, we screened and collected data in the presence of MeTHF, reasoning that it would stabilise MTR^FL^ as found in our thermal shift experiments (Supplementary Fig 1e, g). Initial screening resulted in separate 2D classes of both halves of MTR (Supplementary Fig. 2b). Despite multiple processing strategies no classes showed the concurrent presence of both halves of MTR. A larger dataset was then collected which allowed us to determine a 3.1 Å resolution structure of the 70 kDa N-half bound to MeTHF and a 2.8 Å resolution structure of the 68 kDa apo C-half (Fig. 1, Supplementary Fig. 3, 9, 10).

The N-half structure of MTR^FL^ confirmed that the Hcy and Fol domains are TIM barrel folds with the short Linker 1 (Pro347 to Phe371) that tightly locks the domains side by side (Fig. 1b). This arrangement implies that large conformational changes are required for the Cob domain to interact with the N-half domains during catalysis as has been suggested for bacterial MetH^7,9^. The Hcy domain showed weak density for a disulfide bond between Cys260 and Cys323 with no other apparent density for substrate (Fig 1b, Supplementary Fig. 4a). On the other hand, the Fol domain showed clear density for the MeTHF ligand (Fig. 1c) making interactions with the conserved residues Asn470, Asp537, and Asn579 through its pterin ring containing the methyl group. Additionally, the p-aminobenzoic acid (PABA) side chain of MeTHF forms hydrogen bonds with the residues Arg384, Arg585, and Arg591 (Fig. 1c). This MeTHF bound structure is highly similar to our previously deposited 2.7 Å crystal structure of the MTR^N-Half^ construct alone bound to THF (PDB 4CCZ), showing that the binding of MeTHF is near identical to THF, with a RMSD of 0.68 Å (Supplementary Fig. 4a). Comparing our human N-half cryo-EM structure to the reported N-half sections from *Thermus filiformis* (PDB 8G3H) and *Thermus thermophilus* HB8 (PDB 8SSC) MetH structures demonstrates how this tight arrangement between the Hcy and THF domains is evolutionarily conserved (Supplementary Fig. 4b)^24,27^.

The Cbl free C-half structure of MTR^FL^ was found to be in a conformation where the Cap domain is displaced (Cap-off) from the Cob domain, allowing free access to the Cbl binding pocket and engagement with the Act domain (Fig. 1d). No density was apparent for the Cbl cofactor as expected (Fig. 1e). This domain arrangement is similar to that reported in crystal structures of the isolated C-half from bacteria MetH bound to cob(II)alamin, which has been suggested to be the reactivation conformation productive for reductive methylation^10,28,29^ (Supplementary Fig. 4c). Structural alignment against the *Escherichia coli* C-half structure bound to cob(II)alamin (PDB 3BUL) and the apo C-half section from *T. thermophilus* (PDB 8SSC) shows only minor differences in the relative position of the Cap to the other domains (Supplementary Fig. 4c). The expected lower axial ligand for the Cbl cofactor, His785, is in a His-off conformation and the corresponding histidine sidechain adopts variable positions in the bacteria MetH structures of the His-off state (Fig 1e, Supplementary Fig. 4c). Tyr1177, which promotes the tetracoordinated cob(I)alamin form during MTR reactivation^29^, is positioned 3.7 Å closer to the putative position of the cobalt atom, when compared to the *apo* MeTH structure from *T. thermophilus* (Supplementary Fig. 4c)^24^.

### Binding of HOCbl to MTR causes local structural changes

The conformation of the C-half of MTR^FL^ suggests that the apo enzyme forms this “reactivation-like” conformation before the binding of free Cbl from solution or Cbl loading assisted by MMADHC. Therefore, to determine the conformational changes upon initial binding of Cbl, we collected cryo-EM data of MTR^FL^ in the presence of MeTHF and HOCbl. SAM was also added in an attempt to determine what conformational changes could occur before methylation of the bound Cbl. As with the MeTHF dataset, only separate 2D classes of the N-half and C-half alone could be determined, suggesting their independence when MTR^FL^ is in the presence of these ligands. Processing of the N-half particles generated a 2.9 Å resolution structure (Supplementary Fig. 5, 9, 10), identical to that with MeTHF alone (RMSD 0.356 Å) (Supplementary Fig. 6a).

Processing of the C-half particles revealed two states with an overall conformation as the Cbl free form (Supplementary Fig. 5, 9, 10). Inspection of the Cob domain in both states showed clear density for the bound cofactor, HOCbl (Fig. 2a, b, Supplementary Fig. 6b); however, the density for SAM in the Act domain was discontinuous and did not allow for modelling, suggesting flexibility or low occupancy of this ligand (Supplementary Fig. 6c). The first state, determined to a resolution of 2.9 Å, is representative of the majority of the C-half particles. In this state, density was apparent for HOCbl bound in a cob(II)alamin, base-off, His-off state (Fig. 2a, Supplementary Fig. 6b). Density for the dimethylbenzimidazole (DMB) tail bound in the pocket of the Cob domain was clear, interacting with conserved residues Ser830 and Gly860 (Supplementary Fig. 6b). The corrin ring showed less apparent density, suggesting flexibility with Leu832 in the location of the lower axial ligand (Fig. 2c, Supplementary Fig. 6b).

**Fig. 2.**
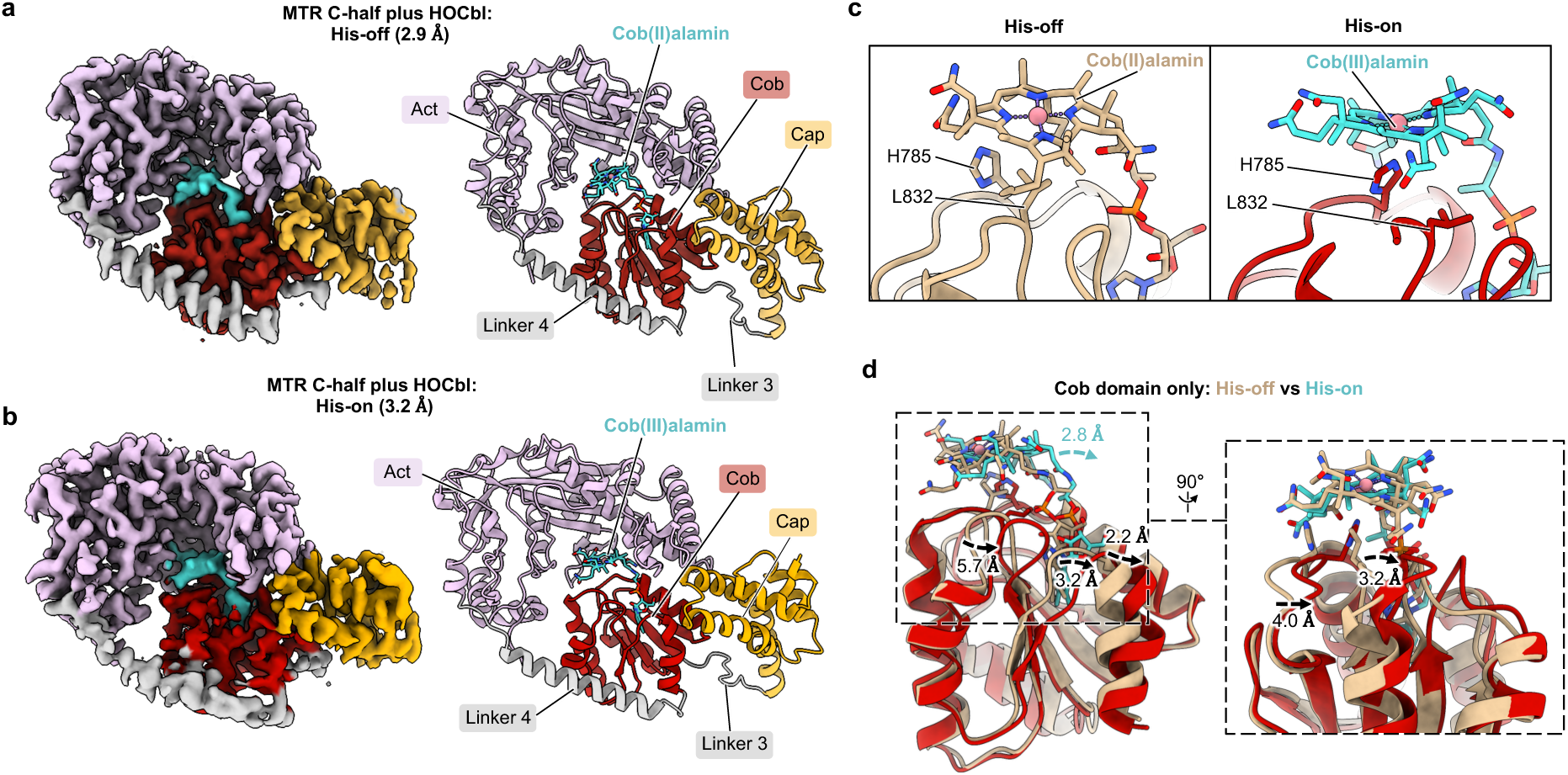
Cryo-EM analysis of the C-half of MTR^FL^ bound to HOCbl. **a**, The 2.9 Å resolution map and model of the C-half of MTR^FL^ bound to HOCbl in a His-off state. Domains are coloured as in Fig. 1a and HOCbl is coloured cyan. **b**, The 3.2 Å resolution map and model of the C-half of MTR^FL^ bound to HOCbl in a His-on state. Domains are coloured as in Fig. 1a and HOCbl is coloured cyan. **c**, Positioning of HOCbl and His785 in the His-off state (left, beige) and His-on state (right, cyan). **d**, Comparison of the movements of HOCbl and the Cob domain between the His-off and His-on states. The HOCbl and Cob domain in the His-off state are shown in beige, while HOCbl in the His-on state is shown in cyan, and the Cob domain in the His-on state is shown in red.

The second state was determined to a resolution of 3.2 Å (Fig. 2b, Supplementary Fig. 6b) and is representative of the minority of the particles. Similar to the His-off state described above, density was apparent for HOCbl, however its corrin ring has shifted by 2.8 Å (Fig. 2c, d) and is now in a His-on state, where His785 becomes the lower axial ligand (Fig. 2c). Furthermore, conformational changes are apparent in the Cob domain when comparing the His-off and His-on structures (Fig. 2d). These changes occur throughout the entire Cob domain, involving a shift away from the Act domain during the transition from the His-off to the His-on state. The density of the helical Linker 4 between the Cob and Act domains is also less defined (Fig. 2b), suggesting a degree of flexibility in this state.

In the C-half maps of both states, the Cap domain showed less detail than the other domains, suggesting heterogeneity possibly due to flexibility. Interested in analysing this further, we turned to 3D variability analysis (3DVA)^30^ where we compared the apo C-half map against the Cbl bound C-half map with both His-off and His-on particles pooled together (Supplementary Fig. 7, Supplementary Movies 1-4). For both the apo and Cbl bound states, flexing of the Cap domain was evident where it moves towards the Act domain. Simultaneously, the Cob domain appears to be moving away from the Act domain (Supplementary Fig. 7a, b, Supplementary Movies 1, 3). Other movements were observed in the Cbl bound state where the Cbl cofactor transitions from His-off to His-on correlating with larger movements of the Cob domain away from the Act domain (Supplementary Fig. 7b, Supplementary Movie 3) along with some rotation towards the Cap (Supplementary Fig. 7d, Supplementary Movie 4). We believe that this displacement of the Cob domain is representative of the conformational changes that occur when MeCbl is formed via reductive methylation and the subsequent change from a five-coordinate His-off Cap-off state to a six-coordinate His-on Cap-on state^31^. As no electron source was added to our sample, reductive methylation cannot occur, thereby allowing us to trap a small percentage of the MTR protein bound to HOCbl in this putative prelude state (His-on, Cap-off) before entering the catalytic cycle, similar to what has been suggested for a crystal structure of the His-on HOCbl bound state of the C-half from E*. coli* (PDB 3IV9)^29^.

### Binding of MeCbl remodels the C-half of MTR^FL^

After Cbl loading and reductive methylation aided by MTRR, MTR is activated and transforms into a six-coordinate His-on MeCbl-bound conformation, poised to begin the catalytic cycle^32^. Recent studies have suggested that *T. filiformis* and *T. thermophilus* MetH enters a so-called “resting” or “gating” state in the presence of Cbl and without substrates. In this state the N-half (Hcy and Fol domains) stably interacts with the Cap and Cob domains of the C-half in a Cap-on and a disengaged Act domain conformation^27,33^. Interested in capturing a similar Cap-on state of human MTR without substrate, we prepared MTR^FL^ in the presence of MeCbl for cryo-EM analysis. MeCbl has been shown to readily bind to the Cob domain from solution^17^ and also stabilised the protein in our thermal shift experiments (Supplementary Fig. 1e, g).

Preliminary screening of MTR^FL^ in the presence of MeCbl resulted in discernible 2D classes of the 70 kDa N-half only, and unlike in our previous datasets no apparent 2D classes could be determined corresponding to the C-half. However, as this sample of MTR^FL^ lacked any substrates, a larger dataset was then collected that resulted in a 2.7 Å resolution structure of the apo N-half of MTR^FL^ (Fig. 3b, Supplementary Fig. 8-10). Comparing this structure with our MeTHF- and THF-bound structures shows little conformational changes in both Hcy and Fol domains (Fig. 3c). No other domains were present in the map, and despite further processing no other states containing the N-half could be found unlike what has been reported for structures of thermophilic MTR homologues^27,33^. In these structures a loop located in the Fol domain changes conformation due to the absence of folate, and was suggested to play a role in altering the conformation of Linker 2 which connects the two halves of the enzyme, resulting in the “resting/gating” state^27,33^. However, unlike thermophilic MetH, the folate sensing loop in human MTR (residues 613-619) shows no structural changes due to the absence of folate and is positioned in a similar manner as the ligand bound states (Fig. 3d).

**Fig. 3.**
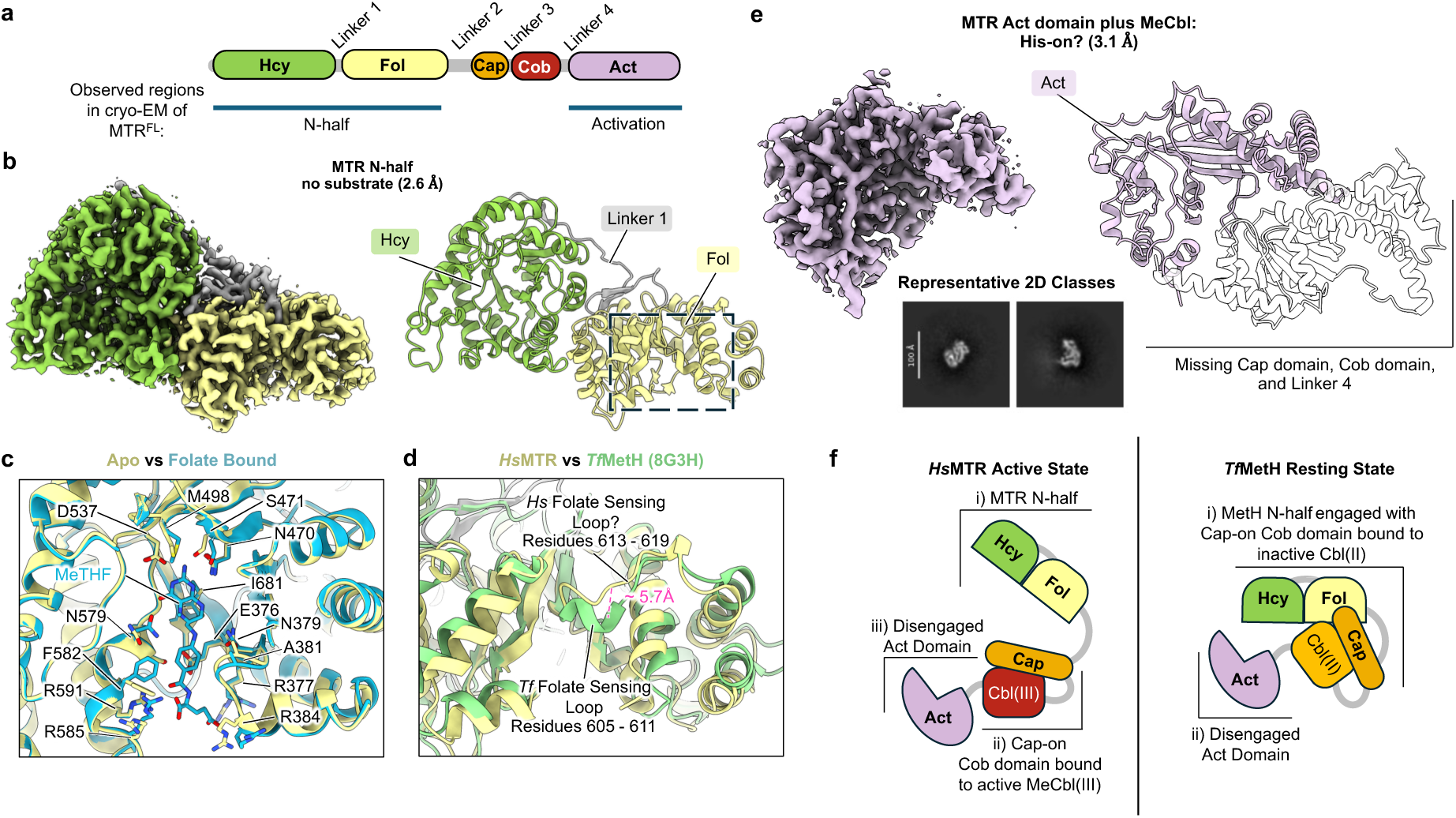
Cryo-EM analysis of MTR bound to MeCbl. **a**, Cartoon schematic of the domain organization of MTR. Domains resolved by cryo-EM are indicated. **b**, The 2.6 Å resolution map and model of the N-half of MTR^FL^ in the apo state. Domains are coloured as in Fig. 3a. **c**, Structural alignment of our apo (yellow) and folate bound (cyan) structures focused on the folate binding site. Little conformational changes occur due to folate binding with only a slight shift away from the active site for the small loop containing Ala381. **d**, Structural alignment of our apo N-half structure (yellow) and thermophilic MetH structure (8G3H, light green) focused on the folate binding site. Unlike the position of residues 605-611 in thermophilic MetH, corresponding to a putative folate sensing loop, no structural changes occur in residues 613-619 in human MTR due to the absence of folate. **e**, The 3.1 Å resolution map and model of the MTR^FL^ Act domain. The 2D classes shown are representative of the Act domain as observed in the cryo-EM data analysis. In the model the missing linkers, Cap, and Cob domains are also shown as a transparent model arranged as found in our C-half structures. **f**, Proposed domain arrangements as cartoon representations of MeCbl-bound active human MTR and thermophilic HOCbl-bound inactive MetH, without substrate. In human MTR MeCbl binding results in a conformation with the N-half, Cap-Cob domains, and Act domain acting as three independent modules. In contrast, HOCbl binding to thermophilic MetH results in a two-module conformation with the N-half plus Cap-Cob domains, and the Act domain acting independently.

The lack of a C-half structure from this dataset could have been due to a combination of intrinsic flexibility and air-water denaturation. In theory the binding of MeCbl should induce large conformational changes in the C-half, shifting MTR into a catalytical competent state and displacing the Act domain from the Cob domain to allow interactions of the bound Cbl with the N-half domains and Cap during catalysis. Further careful processing of our MeCbl-bound dataset revealed classes equating to the Act domain of MTR^FL^ (≈ 38 kDa), showing its disengagement from the Cob domain (Fig. 3e). Despite its small size, further processing resulted in a 3.1 Å resolution map (Fig. 3e, Supplementary Fig. 8-10). Comparison of this Act domain-alone structure with the Act domain in the context of our C-half structures (i.e. together with Cob and Cap domains) shows little structural changes within the Act domain (Supplementary Fig. 11a). Comparison to the crystal structure of an isolated Act domain construct (PDB 2O2K)^34^ showed that only a few residues in flexible regions displayed any significant conformational differences, likely resulting from crystal packing (Supplementary Fig. 11b). Therefore, we interpret that our MeCbl-bound MTR^FL^ dataset is likely in a His-on, Cap-on state consisting of three flexible independent modules, namely the N-half (≈ 70 kDa), the Cap and Cob domains (≈ 28 kDa), and the Act domain (≈ 38 kDa). Despite considerable effort in data processing, we failed to find any 2D classes equating to the MeCbl-bound Cap-Cob domains, likely due to its small size and its position between the N-half and Act domain. Indeed, inspecting the 2D classes and low pass filtering our Act domain map show the presence of blurry low-resolution density at the expected location of these two domains, suggesting their presence (Supplementary Fig. 11c, d). Although the difference between our MeCbl-bound structure and the reported structures of thermophilic MetH bound to inactive Cbl^27,33^ forms may be due to variations in sample preparation, it suggests that MTR can adopt more conformations than has been previously suggested (Fig. 3f).

### MMADHC, not MMACHC, binds MTR directly to load cobalamin

The cobalamin chaperone enzymes MMACHC and MMADHC can form a heterodimer in the presence of Cbl^14^, and previous studies have implied a stable complex of MTR with MMACHC and MMADHC^12,35^. Additionally, Cbl transfer from MMADHC to MTR has been monitored through size-exclusion chromatography^17^. However, no structural insights into their complex formation have been determined experimentally. Therefore, we turned to AlphaFold3 (AF3)^36^ modelling to predict how both MMACHC and MMADHC could interact with MTR to load Cbl.

AF3 predicted modelling of the MTR binary complexes was initially carried out using the sequences of the full-length proteins. This was then followed by using only the sequences of the N- and C-halves of MTR to confirm which domains were involved in protein-protein interactions. AF3 models for the MTR^FL^-MMACHC^FL^, MTR^N-Half^-MMACHC^FL^, and MTR^C-Half^-MMACHC^FL^ complexes demonstrated random spatial positions of MMACHC on MTR (Supplementary Fig 12, 14a, b). This combined with their low ipTM scores^36^ (∼ 0.2), which is a confidence measure of the accuracy of predicted complexes, suggest that MMACHC may not form a complex with MTR. In contrast the AF3 models for the MTR^FL^-MMADHC^FL^ complex (Fig. 4a, Supplementary Fig. 13) and the MTR^C-Half^-MMADHC^FL^ subcomplex (Fig. 4b, Supplementary Fig. 14d) resulted in highly similar and consistent predicted models with correspondingly moderate to high ipTM scores^36^ above 0.5. On the other hand, the MTR^N-Half^-MMADHC^FL^ subcomplex showed random spatial positions of each protein with a corresponding low ipTM score suggesting that MMADHC does not interact with either the Hcy or Fol domains of MTR (Supplementary Fig. 14c).

**Fig. 4.**
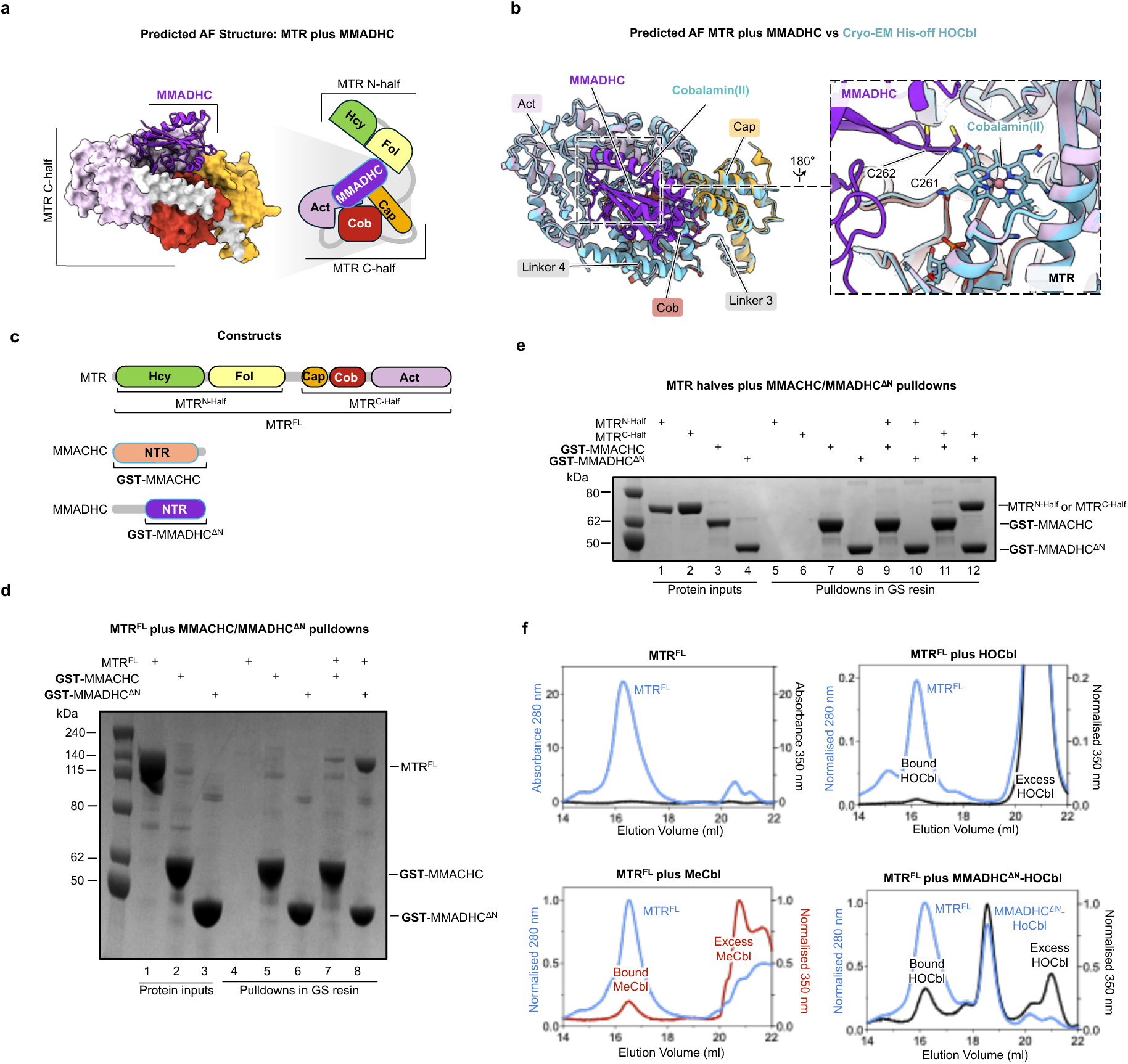
MMADHC forms a complex with MTR to load cobalamin. **a**, Representative predicted AF3 model of the MTR-MMADHC complex. The arrangement of the full-length complex is shown as a cartoon depicted with the domain colours as shown in Fig. 4a. The zoom-in cartoon shows only MTR^C-Half^ with MMADHC; MMADHC is a ribbon model whereas MTR is shown as a surface model. **b**, Structural overlay of the predicted MTR^C-Half^-MMADHC AF structure against our cryo-EM derived model of His-off MTR^C-Half^. The zoom-in panel focuses on the MTR-MMADHC interface showing the relative location of the functionally important MMADHC residues C261 and C262. **c**, Constructs and domain arrangements of MTR, MMACHC, and MMADHC used in cobalamin transfer and pulldown assays. Affinity tagged constructs are indicated. **d**, GS resin affinity pulldown of MTR^FL^ against GST-MMACHC and GST-MMADHC^ΔN^. Lanes 1–3 represent the protein inputs used for the assay. Lanes 4–6 represent the elution fractions of each individual protein incubated with the resin to verify their ability to bind the resin. Lanes 7 and 8 represent the pulldown of MTR^FL^ with GST-MMACHC and the pulldown of MTR^FL^ with GST-MMADHC, respectively. **e**, GS resin affinity pulldown of MTR^N-Half^ and MTR^C-Half^ against the GST-MMACHC and GST-MMADHC^ΔN^. Lanes 1–4 represent the protein inputs used for the assay. Lanes 5–8 represent the elution fractions of each individual protein incubated with the resin to verify their ability to bind the resin. Lanes 9 and 10 represent the pulldown of MTR^N-Half^ with GST-MMACHC and GST-MMADHC, respectively. Lanes 11 and 12 represent the pulldown of MTR^C-Half^ with GST-MMACHC and GST-MMADHC, respectively. **f**, Size-exclusion chromatography of cobalamin loading assay showing MTR^FL^, MTR^FL^ plus HOCbl, MTR^FL^ plus MeCbl, and MTR^FL^ plus HOCbl-loaded MMADHC^ΔN^ monitored at 280 nm (blue) and 350 nm (black for HOCbl or red for MeCbl).

In both sets of predictions with either MTR^FL^ or MTR^C-Half^, MMADHC is straddled above the Cob domain of MTR in the reactivation conformation, forming potential interactions with all three domains of the C-half of MTR (Fig. 4a, b, Supplementary Fig. 13, 14a, b). This interface results in the loop aa 255-266 from MMADHC to point towards the Cbl-binding pocket in the Cob domain of MTR. This loop harbours the residues Cys261 and Cys262, which are responsible for the binding of Cbl and its loading into MTR^17^. Superimposing the predicted models with our HOCbl-bound C-half structure of MTR^FL^ and the crystal structure of MMADHC coordinating a thiolato-cobalamin ligand (PDB 6X8Z)^37^ suggests that some conformational changes are required for Cbl loading. The Cbl cofactor needs to move 4.3 Å towards the binding pocket within the Cob domain of MTR when being loaded by MMADHC (Supplementary Fig 15).

To verify MTR’s predicted lack of interaction with MMACHC and its predicted positive interaction with MMADHC we performed affinity-based pulldowns using MTR^FL^ against a GST fusion of MMADHC^ΔN^ (MMADHC lacking the first 123 residues that were predicted to be disordered) and a GST fusion of full-length MMACHC (Fig. 4c, d, Supplementary Fig. 16a). These pulldowns show that MMACHC weakly interacts with MTR^FL^ (lane 7) whereas MMADHC appears to strongly interact (lane 8). To further confirm which halves of MTR are involved we carried out similar affinity-based pulldowns using MTR^N-Half^ and MTR^C-Half^ constructs (Fig. 4c, e). These pulldowns confirmed the weak, to potentially absent, interaction of MMACHC against the MTR^N-Half^ and MTR^C-Half^ constructs (Fig. 4e, Supplementary Fig. 16a), in which no positive pulldowns were observed (lanes 9, 11). Similar to previous published analyses on a C-terminal construct of MTR^17^, we observed that MMADHC^ΔN^ interacts with MTR^C-Half^ (lane 12) but not with MTR^N-Half^ (lane 10). Overall, these collection of pulldowns are consistent with their corresponding AF3 predictions.

We next used size-exclusion gel filtration to determine the loading of HOCbl onto MTR^FL^ by MMADHC^ΔN^ (without a GST tag), through monitoring absorbance at 350 nm, indicative of the presence of Cbl. MTR^FL^ displays no absorbance at 350 nm, indicating that it is not bound to cobalamin (Fig. 4f upper left panel) agreeing with our native mass spectrometry measurements (Supplementary Fig. 1c). In the absence of MMADHC, low amounts of HOCbl binds MTR (Fig. 4f upper right panel) whereas MeCbl binds more readily (Fig. 4f lower left panel), agreeing with a previous analysis on a C-terminal construct of MTR^17^. When preincubated with MMADHC^ΔN^, HOCbl is loaded into MTR^FL^ through MMADHC^ΔN^, suggesting a direct interaction (Fig 4f lower right panel). Together our pulldown and Cbl loading experiments suggest that MTR not only interacts with MMADHC for Cbl loading but also appears to interact with MMADHC when Cbl is not present.

### MTRR interacts with MTR through multiple domains

For the reactivation cycle, MTR requires the diflavin oxidoreductase MTRR to reduce the bound cob(II)alamin for methylation by SAM^19^. Beyond reports that the FMN domain of MTRR can interact with the Act domain of MTR^38^, little is known on how MTRR interacts with MTR to transfer electrons from NADPH. To gain structural insights into their interactions we turned to AF3 to predict possible models of the MTR-MTRR complex.

AF3 modelling of the complex between MTR^FL^ and MTRR^FL^ produced predicted models with moderate ipTM scores^36^, in which the MTR N-half location varies between the individual predictions and the MTR C-half adopts a reactivation conformation (Fig. 5a, Supplementary Fig 17). Consistent across the five predicted structures is the relative location of MTRR where the MTRR FMN domain is modelled to sit atop the MTR Cob domain and appears to interact with the Act domain (Fig. 5a). This agrees with previous biophysical studies demonstrating a physical interaction between the MTRR FMN domain and MTR Act domain^38^. To our surprise, the models reveal that in addition to the FMN domain, the other regions of MTRR (i.e. Conn domain, FAD domain, and NADPH domain; referred thereafter as MTRR ΔFMN) can also interact with MTR which was consistent across each prediction. This interaction occurs through two interfaces of MTRR ΔFMN, one involving the Conn domain that interacts with Linker 4 of MTR, and the other involving the NADPH domain that interacts with the MTR Cob domain. (Fig. 5a, 6a, Supplementary Fig. 17). This previously unidentified interaction was also consistently replicated in the AF3 modelling of complexes between the MTR C-half and the MTRR ΔFMN (Supplementary Fig. 18a). On the contrary, no consistent interactions with MTRR were predicted when the human MTR C-half sequence is replaced with bacterial MetH C-half sequences, suggesting that the interaction between the MTR C-half and MTRR ΔFMN is unique to eukaryotes (Supplementary Fig. 18b-d). Together, these predictions suggest that human MTRR interacts with human MTR, involving not only the FMN domain of MTRR, but also its C-terminal Conn, FAD, and NADPH domains.

**Fig. 5.**
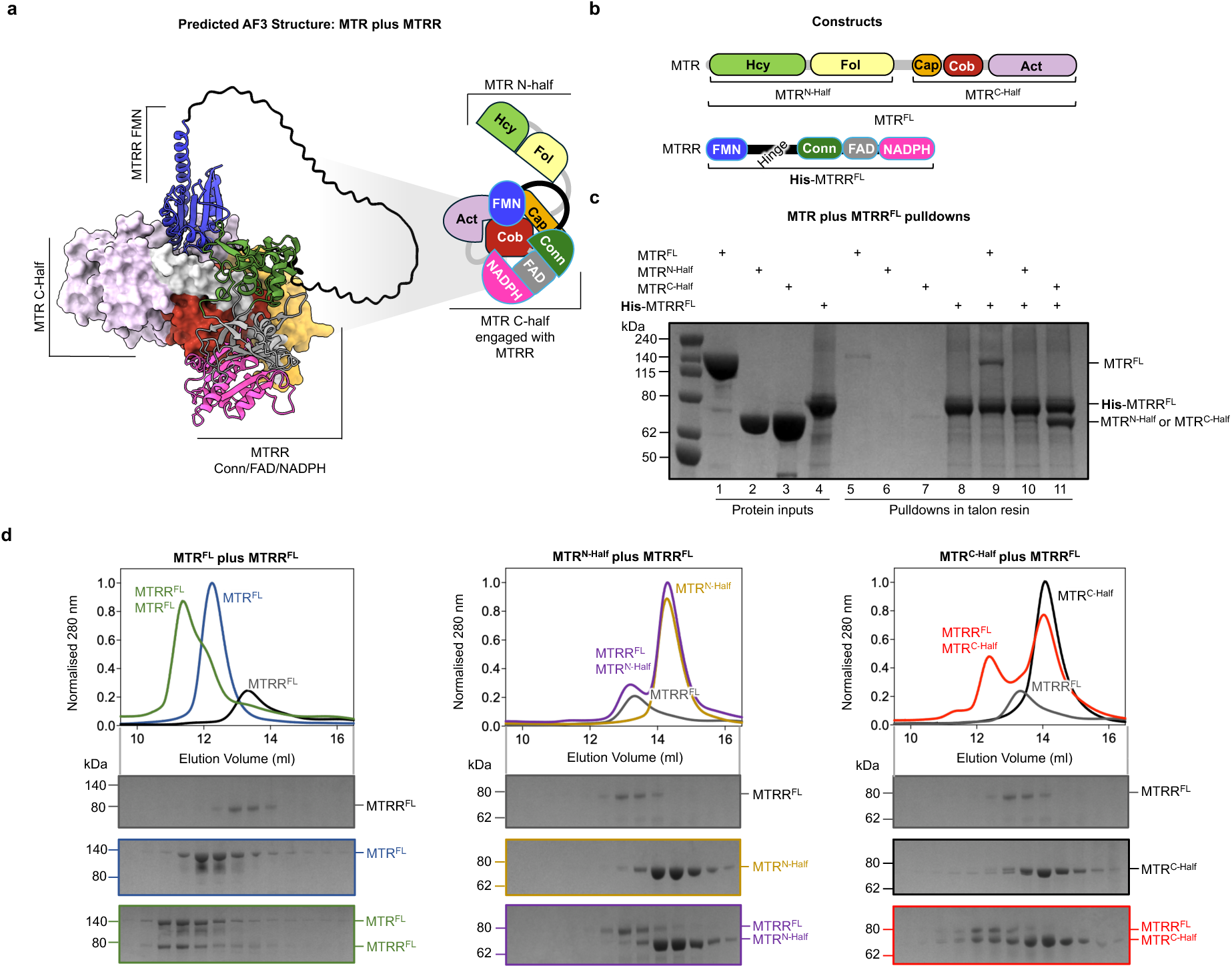
MTRR forms a stable complex with MTR. **a**, Constructs and domain arrangements of MTR and MTRR used in pull-down and gel filtration assays. Affinity tagged constructs are indicated. **b**, Representative predicted AF3 structure of the MTR-MTRR complex. The arrangement of the complex is shown as a cartoon depicted with the domain colours as shown in Fig. 5a. **c**, Talon resin affinity pulldown of the constructs shown in Fig. 5b. Lanes 1–4 represent the protein inputs used for the assay. Lanes 5–8 represent the elution fractions of each individual protein incubated with the resin to verify their ability to bind the resin. Lanes 9–11 represent the pulldown of the different protein combinations (His-MTRR^FL^ with MTR^FL^, MTR^N-Half^, or MTR^C-Half^, respectively). **d**, Analytical size-exclusion chromatography of MTRR^FL^ in complex with MTR^FL^ (left), MTR^N-half^ (middle), or MTR^C-half^ (right). Top panels show the normalised spectrum at 280 nm and bottom panels show the corresponding SDS-PAGE of the eluted fractions.

To confirm these interactions *in vitro*, we used a combination of recombinant co-expression and reconstitution pulldown experiments. Firstly, co-expressing MTR^FL^ with MTRR^FL^ in insect cells and purifying using the FLAG tag on MTR confirmed the presence of a complex without Cbl (Supplementary Fig. 19a). Additionally, in blue native PAGE experiments MTR^FL^ co-expressed with MTRR^FL^ ran at a larger molecular weight than purified MTR^FL^ alone showing complex formation (Supplementary Fig. 19b). We then employed talon resin-based affinity pulldowns using purified MTRR^FL^ as bait against the purified MTR^FL^, MTR^N-Half^, and MTR^C-Half^ (Fig. 5b, Supplementary Fig. 19c), where MTRR^FL^ showed an interaction with MTR^FL^ (Fig. 5c lane 9) and MTR^C-Half^ (lane 11), but not MTR^N-Half^ (lane 10). Importantly, these pulldown experiments confirm that the interaction between MTR and MTRR occurs through MTR^C-Half^, even in the absence of Cbl or other ligands. Analytical size exclusion assays further confirmed that both full-length MTR and MTRR can form a complex (Fig. 5d left) specifically mediated by the C-half of MTR (right), but not its N-half (middle).

To confirm the putative interaction between MTRR ΔFMN and MTR, we expressed and purified a GST tagged construct of MTRR^ΔFMN^ (Fig. 6b, Supplementary Fig. 19d) for interaction studies. In GST pulldown experiments this construct showed an interaction with both MTR^FL^ (Fig. 6c lane 9) and MTR^C-Half^ (lane 11), but not MTR^N-Half^ (lane 10). Furthermore, analytical size exclusion using MTRR^ΔFMN^ without the GST tag confirmed the interactions of this region of MTRR with MTR^FL^ (Fig. 6d left) and MTR^C-Half^ (right), but not with MTR^N-Half^ (middle). Overall, our co-expression, pulldown, and gel filtration experiments are consistent with the AF3 predictions of the complex between MTR and MTRR.

**Fig. 6.**
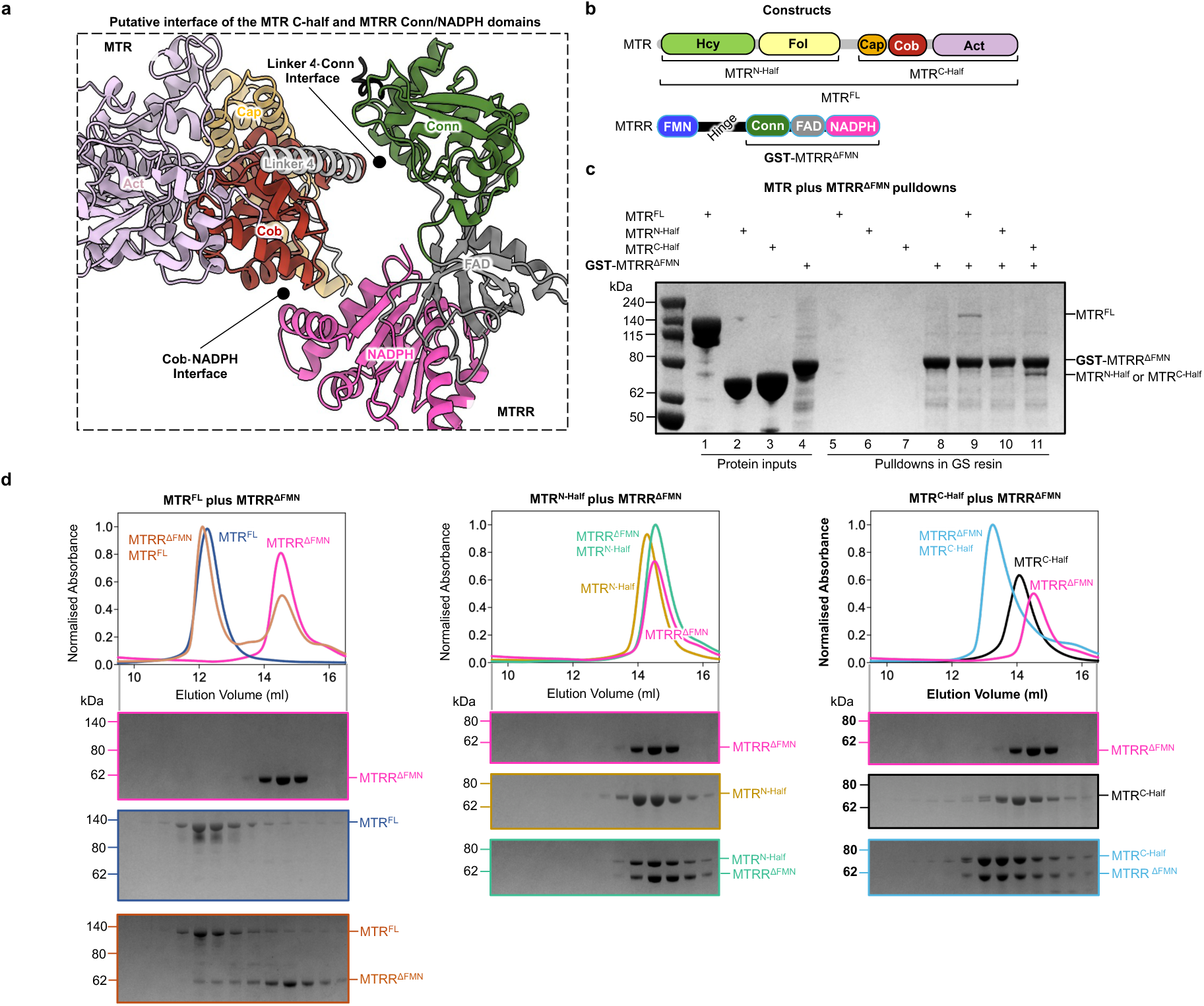
MTRR without the FMN domain can interact with MTR. **a**, Constructs and domain arrangements of MTR and MTRR used in pull-down and gel filtration assays. Affinity tagged constructs are indicated. **b**, Representative predicted AF3 structure of the MTR^C-half^-MTRR^ΔFMN^ complex. The arrangement of the complex is shown as a cartoon depicted with the domain colours as shown in Fig. 6a. **c**, GS resin affinity pulldown of the constructs shown in Fig. 6b. Lanes 1–4 represent the protein inputs used for the assay. Lanes 5–8 represent the elution fractions of each individual protein incubated with the resin to verify their ability to bind the resin. Lanes 9–11 represent the pulldown of the different protein combinations (GST-MTRR^ΔFMN^ with MTR^FL^, MTR^N-Half^, or MTR^C-Half^, respectively). **d**, Analytical size-exclusion chromatography of MTRR^ΔFMN^ in complex with MTR^FL^ (left), MTR^N-half^ (middle), or MTR^C-half^. Top panels show the normalised spectrum at 280 nm and bottom panels the corresponding SDS-PAGE analyses of the fractions indicated.

## Discussion

Utilising vitamin B12 as a cofactor, MTR coordinates folate and methionine metabolism by performing three methylation reactions through its catalytic and reactivation cycles. Early x-ray crystallography and biochemical studies^9–11,39,40^ on the bacterial homologue MetH demonstrated that the enzyme has structural modularity in order to carry out these reactions. Similarly, our cryo-EM data reveals human MTR to be modular comprising either two or three independent modules linked by flexible linkers, representing inactive and active states respectively. Studies on MetH have suggested that when the enzyme is active, large conformational changes occur as the Cob domain shuttles between the Fol and Hcy domains to transfer a methyl group from MeTHF to Hcy using MeCbl during catalysis^9^. Every ∼2000 reactions the bound cob(I)alamin intermediate can become oxidised to inactive cob(II)alamin^32^. Reactivation requires another large conformational change that repositions the Act domain to interact with the Cob domain to allow methylation of the inactive Cbl by SAM^7^. These conformational changes are highly likely to be regulated and to be conserved in human MTR. Although Cbl oxidation state and substrates have been suggested to be important contributors to conformational changes^40^, little is known on how and when they are triggered.

Recent MetH structures together with our findings on human MTR are beginning to suggest a model for how these domain movements are regulated (Figure 7). Though revealing different conformational states, as will be discussed below, our structures are consistent with recent reports of a tetra-domain structure of *T. filiformis* MetH^27^, along with a full-length structure^24^ and multiple crystal structures of different constructs of *T. thermophilus* MetH bound to various ligands^33^ (Supplementary Fig. 20). The collection of thermophilic MetH structures has revealed long sought snapshots of the full-length enzyme, and of the Cob domain engaged with either the Hcy or Fol domains, giving insights into catalysis^33^.

**Fig. 7.**
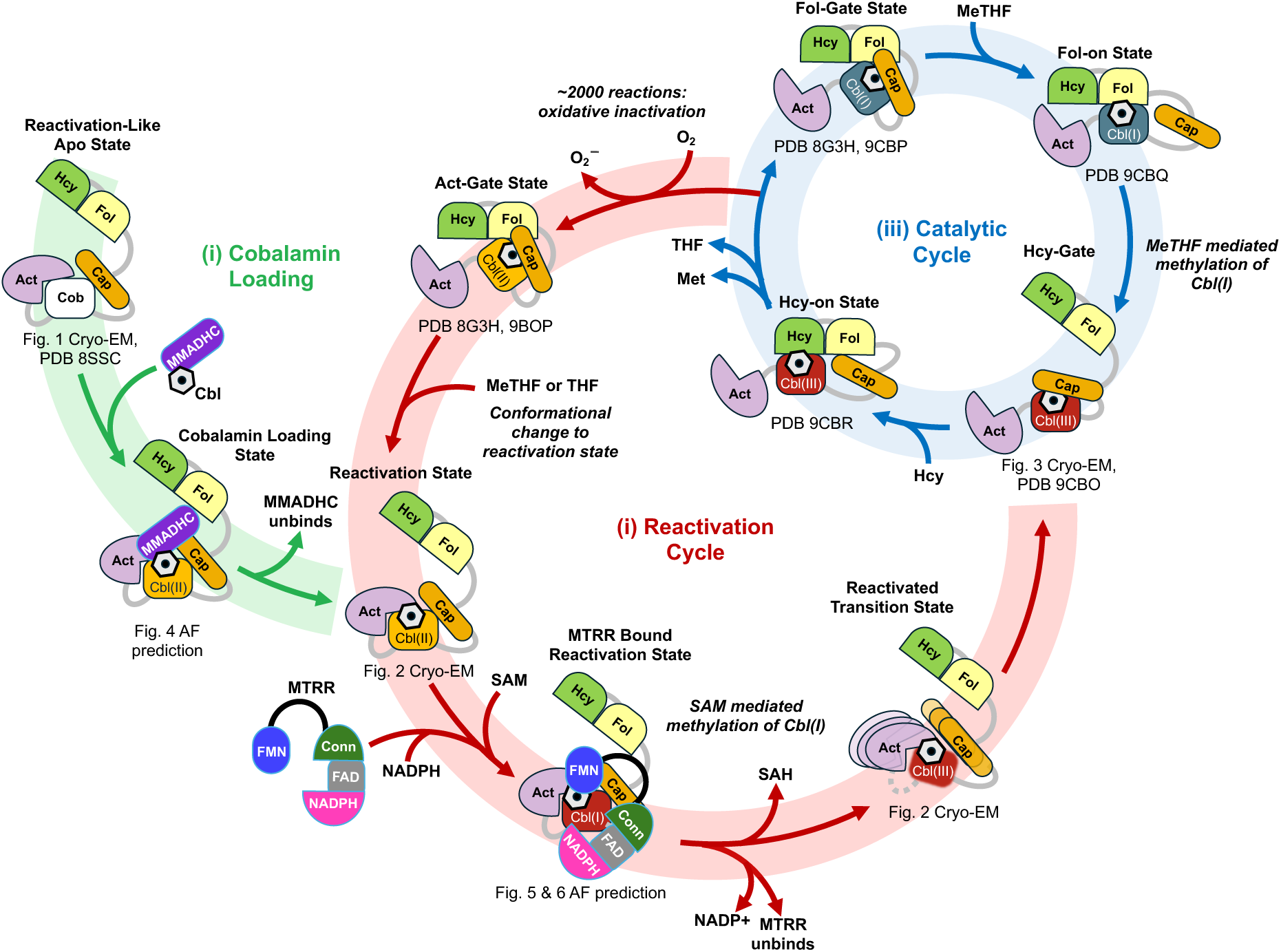
Proposed mechanism of cobalamin loading, reactivation, and catalysis of MTR. Model showing various cartoons of proposed conformations and complexes for human MTR with MMADHC and MTRR to illustrate three major aspects of its function: (i) cobalamin loading, (ii) reactivation cycle, and (iii) catalytic cycle. Please refer to the main text for more details for each aspect. Different Cbl oxidation states are stated and represented as light orange for Cbl(II) (cob(II)alamin), red for Cbl(III) (cob(III)alamin), and dark cyan for Cbl(I) (cob(I)alamin). Domains are coloured as previously depicted in previous figures. This proposed mechanism is not comprehensive. Therefore, other conformations and complexes may also occur.

Surprisingly a new conformation in MetH was initially reported by cryo-EM termed the “resting state” - a hydroxycob(II)alamin bound, substrate free conformation where the Act domain is disengaged, the Cap-on Cob module intimately interacts with the N-half, and a folate sensing loop is positioned into the folate binding site (PDB 8G3H)^27^. This resting state was replicated by crystallisation with a MetH construct without the Act domain and bound to propylcob(III)alamin (PDB 9CBP). Two additional crystal structures of a MetH Fol-Cap-Cob construct bound to MeCbl in a Cap-on conformation were also determined (PDB 9CBO)^33^. These three states were subsequently termed “gating states”, as they are interpreted to be the “Hcy-gate” (PDB 9CBP), “Fol-gate” (PDB 9CBO), and “Act-gate” (PDB 9CBO) before the requisite conformational change that leads to the Hcy-on, Fol-on, and Act-on states respectively^33^. However, both “resting state” and “Hcy-gate” structures are of inactive Cbl forms without any bound substrates, suggesting that they are more likely representative of the enzyme inactivated by oxidation and of the cob(I)alamin state before the binding of MeTHF. Therefore the resting state is more likely the shared conformation between the catalytic and reactivation cycles^27^ for both the “Act-gate” (i.e. before reactivation) and “Fol-gate” (i.e. before catalysis), depending on the oxidation state of Cbl.

Favouring this alternative interpretation is that *E. coli* MetH shows ordered ping-pong kinetics where MeTHF is the first substrate to bind and THF is the last product to leave^41^, meaning that the resting state/Hcy-gate structures could not be substrate bound intermediates during catalysis. Additionally, it is likely that the probability of oxidation of the bound cob(I)alamin is increased when MeTHF is limited, and a recent study showed that the rate of reactivation for human MTR is increased in the presence of both the substrate MeTHF and the product THF^42^. This increased reactivation likely occurs due to substrate MeTHF binding to the Fol domain and displacing the folate sensing loop^27^ allowing the formation of a Cap-off Act-on state productive for reactivation. A similar conformational change due to MeTHF binding to the cob(I)alamin state would occur resulting in a Fol-on conformation. In the MeCbl-bound “Fol-gate” and “Act-gate” crystal structures of MetH, the folate sensing loop is not displaced into the folate binding site suggesting that folate binding has little to no influence in any structural transitions in these states^33^. These two structures also have different relative orientations of the Fol domain to the Cap-on Cob domain module that is suggestive of flexibility (Supplementary Fig. 20). These structures also agree with our MeCbl-bound cryo-EM data of human MTR, where the folate sensing loop has not moved into the folate binding site, and the Act domain is disengaged from the highly flexible Cap-on Cob domain module. This conformation seen in our MTR structure, along with the MetH crystal structures, should be more aptly termed the “Hcy-gate” as the enzyme is bound to MeCbl, a Cbl form that is productive for methyl transfer to the substrate Hcy. The MeCbl-bound enzyme without bound folate adopts this more flexible state, instead of the more compact “resting state”, likely due to the distance that the Cob domain has to travel between the Fol and Hcy domains (∼50 Å) during the catalytic cycle.

Despite sharing a catalytic mechanism with MetH that exploits conformational dynamics between domains, human MTR has built in distinctive features, not present in MetH, to accommodate human Cbl metabolism. For example, while both MetH and MTR share similar reactivation conformations prior to receiving Cbl, MetH does not require a highly specialised chaperone to load the cofactor^43^. On the contrary, MMADHC has a central role in Cbl trafficking in eukaryotes and is known to direct the co-factor to either methylmalonyl-CoA mutase (MMUT) in the mitochondria or MTR in the cytosol^14^. AlphaFold predictions suggest that MMADHC engages the exposed Cbl-binding pocket via all three domains in the C-half of MTR when in a reactivation conformation. This interaction model aligns with experimental data on Cbl loading, suggesting a structural rationale for how MMADHC and MTR form a complex. Cbl is loaded as cob(I)alamin onto MTR, but most of it is readily oxidised into cob(II)alamin^17^ which cannot support catalysis. The reductive methylation of cob(II)alamin, required for the reactivation, is assisted by electron transfer from the essential flavoprotein MTRR^20^. MTRR is an unusual member of the diflavin reductase family where it has a much longer hinge region linking the N-terminal FMN domain with the rest of the enzyme, than other family members^44^. Previous studies revealed that the FMN domain of MTRR engages with MTR to reduce the cobalamin, which is subsequently methylated to MeCbl^19^. Our AF3 modelling and biophysical analyses of the MTR-MTRR complex revealed previous unknown interacting interfaces between these proteins and explains its long hinge region. AF3 predicted models suggested that MTRR can also interact with MTR through its NADPH domain and Conn domain when the FMN domain is engaged. This unexpected additional interaction was confirmed by our biophysical assays using truncated constructs. The additional interfaces could indicate a mechanism of MTR to optimize the efficiency of the reductive methylation by forming a stable complex with MTRR even prior to entering the reactivation cycle. Expectedly, this interaction is not predicted between MTRR and MetH, the latter relying on a different reactivation system supported by flavodoxin^21^. The fact that MTR cannot be reactivated by flavodoxin, and conversely MetH cannot be reactivated by MTRR, suggests that this newly identified interaction may be essential for Cbl reduction^25^. This lack of cross-reactivity in reactivation could be due to the lower sequence conservation in the region of MTR that is responsible for forming this extended interaction with MTRR (Supplementary Fig. 21 α-helix 35).

Figure 7 integrates our cryo-EM snapshots, AF3 predictions, biophysical data, and interpretation of previous studies along with other structures, into a proposed composite mechanism for Cbl loading, reactivation, and catalysis. During cobalamin loading (i), apo MTR is poised in the Cap-off reactivation conformation, ready to interact with MMADHC for receiving Cbl. MMADHC loads cob(I)alamin which without SAM can be readily oxidised to cob(II)alamin. Therefore, during the reactivation cycle (ii) MTR still in the reactivation conformation is ready to interact with MTRR, for the reductive methylation of the bound Cbl using SAM. This interaction is secured by two interfaces and allows an efficient electron transfer from the FMN domain of MTRR to the Cbl. Following Cbl methylation, MTR undergoes a large conformational change as the bound Cbl goes from His-off to His-on, triggering MTRR unbinding. This results in the flexible MeCbl-bound state (Hcy-gate), where the Act and Cob domains disengage, and the Cap domain takes its position atop the Cbl to protect it from oxidation before beginning catalysis. This primes MTR within the catalytic cycle (iii) ready to engage with the Hcy domain to carry out a methyl transfer reaction to form Met. After catalysis, the products THF and Hcy leave MTR and result in a “Fol-gate” Cap-on conformation to protect the cob(I)alamin intermediate. If oxidation has occurred that results in cob(II)alamin, MTR would also form the same Cap-on conformation. In these two scenarios, binding of the substrate MeTHF would elicit different responses. When MTR is bound with cob(I)alamin, MeTHF binding to the Fol domain disrupts its interaction with the Cap and Cob domains to promote a “Fol-on” conformation to produce MeCbl and continue another catalytic cycle (iii). When MTR is bound with cob(II)alamin, MeTHF binding to the Fol domain also disrupts its interaction with the Cap and Cob domains but promotes the Act-on reactivation conformation which allows MTR to proceed with the reactivation cycle (ii).

While our proposed mechanism attempts to show how MTR traverses from a Cbl-free apo state to a MeCbl-bound active state, there are a number of caveats and unanswered questions. While our cryo-EM structures show for the first time Cbl binding of human MTR and its intrinsic flexibility, these structures are determined from samples treated with free Cbl in solution and thus may not be fully representative of when Cbl is loaded by MMADHC and reactivated by MTRR. Additionally, we have carried out two-way interaction studies between MTR and its proposed interacting partners without presence of any ligands or additional proteins, which may further modulate how these proteins interact. For example, MMACHC may stably interact with MTR when MMADHC is present as a ternary complex^14,15^ with possible consequences for cofactor loading. The findings that both MMADHC and MTRR can bind to MTR without any bound Cbl also suggests that these proteins may have further roles beyond Cbl loading and reactivation. Supporting this hypothesis is a previous report which suggested that MTRR could act as a chaperone of apo MTR as it increases MTR protein stability^25^. A recent study also proposed a chaperone role of MMADHC for MTR^42^. These studies point to the possibility of ternary complexes where a Cbl-free complex of MTR and MTRR may interact with Cbl-free or Cbl bound MMADHC. This is also underlined by reports of a quaternary complex of all four proteins namely MTR, MTRR, MMACHC and MMADHC^12^. Another report suggested a direct interaction of MMACHC with MTR^35^. However, this does not appear to be supported by our AF3 predictions and pulldown assays where MMACHC shows little to no interaction. Clearly, more studies delving into three/four-way complexes are warranted in future.

Overall, our findings expand current knowledge on human MTR and contextualize previous reported findings on both human and bacterial orthologues. Importantly we reveal insights into eukaryote-unique aspects in terms of the complexes that MTR forms to support its function. Given that pathogenic mutations in the MTR, MTRR, MMACHC, and MMADHC genes are linked to Cbl deficiency, it is likely that some variants effect residues at protein-protein interfaces of these MTR complexes. Indeed, disease-causing variants of MMADHC have been suggested to weaken its interaction with MTR hindering Cbl loading^17^. Consequently, future experimental structure determination of the MTR complexes with MMADHC and MTRR will be essential in understanding how pathogenic mutants alter their complex formation. Such molecular insights could inform the development of targeted therapies towards the treatment of these rare metabolic disorders.

## Methods

### Cloning, expression, and purification of MTR constructs

Full-length MTR (MTR^FL^) for cryo-EM, activity, and thermal stability assays was cloned into pFB-CT10HF-LIC encoding for a C-terminal TEV cleavable His-tag and FLAG-tag for insect cell expression. MTR^FL^ was expressed in Sf9 cells infected with P3 baculovirus and grown in Sf-900 III SFM (Life Technologies) for 72 hrs. Cell pellets were harvested and homogenized in lysis buffer (50 mM HEPES pH 7.5, 500 mM NaCl, 5% glycerol, 0.5 mM Tris(2-carboxyethyl)phosphine hydrochloride (TCEP) and 20 mM imidazole) using sonication, and insoluble material was removed by centrifugation. MTR was then purified by affinity (Ni-sepharose; Cytiva) and size-exclusion (Superdex 200; Cytiva) chromatography to separate monomeric MTR from aggregates. Purified MTR^FL^ was then concentrated to 3–10 mg/ml, aliquoted, and stored in storage buffer (50 mM HEPES pH 7.5, 500 mM NaCl, 5% glycerol, 0.5 mM TCEP) at −80 °C.

Full-length MTR (MTR^FL^) for pull-down and gel filtration assays, was cloned into pNIC-CT10HSII encoding for a C-terminal TEV cleavable His-tag and tandem StrepII-tag for bacterial expression. MTR^FL^ was expressed in ArcticExpress DE3 cells. Cells were cultured in Terrific Broth (Formedium) at 30 °C in Terrific Broth and expression was carried out for 60 hrs at 10 °C induced with 0.1 mM isopropyl-β-D-thiogalactopyranoside (IPTG). Cell pellets were harvested and homogenized in lysis buffer (50 mM sodium phosphate pH 7.5, 500 mM NaCl, 5% glycerol, 0.5 mM TCEP, 1.0 % triton X-100, and 20 mM imidazole) using sonication, and insoluble material was removed by centrifugation. MTR^FL^ was then purified from the supernatant by tandem affinity using Ni-sepharose followed by Streptactin XT resin (Cytiva). MTR^FL^ was then treated with His-tagged TEV protease overnight at 4 °C and passed over Ni-sepharose resin to remove the TEV protease and any uncleaved protein. A final size-exclusion (Superdex 200; Cytiva) chromatography step was used to separate monomeric MTR^FL^ from degradation products. Purified MTR^FL^ was then concentrated to 3–10 mg/ml, aliquoted, and stored in storage buffer (50 mM HEPES pH 7.5, 500 mM NaCl, 5% glycerol, 0.5 mM TCEP) at−80 °C.

The N-terminal construct of MTR, residues 16-657 (MTR^N-Half^), was cloned into pNIC-Bsa4 encoding for a N-terminal TEV cleavable His-tag. The C-terminal construct of MTR, residues 661-1265, (MTR^C-Half^) was cloned and into pNIC-CH as a TEV cleavable His-GST N-terminally tagged for bacterial expression (Twist Bioscience). A stop codon was added at the end of the MTR^C-Half^ coding sequence before the vector encoded His-tag. Both constructs were expressed in Rosetta DE3 cells. Cells were cultured in Terrific Broth (Formedium) at 37 °C and expression was induced overnight at 18 °C with 0.2 mM IPTG. Pellets were harvested, homogenized in lysis buffer (50 mM HEPES pH 7.5, 500 mM NaCl, 5% glycerol, 0.5 mM TCEP and 20 mM imidazole) by sonication, and insoluble material was removed by centrifugation. The supernatant was subjected to affinity (Ni-sepharose; Cytiva) and the eluted protein applied to size-exclusion (Superdex 200; Cytiva) chromatography. Each construct was then treated with His-tagged TEV protease overnight at 4 °C and passed over Ni-sepharose resin to remove the TEV protease and uncleaved protein. In some incidences, an additional size-exclusion step was carried out to remove residual TEV protease. Purified MTR^N-Half^ and MTR^C-Half^ proteins were concentrated to 10–20 mg/ml and stored in storage buffer (50 mM HEPES pH 7.5, 500 mM NaCl, 5% glycerol, 0.5 mM TCEP) at −80 °C.

### Cloning, expression, and purification of MTRR constructs

Full-length codon optimised MTRR (MTRR^FL^) was cloned into pFastBac (Genscript) encoding for a TEV cleavable N-terminal His-tag and was expressed in Sf9 cells grown in Sf-900 III SFM (Life Technologies) for 72 hrs. Cell pellets were harvested and resuspended in lysis buffer (50 mM sodium phosphate pH 7.5, 500 mM NaCl, 5% glycerol, 0.5 mM TCEP, 1.0% triton x-100, 100 μM flavin mononucleotide). The resuspended pellet was incubated at room temperature for 30 minutes with spinning, followed by sonication, and then subsequent centrifugation to remove insoluble material. MTRR^FL^ was then purified from the soluble fraction by affinity (Ni-sepharose; Cytiva). Purified MTRR^FL^ was then diluted in low salt buffer (50 mM HEPES pH 7.5, 50 mM NaCl, 5% glycerol, 0.5 mM TCEP) and applied to a HiTrap Q column (Cytiva). A gradient from 50 mM to 1M NaCl was used to elute MTRR^FL^. Appropriate fractions were pooled, concentrated, and buffer exchanged into storage buffer (50 mM HEPES pH 7.5, 500 mM NaCl, 5% glycerol, 0.5 mM TCEP.)

The MTRR construct without the FMN domain, residues 217-698, (MTRR^ΔFMN^) was cloned from a codon optimised sequence (Twist Bioscience) into pGTVL2 to produce a TEV cleavable His-GST N-terminally tagged protein for bacterial expression. MTRR^ΔFMN^ was expressed in Rosetta DE3 cells. Cells were cultured in Terrific Broth (Formedium) at 37 °C and expression was induced overnight at 18 °C with 0.2 mM IPTG. Pellets were harvested, homogenized in lysis buffer (50 mM sodium phosphate pH 7.5, 500 mM NaCl, 5% glycerol, 0.5 mM TCEP, 1.0% triton X-100, and 20 mM imidazole) by sonication, and insoluble material was removed by centrifugation. To produce tagged protein for pull-down assays the supernatant was subjected to affinity (Ni-sepharose; Cytiva) and the eluted protein was further purified by anion exchange using a HiTrap Q column (Cytiva). Clean fractions containing the GST tagged construct were pooled and further treated to size-exclusion (Superdex 200; Cytiva) chromatography. To produce cleaved MTRR^ΔFMN^ protein for analytical gel filtration assays the supernatant was subjected to affinity (Ni-sepharose; Cytiva), and the eluted protein incubated with His-tagged TEV protease overnight at 4 °C. The cleaved protein was then subjected to size-exclusion (Superdex 200; Cytiva) chromatography. Peak fractions of cleaved MTRR^ΔFMN^ were pooled and passed over Ni-sepharose resin to remove any remaining TEV protease. Both the tagged and cleaved MTRR^ΔFMN^ proteins were concentrated to 5-10 mg/ml and stored in storage buffer (50 mM HEPES pH 7.5, 500 mM NaCl, 5% glycerol, 0.5 mM TCEP) at −80 °C.

### Co-expression, and purification of the MTR-MTRR complex

Using the biGBac cloning system the genes for full-length MTR and full-length MTRR were cloned into pBIG1a. The MTR^FL^ gene encoded for a C-terminal TEV cleavable His and FLAG whereas the MTRRFL gene encoded a N-terminal His-tag. The MTR^FL^-MTRR^FL^ complex was expressed in Sf9 cells grown in Sf-900 III SFM (Life Technologies) for 72 hrs. Cell pellets were harvested and homogenized in lysis buffer (50 mM HEPES pH 7.5, 500 mM NaCl, 5% glycerol, 0.5 mM TCEP) using sonication, and insoluble material was removed by centrifugation. The complex was then purified by affinity (Ni-sepharose; GE Healthcare) and size-exclusion (Superdex 200; GE Healthcare) chromatography. Fractions corresponding to the complex were pooled, concentrated to 5–10 mg/ml and stored in storage buffer (50 mM HEPES pH 7.5, 500 mM NaCl, 5% glycerol, 0.5 mM TCEP) at −80 °C.

### Cloning, expression, and purification of MMACHC and MMADHC**^ΔN^**

The cloning, expression, and purification of MMACHC and MMADHC^ΔN^ (residues 124-296) has been described in detail elsewhere^13,14^. In brief His-GST tagged MMACHC and MMADHC^ΔN^ were expressed in Rosetta DE3 cells. Cells were cultured in Terrific Broth (Formedium) at 37 °C and expression was induced overnight at 18 °C with 0.1 mM IPTG. Pellets were harvested, homogenized in lysis buffer (50 mM HEPES pH 7.5, 500 mM NaCl, 5% glycerol, 0.5 mM TCEP and 20 mM imidazole) by sonication, and insoluble material was removed by centrifugation. The supernatant was subjected affinity (Ni-sepharose; Cytiva) and the eluted protein applied to size-exclusion (Superdex 200; Cytiva) chromatography. To produce cleaved MMADHC^ΔN^ the His-GST tagged protein after size exclusion was treated with His-tagged TEV protease overnight at 4 °C and passed over Ni-sepharose resin to remove the TEV protease and uncleaved protein. Purified tagged MMACHC, tagged MMADHC^ΔN^, and cleaved MMADHC^ΔN^ proteins were concentrated to 10–20 mg/ml and stored in storage buffer (50 mM HEPES pH 7.5, 500 mM NaCl, 5% glycerol, 0.5 mM TCEP) at −80 °C.

### Native mass spectrometry

By dialysis, purified MTR^FL^ was exchanged from into 50 mM ammonium acetate, pH 6.5. The sample was loaded into Quantitative Time-of-Flight (Q-TOF) 6530 mass spectrometer attached to a standard electrospray ionisation source (Agilent Technologies), with 77 a constant flow rate of 6 μl min^-1^, using a 1.0-ml gas-tight positive displacement syringe (Hamilton) and PEEK capillary tubing with an inner diameter of 0.005 inches.

### Enzyme activity assay

MTR activity assay was adapted from the previously described cuvette-based nonradioactive assay^26^. The current assay was miniaturised for a U-bottom 96-well plate. The following reagents were added to each well in the mentioned order: 24.4 µL of 100 mM phosphate buffered saline (PBS) pH 7.5, 2 µL of 500 mM dithiothreitol (DTT), 2 µL of 3.8 mM S-adenosylmethionine (SAM), 2 µL of 100 mM L-homocysteine (Hcy), 5 µL of 4 µM MTRF^L^, and 4 µL of 500 µM HOCbl. The plate was then incubated for 5 minutes at 37 °C, allowing DTT and HOCbl to create a semi-anaerobic environment by reducing O_2_ into H_2_O_2_. After this time, 2.4 µL of 6.6 mM 5-methyltetrahydrofolate (MeTHF) was added to initiate the catalytic cycle of MTR, thus producing tetrahydrofolate. After 10 minutes at 37 °C, the reaction was quenched by adding 10 µL of 5 M HCl in aqueous 60% formic acid and immediately heated at 80 °C for 10 minutes to convert the produced tetrahydrofolate to methenyltetrahydrofolate, which concentration can be determined using Δε350 = 26,500 M⁻¹ cm⁻¹. Absorbance was recorded on a OmegaSTAR (BMG Biotech), which were pathlength corrected. The data was analysed and plotted on GraphPad Prism 10.

### Thermal shift assay

MTR proteins were diluted in thermal shift buffer (25 mM HEPES, pH 7.5, 200 mM NaCl) to 0.4 mg/ml with SYPRO-Orange (Invitrogen) diluted 1000X in a total volume of 20 μl. When appropriate, ligands were added at a final concentration of 1 mM, except for HOCbl, CNCbl, and MeCbl which were added at a final concentration of 50 µM. A QuantStudio 3 RT-PCR machine (Thermo Fisher Scientific) was used to monitor protein melting across 25 °C – 95 °C with a heating rate of 0.5 °C per min. Data were analysed in GraphPad Prism software, and the thermal melting temperatures (T*_m_*) were determined using a nonlinear Boltzmann equation.

### Negative stain electron microscopy

Purified recombinant monomeric MTR^FL^ from size exclusion chromatography was diluted to a final concentration of 40 nM in buffer (25 mM HEPES pH 7.5, 250 mM NaCl). 10 µl sample was added to copper grids with carbon coating (300 mesh, Quantifoil Micro Tools GmbH), plasma treated for 1 min using PELCO easiGlow™ (Ted Pella, Inc). After 30 seconds excess sample was dried off using filter paper (Whatman), and the grid was placed on a drop of filtered 1.5 % uranyl acetate (Sigma-Aldrich) for 30 seconds. Grids were dried again using filter paper before visualisation. Imaging was performed using a Hitachi HT7800 microscope operating at 120 kV and with a magnification of 60k, located at the Electron Microscopy Research Services (EMRS) facility at Newcastle University.

### Grid preparation and cryo-electron microscopy

MTR^FL^ was diluted to 0.5 mg/ml into 25 mM HEPES, pH 7.5, 250 mM NaCl with either 1 mM MeTHF, 1 mM MeTHF, 1 mM SAM and 50 μM HOCbl, or 50 μM MeCbl. Grids were prepared using a FEI Vitrobot Mark III (Thermo Fisher Scientific) at 4 °C and 100% humidity. 3 µl of sample was applied to a plasma treated R1.2/1.3 300 mesh UltrAuFoil (Quantifoil) grid, with a blot force of 0, a blot time of 3 seconds and a wait time of 10 seconds. All grids were initially screened by collecting a small ≈ 1,000 micrograph dataset on a 200 kV Glacios microscope equipped with a Falcon 4 direct electron detector (ThermoFisher Scientific) at York Structural Biology Laboratory. Data was collected at a magnification of 240k with a pixel size of 0.574 Å and a total dose of 50 e^-^/Å^2^. Glacios data was processed in cryoSPARC^45^ and only grids that resulted in clear 2D classes were sent for larger data collections on a 300 keV Krios microscope.

All high-resolution data collection was carried out at Electron Bio-Imaging Centre (eBIC, Diamond Light Source, UK). For MTR^FL^ with MeTHF, and MTR^FL^ with MeTHF, SAM and HOCbl were collected on a 300 kV Titan Krios equipped with a K3 direct electron detector and energy filter (Gatan). K3 data was collected at a magnification of 165k equating to a pixel size of 0.508 Å (super resolution mode 0.254 Å) with an exposure time of 1.00 s, and a total dose of 65.20 e^-^/Å^2^. For MTR^FL^ with MeCbl data was collected on a 300 kV Titan Krios equipped with a Falcon 4i direct electron detector and selectris X energy filter (ThermoFisher Scientific). Falcon 4i data was collected at a magnification of 215k equating to a pixel size of 0.576 Å with an exposure time of 2.46 s and a total dose of 70 e^-^/Å^2^.

Data processing for each dataset followed the same following general process: All movies were imported into cryoSPARC^45^ where they were subjected to patch motion correction and patch CTF estimation. Micrographs were into 2,000 micrograph subsets for particle picking, extraction, and 2D classification. All 2D classifications used the standard preset parameters expect that force over max poses/shifts was turned off, the number of online-EM iterations was increased from 20 to 40, the number of full EM iterations was increased from 1 to 3, and the batch size was increased to 400. Initially blob picking was used on each micrograph subset. Particles were extracted in a 400-pixel box Fourier cropped to 200 pixels and subjected to multiple rounds of 2D classification to remove bad picks and junk. Once good 2D classes were determined it was clear that two types of particles were present in each dataset. Separate 2D classes for these particle types were used for separate template-based picking which were then again extracted in a 400-pixel box Fourier cropped to 200 pixels and subjected to rounds of 2D classification. Finally for both particle types, these particles were used in combination with Topaz^46^ to pick particles that were missed or poorly centred by blob and template picking. Again, these particles were extracted in a 400-pixel box Fourier cropped to 200 pixels and subjected to rounds of 2D classification. For each particle type particles from blob, template, and Topaz^46^ based picking were separately pooled, duplicates removed and subjected to another round of 2D classification. Ab-initio with three to six classes followed by heterogenous refinement was used to determine the best particles that were then subjected to non-uniform refinement followed by local refinement. Reference based motion correction was then used followed by another round of ab-initio, heterogenous, non-uniform, and local refinement. In the case of the MTR^C-Half^ maps 3D variability analysis^30^ was carried out with 3 components filtered to 5.0 Å. For the MTR^C-Half^ map from the MeTHF, SAM, and HOCbl dataset 3D variability was done in cluster mode followed by local refinement. One cluster equated to the His-off state whereas one cluster equated to the His-on state. As none of the maps reached Nyquist the extracted particle images were left Fourier cropped meaning that maps from the K3 datasets have a pixel size of 1.016 Å and the Falcon 4i maps have a pixel size of 1.152 Å. More details are shown in the supplementary information (Supplementary Fig. 3, 5, and 8). Local resolution, conical FSC area ratio (cFR), and sampling compensation factor (SCF) were determined within cryoSPARC^45^ for each map (Supplementary Fig. 9).

### Atomic model fitting, refinement, and validation

For all MTR structures described we used the predicted AlphaFold structure of MTR^FL^ (<u>Q99707</u>)^47^ as an initial starting model. For each map the appropriate domains were manually fitted and docked using ChimeraX^48^. Isolde^49^ global flexible fitting was then carried out followed by real space refinement as implemented in Phenix^50^. Ramachandran outliers were inspected and manually corrected in Coot^51^ followed by a final Phenix real space refinement. For the ligands methyltetrahydofolate and cobalamin, Phenix eLBOW^52^ was used to produce their restraints. Each ligand was manually docked into their appropriate density using Coot and were subjected to rounds of manual refinement and Phenix real space refinement. All models were validated using Molprobity^53^. Model fit was assessed by model to map FSC determination in Phenix^50^ and using the Q-score^54^ implementation in ChimeraX^48^ (Supplementary Fig. 10)

### Structure prediction using AlphaFold3

The AlphaFold3 server^36^ was used to predict the complex structures of MTR (<u>Q99707-1</u>), MTRR (<u>Q9UBK8-2</u>), MMACHC (<u>Q9Y4U1</u>), and MMADHC (<u>Q9H3L0</u>) using their amino acid sequences as input. The sequences of MTR from *Thermus thermophilus* (<u>Q9RA53</u>), *T. filiformis* (<u>A0A0A2XCD7</u>), and *Escherichia coli* (<u>P13009</u>) were also used for the AF3 complex predictions. In some incidences the sequences corresponding to the boundaries of MTR^N-Half^, MTR^C-Half^, and MTRR^ΔFMN^ were used. All five output models for each prediction were manually inspected in ChimeraX^48^. pLDDT^55^ values from the server were used to analyse the confidence of modelling for individual proteins. Both pTM and ipTM scores calculated from the server were used to judge the confidence of each predicted complex structure^36,56,57^.

### Blue native-PAGE

Blue native-PAGE was carried out according to manufacturer’s instructions (Life Technologies). Both MTR^FL^ and the MTR^FL^-MTRR^FL^ complex expressed and purified from insect cells were diluted to 1.0 mg/ml in 25 mM HEPES, pH 7.5, 250 mM NaCl.

### Talon pull-down assay

The bait, MTRR^FL^ with a N-terminal His-tag at 0.5 mg ml^−1^ was mixed with either prey proteins MTR^FL^, MTR^N-Half^ or MTR^C-Half^ without His-tags at 0.5 mg ml^−1^ in a total volume of 100 μl diluted in binding buffer (25 mM HEPES, pH 7.5, 100 mM NaCl, 1 mM TCEP, 0.2% Tween-20). Protein mixes were incubated with rotation at room temperature for 30 min. Next, 80 μl of a 50% slurry of talon resin (Clontech) in binding buffer was added, followed by incubation for a further 30 min at room temperature. The resin was washed with 2 ml of binding buffer with 10 mM imidazole and eluted with 40 μl 4× sodium dodecyl sulfate (SDS)–PAGE sample buffer. Controls consisted of adding prey to resin without bait. All samples were run on SDS–PAGE and stained with Coomassie blue. Each experiment was replicated two times.

### Glutathione pull-down assay

The bait proteins, MMACHC, MMADHC^ΔN^ or MTRR^ΔFMN^ each with a N-terminal GST-tag at 0.3 mg ml^−1^ were mixed with either prey proteins MTR^FL^, MTR^N-Half^ or MTR^C-Half^ without GST-tags at 0.3 mg ml^−1^ in a total volume of 100 μl diluted in binding buffer (25 mM HEPES, pH 7.5, 100 mM NaCl, 1 mM TCEP, 0.2% Tween-20). Protein mixes were incubated with rotation at room temperature for 30 min. Next, 200 μl of a 50% slurry of glutathione sepharose 4 fast flow resin (Cytiva) in binding buffer was added, followed by incubation for a further 60 min at room temperature. The resin was washed with 2 ml of binding buffer and eluted with 40 μl 4× sodium dodecyl sulfate (SDS)–PAGE sample buffer. Controls consisted of adding prey to resin without bait. All samples were run on SDS–PAGE and stained with Coomassie blue. Each experiment was replicated two times.

### Cobalamin loading assay

Cobalamin loading from MMADHC to MTR was carried out as similar to described^17^. 100 µM of HOCbl was mixed with 120 µM of MMADHC^ΔN^ in KCl Buffer (100 mM HEPES, 150 mM KCl, 10% glycerol) for a final volume of 40 µL. The reaction was incubated at room temperature for 1 hour, and the UV-Vis spectrum was recorded every 3 minutes to check the reaction. To detect the cobalamin transfer from MMADHC^ΔN^ to MTR^FL^, MMADHC^ΔN^ bound was concentrated by multiple rounds of centrifugation (4000 xg, 4 °C) in Amicon® concentrators with a 10 kDa molecular weight cut-off (Sigma Aldrich) followed by adding more KCl buffer to remove free HOCbl. 15 µM MMADHC^ΔN^-HOCbl was then incubated with 8.8 µM of MTR in a final volume of 100 µL of KCl buffer for 1 hour at room temperature. After this time, the sample was loaded into a Superose® 6 10/300 GL gel filtration column (Cytiva) equilibrated with KCl buffer at 4 °C at a flow rate of 0.2 ml/min. The elution profile was monitored at 280 and 350 nm to detect the protein and cobalamin respectively. For binding of HOCbl and MeCbl, 8.8 µM of MTR^FL^ was incubated with 100 µM Cbl for and the applied onto the gel filtration column. All experiments were performed twice except for MeCbl binding which was carried out only once.

### Analytical size exclusion chromatography

Analytical size exclusion chromatography was carried out using a 10/300 GL S200 Increase column (Cytiva) equilibrated in 25 mM HEPES, pH 7.5, 150 mM NaCl. Proteins were mixed as appropriate in 25 mM HEPES, pH 7.5, 150 mM NaCl to a final concentration of each protein at 1.5 mg/ml in a total volume of 250 ul. After 30 minutes incubation rotating at room temperature each were loaded onto the size-exclusion column and eluted with a flow rate of 0.3 ml/min. 500 µl fractions were collected for analysis by SDS-PAGE.

## Data availability

The authors declare that the main data supporting the findings of this study are available within the article and Supplementary Information. EM maps and models generated in this study of Human Methionine Synthase With Methyltetrahydrofolate, N-Half From Full-Length (EMDB-<u>55190</u>, PDB <u>9SSP</u>), Human Methionine Synthase With Methyltetrahydrofolate, C-Half From Full-Length (EMDB-<u>55191</u>, PDB <u>9SSQ</u>), Human Methionine Synthase With Methyltetrahydrofolate, Hydroxocobalamin, and SAM, N-Half From Full-Length (EMDB-<u>55192</u>, PDB <u>9SSR</u>), Human Methionine Synthase With Methyltetrahydrofolate, Hydroxocobalamin, and SAM, C-Half His-ON From Full-Length (EMDB-<u>55193</u>, PDB <u>9SSS</u>), Human Methionine Synthase With Methyltetrahydrofolate, Hydroxocobalamin, and SAM, C-Half His-OFF From Full-Length (EMDB-<u>55194</u>, PDB <u>9SST</u>), Human Methionine Synthase With Methylcobalamin, N-Half From Full-Length (EMDB-<u>55195</u>, PDB <u>9SSU</u>) and Human Methionine Synthase With Methylcobalamin, Activation Domain From Full-Length (EMDB-<u>55196</u>, PDB <u>9SSV</u>), have been deposited to the Electron Microscopy Data Bank (EMDB) and Protein Data Bank (PDB) Other structures referenced in this article are indicated, including PDB ID <u>4CCZ</u>, <u>8G3H</u>, <u>8SSC</u>, <u>9CBO</u>, <u>9CBP</u>, <u>9CBQ</u>, and <u>9CBR</u>. Source data are provided with this paper.

## Acknowledgements

Negative stain electron microscopy was done at Newcastle University EM Research Services, and we are grateful to Tracey Davey for technical support. The Newcastle University EM Research Services is funded by the BBSRC (grant number BB/R013942/1). We thank Johan Turkenburg and Sam Hart for their assistance in collecting Glacios data at York Structural Biology Laboratory (YSBL). The YSBL is funded by the BBSRC, the Wellcome Trust (grant number 206161/Z/17/Z), Tony Wild, and the Wolfson Foundation. We acknowledge the Diamond Light Source for access and support to the UK’s Electron Bio-imaging Centre (eBIC, under BAG proposal BI34172) funded by the Wellcome Trust, MRC, and BBRSC. We specifically want to thank David Owen and David Farmer for their assistance in data collecting.

D.S.M.F. was supported by the Nuffield Department of Medicine Prize Studentship at the University of Oxford. T.J.M. received cryo-EM training through the Wellcome/MRC funded programme (218785/Z/19/Z). Initial work on this project was carried out at the Structural Genomics Consortium, a registered charity (Number 1097737) that receives funds from AbbVie, Bayer Pharma AG, Boehringer Ingelheim, Canada Foundation for Innovation, Eshelman Institute for Innovation, Genome Canada, Innovative Medicines Initiative (EU/EFPIA) [ULTRA-DD grant no. 115766], Janssen, Merck & Co., Novartis Pharma AG, Ontario Ministry of Economic Development and Innovation, Pfizer, Sao Paulo Research Foundation-FAPESP, Takeda, and Wellcome Trust [092809/Z/10/Z]. W.W.Y. holds a visiting professorship at Nuffield Department of Medicine, University of Oxford.

## Author contributions

D.S.M.F., J.M.E., J.A.C., W.W.Y., and T.J.M. designed the experiments. D.S.M.F. expressed and purified MTR and MTRR constructs, carried out biochemical experiments, screened, and collected EM data, analysed, and refined all cryo-EM structures. K.M., C.D., H.J.B., and J.K. purified MTR constructs and did preliminary experiments. W.K. expressed, purified, and crystallized the N-terminal MTR construct. M.V. solved and refined the N-terminal MTR construct crystal structure. D.S.F. expressed and purified MMADHC and MMACHC. T.J.M. expressed and purified MMACHC, MTRR and MTR proteins along with guiding cryo-EM screening, data collection, data processing, AF3 predictions, and biochemical experiments.

R.C. carried out the native mass spectrometry. A.B. supplied computational resources for cryo-EM data processing. D.S.M.F., W.W.Y., and T.J.M. supervised the work, carried out data analysis, and wrote the manuscript with contributions from all authors.

## Competing interests

The authors declare no competing interests.

Correspondence and requests for materials should be addressed to Wyatt W. Yue or Thomas J. McCorvie

