## Supplementary Material for "Structural insights into cobalamin loading and reactivation of human methionine synthase"

[5]Present address: European Bioinformatics Institute (EMBL-EBI), Main Building, A2-34, Wellcome Genome Campus, Hinxton, CB10 1SD, United Kingdom

[6]Present address: Division of Metabolism and Children's Research Center, University Children's Hospital Zurich, University of Zurich, Zurich 8008, Switzerland

[7]Present address: Institute of Biochemistry II, Medical Faculty, Goethe-University, Frankfurt am Main and Buchmann Institute for Molecular Life Sciences, Frankfurt am Main, Germany

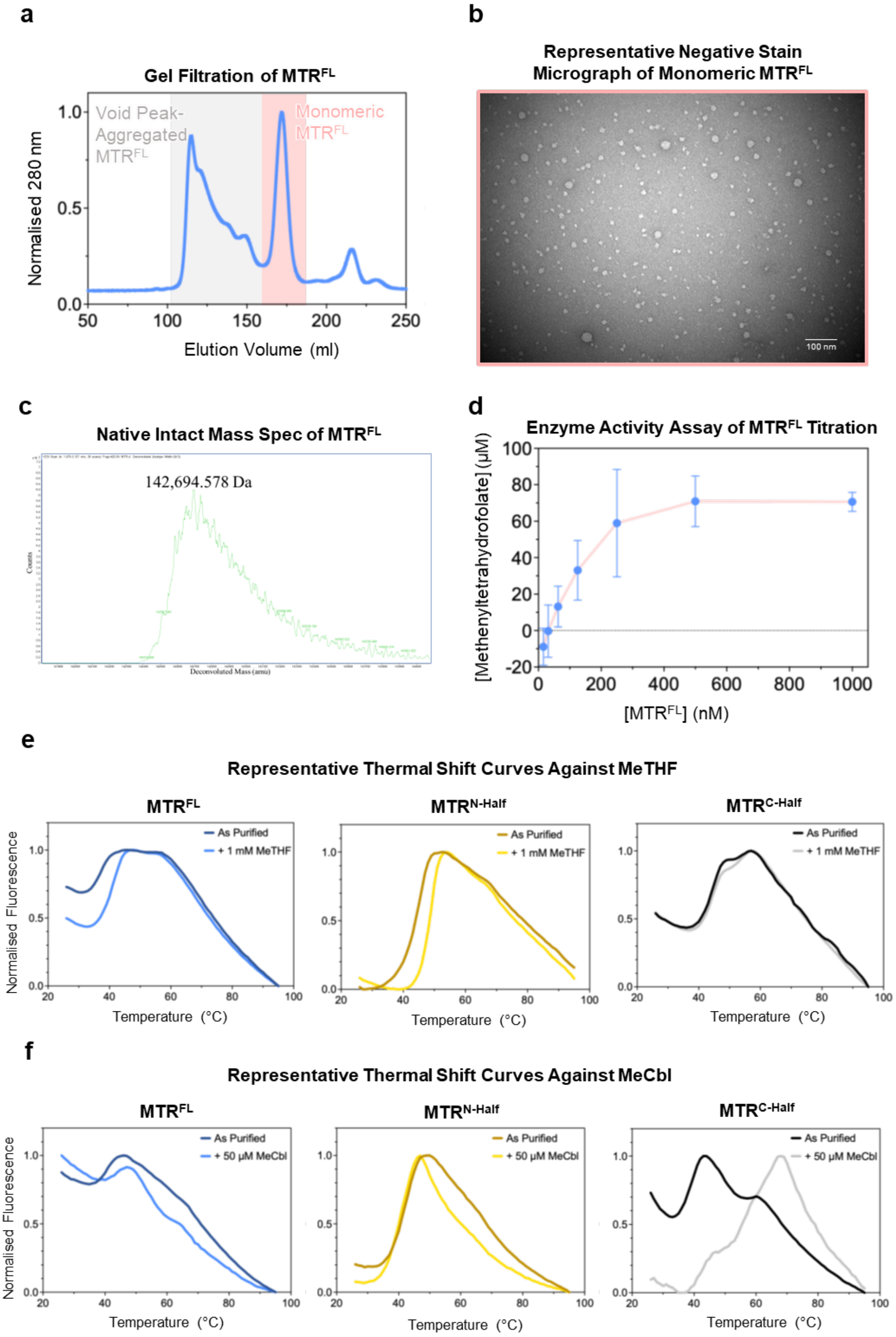


**g**

| $\Delta T_m$ (°C) | MeTHF | Homocysteine | SAM | HOCbl | CNCbl | MeCbl |
|---|---|---|---|---|---|---|
| $MTR^{FL}$ | +3.34 ± 0.31 | 0.00 ± 0.32 (n = 2) | −0.25 ± 0.23 | +0.33 ± 0.36 | +1.38 ± 0.25 | +2.8 ± 0.5 |
| $MTR^{N\text{-}Half}$ | +4.87 ± 0.51 | −0.08 ± 0.52 | −0.10 ± 0.76 | −1.67 ± 0.70 | −0.30 ± 0.46 | -1.1 ± 0.2 |
| $MTR^{C\text{-}Half}$ | +0.94 ± 0.92 | +0.84 ± 0.81 | +0.61 ± 0.46 (n = 2) | +1.58 ± 0.39 | +2.37 ± 0.62 | +19.2 ± 2.3 |

**Supplementary Fig. 1 | Characterization of MTR$^{FL}$ as purified. a,** Size-exclusion chromatography (Superdex 200, Cytivia) of Ni-NTA purified MTR$^{FL}$, showing distinctive peaks corresponding to aggregated and monomeric fractions of MTR$^{FL}$. **b,** Representative negative stain micrograph of monomeric MTR$^{FL}$ demonstrating the protein is monomeric and homogeneous. **c,** Native Mass Spectrometry of MTR$^{FL}$ confirming the absence of bound ligands in the sample. **d,** Enzymatic activity assay of MTR$^{FL}$, showing the protein activity in increasing enzyme concentrations over 10 minutes. **e,** Representative thermal shift profiles of MTR$^{FL}$, MTR$^{N\text{-}Half}$, and MTR$^{C\text{-}Half}$ in the presence of MeTHF. **f,** Representative thermal shift curves of MTR$^{FL}$, MTR$^{N\text{-}Half}$, and MTR$^{C\text{-}Half}$ in the presence of MeCbl. **g,** Changes in melting temperatures ($\Delta T_m$, °C) derived from thermal shift assays of MTR$^{FL}$, MTR$^{N\text{-}Half}$, and MTR$^{C\text{-}Half}$ in the presence of various ligands (n=2 for MTR$^{FL}$ with Hcy and MTR$^{C\text{-}Half}$ with SAM; n=3 for all others).

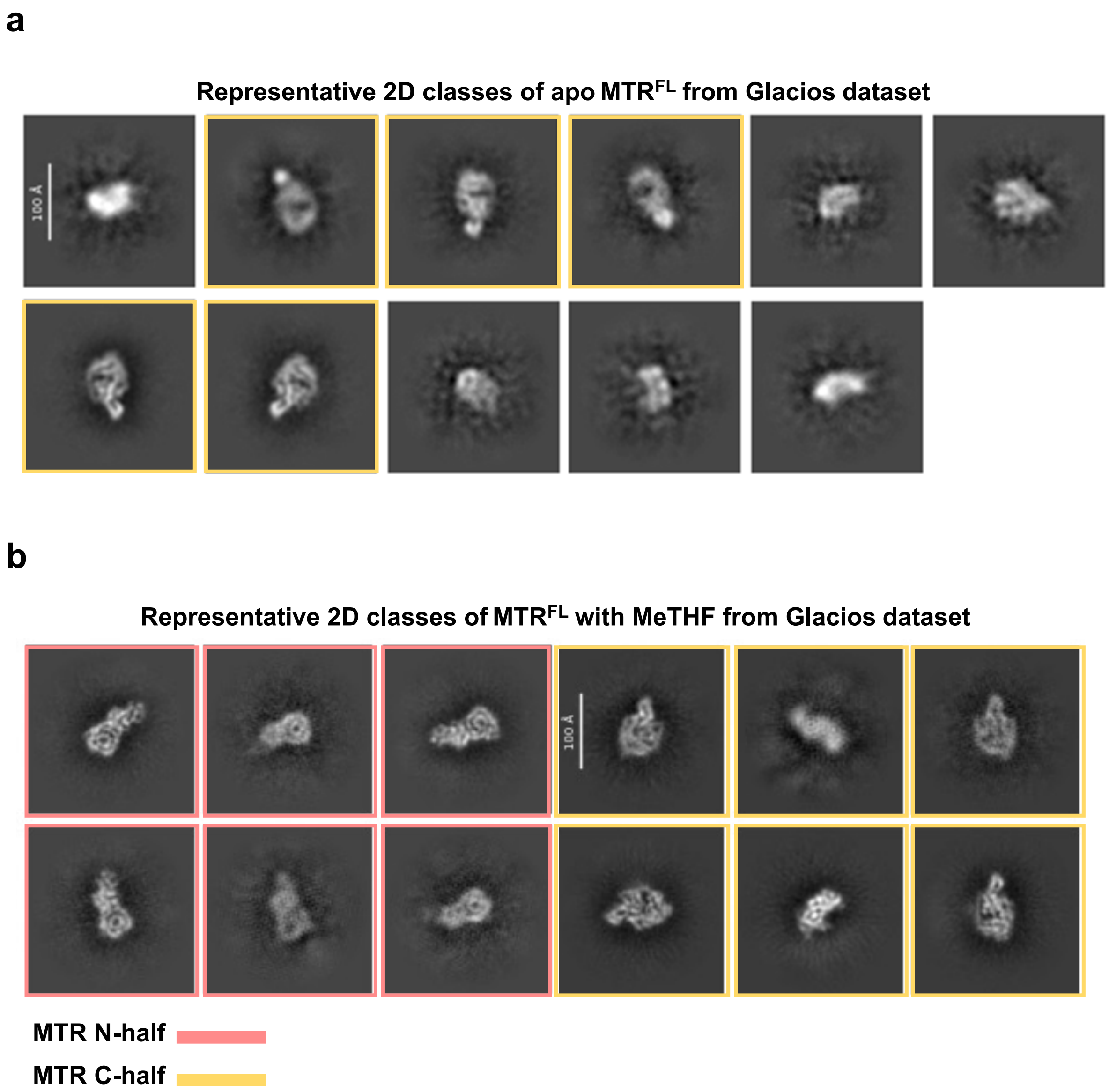


**Supplementary Fig. 2 | Cryo-EM Screening of MTR$^{FL}$ apo (as purified) and with the substrate MeTHF. a**, Representative 2D classes from a Glacios collected dataset of MTR$^{FL}$ apo. Only 2D classes representing the C-half of the protein (pink) were determined. Nonlabelled classes are junk classes. **b**, Representative 2D classes from a Glacios collected screening dataset of MTR$^{FL}$ with MeTHF. 2D classes representing both the N-half (yellow) and C-half (pink) of the protein were determined.

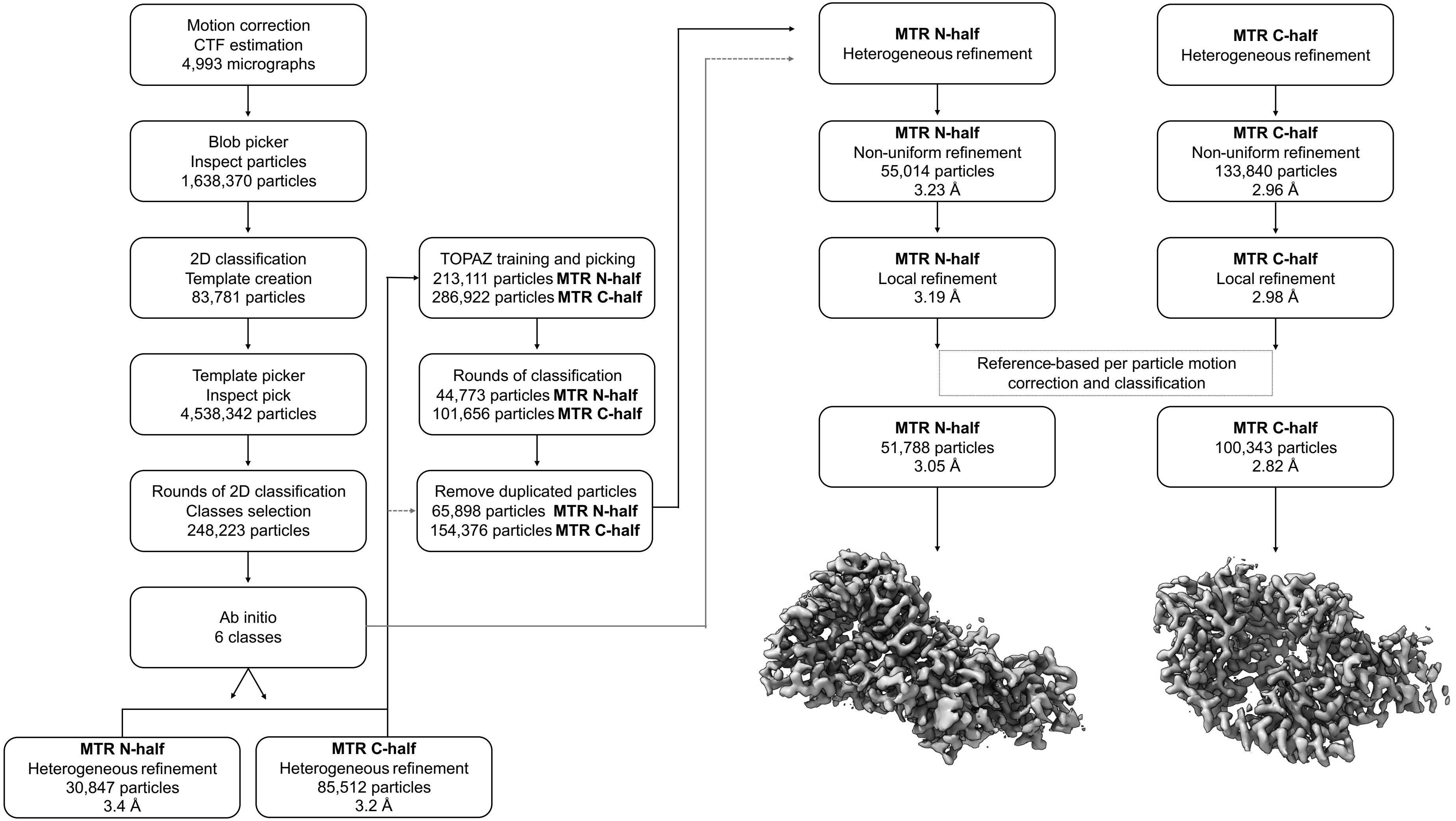


**Supplementary Fig. 3 | Cryo-EM image processing workflow of MTR$^{FL}$ plus MeTHF.** Classes for N-half and C-half were processed separately producing two maps.

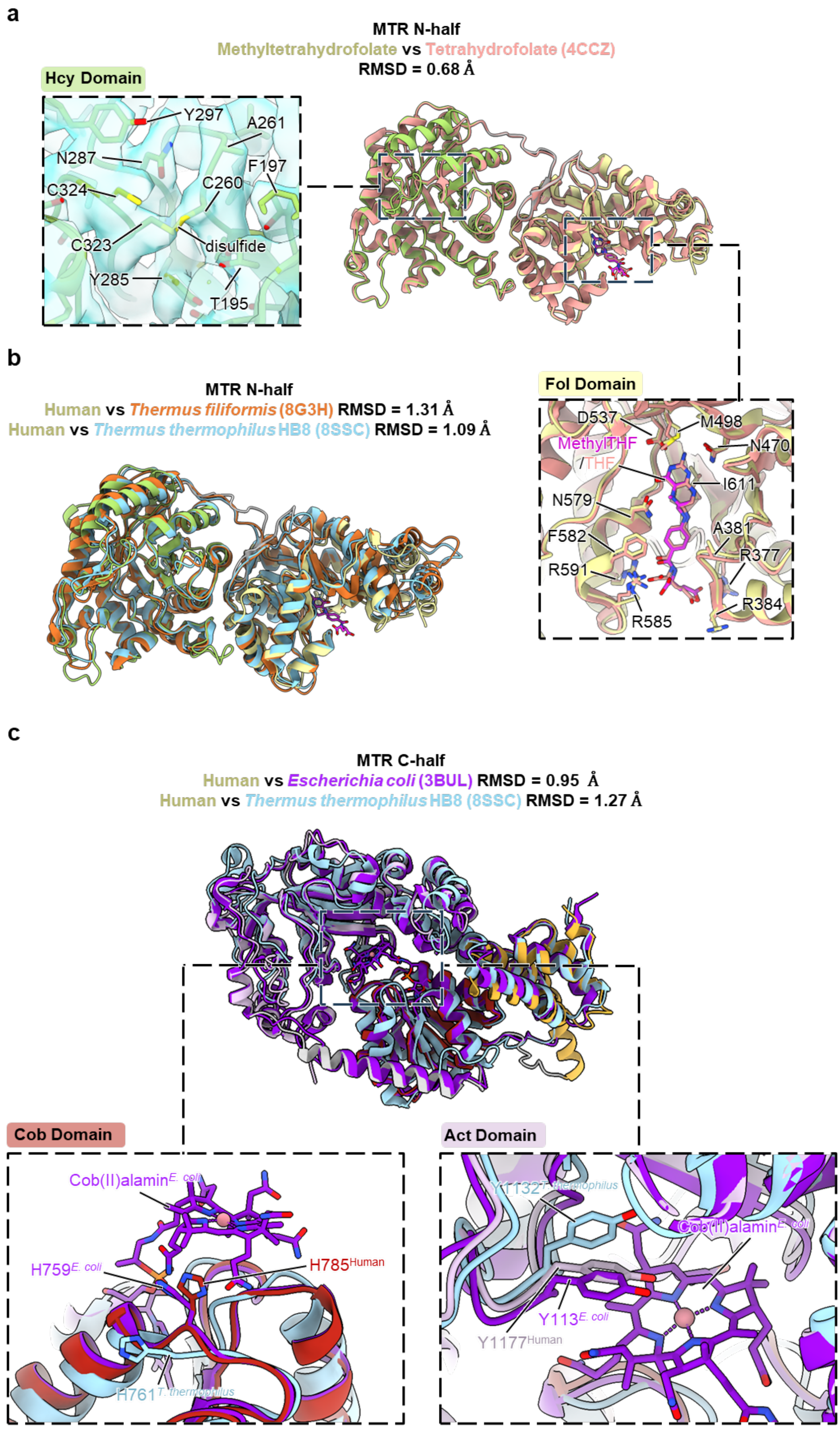
a
MTR N-half
Methyltetrahydrofolate vs Tetrahydrofolate (4CCZ)
RMSD = 0.68 Å
Hcy Domain
Y297
A261
N287
F197
C324
C260
C323
disulfide
Y285
T195
Fol Domain
D537
M498
MethylTHF
/THF
N470
I611
N579
A381
F582
R377
R591
R384
R585
b
MTR N-half
Human vs Thermus filiformis (8G3H) RMSD = 1.31 Å
Human vs Thermus thermophilus HB8 (8SSC) RMSD = 1.09 Å
c
MTR C-half
Human vs Escherichia coli (3BUL) RMSD = 0.95 Å
Human vs Thermus thermophilus HB8 (8SSC) RMSD = 1.27 Å
Cob Domain
Cob(II)alamin E. coli
H759 E. coli
H785 Human
H761 T. thermophilus
Act Domain
Y1132 T. thermophilus
Cob(II)alamin E. coli
Y113 E. coli
Y1177 Human

**Supplementary Fig. 4 | Structural analysis of the N-half and C-half of MTR$^{FL}$ bound to MeTHF structures. a**, Structure alignment of the N-half of MTR$^{FL}$ with MeTHF bound in the folate domain against the crystal structure (PDB 4CCZ) of the equivalent region bound to THF. Weak density is observed in our cryo-EM map for a disulfide bond between Cys260 and Cys323 in the homocysteine domain. Little structural differences are apparent in the folate domain between the MeTHF and THF bound structures. **b**, Structure alignment of the N-half of MTR$^{FL}$ with MeTHF bound in the folate domain against the equivalent regions from the cryo-EM structure of *Thermus filiformis* (PDB 8G3H) and the crystal structure of *Thermus thermophilus* HB8 (PDB 8SSC) MetH proteins. Though highly similar, differences are apparent in the relative locations of α-helices that line one side of the ligand binding site in the folate domain which may be due to the absence of folate in these bacterial structures. **c**, Structure alignment of the C-half of MTR$^{FL}$ without any bound cobalamin against the crystal structure of the cobalamin bound C-half fragment of *Escherichia coli* MetH (PDB 3BUL) and the equivalent region from the crystal structure of *Thermus thermophilus* HB8 (PDB 8SSC). Slight differences are seen in the relative positions of the Cap domains. Though all structures are in the His-off state the three structures show variable positions for this key residue. The position of Tyr1177 is more similar to the cobalamin bound *E. coli* MetH structure than the apo C-half structure from *T. thermophilus*.

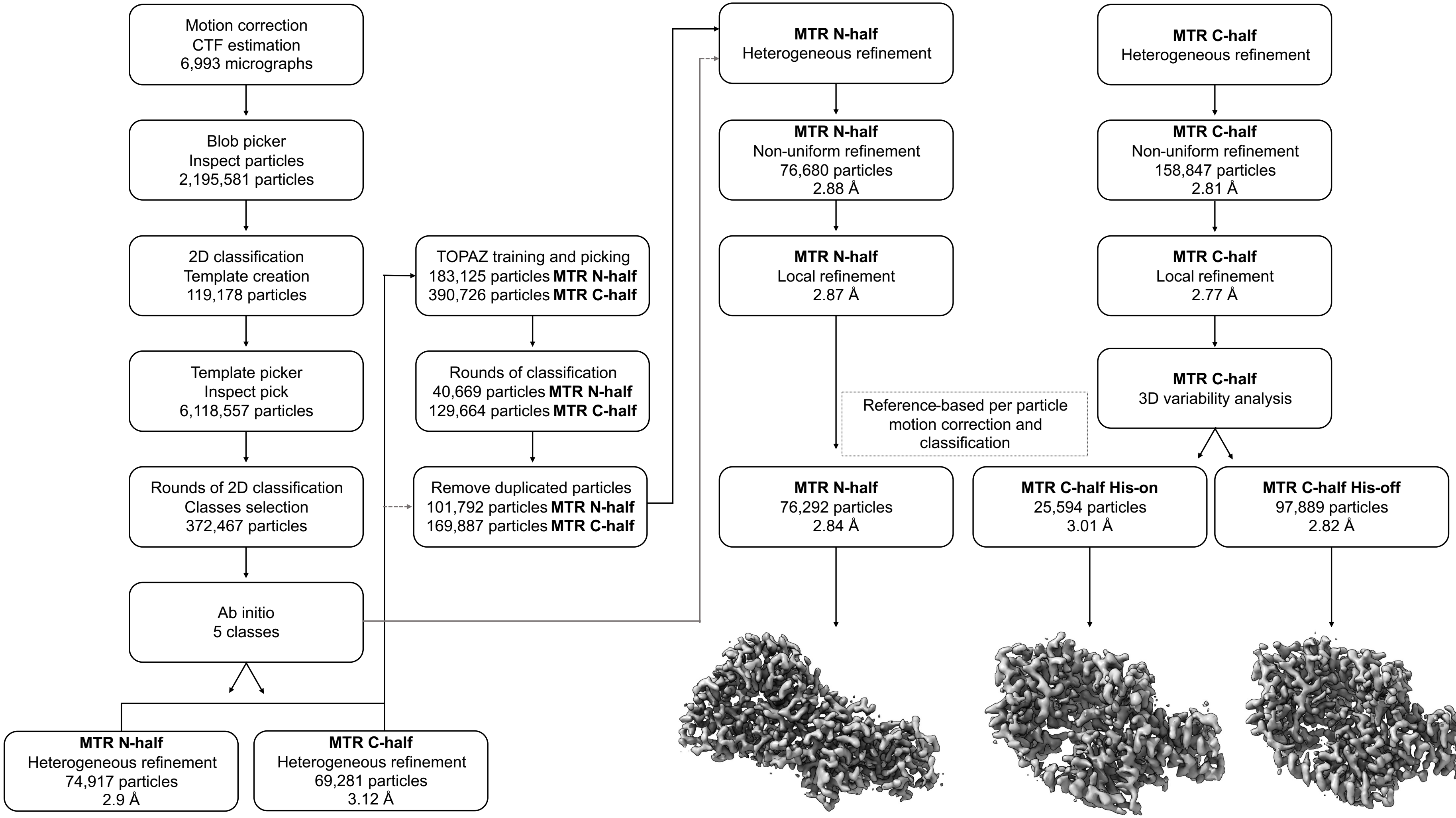


**Supplementary Fig. 5 | Cryo-EM image processing workflow of MTR$^{FL}$ plus MeTHF, SAM, and HOCbl.** Classes for N-half and C-half were processed separately producing three maps, one of the N-half and two of the C-half.

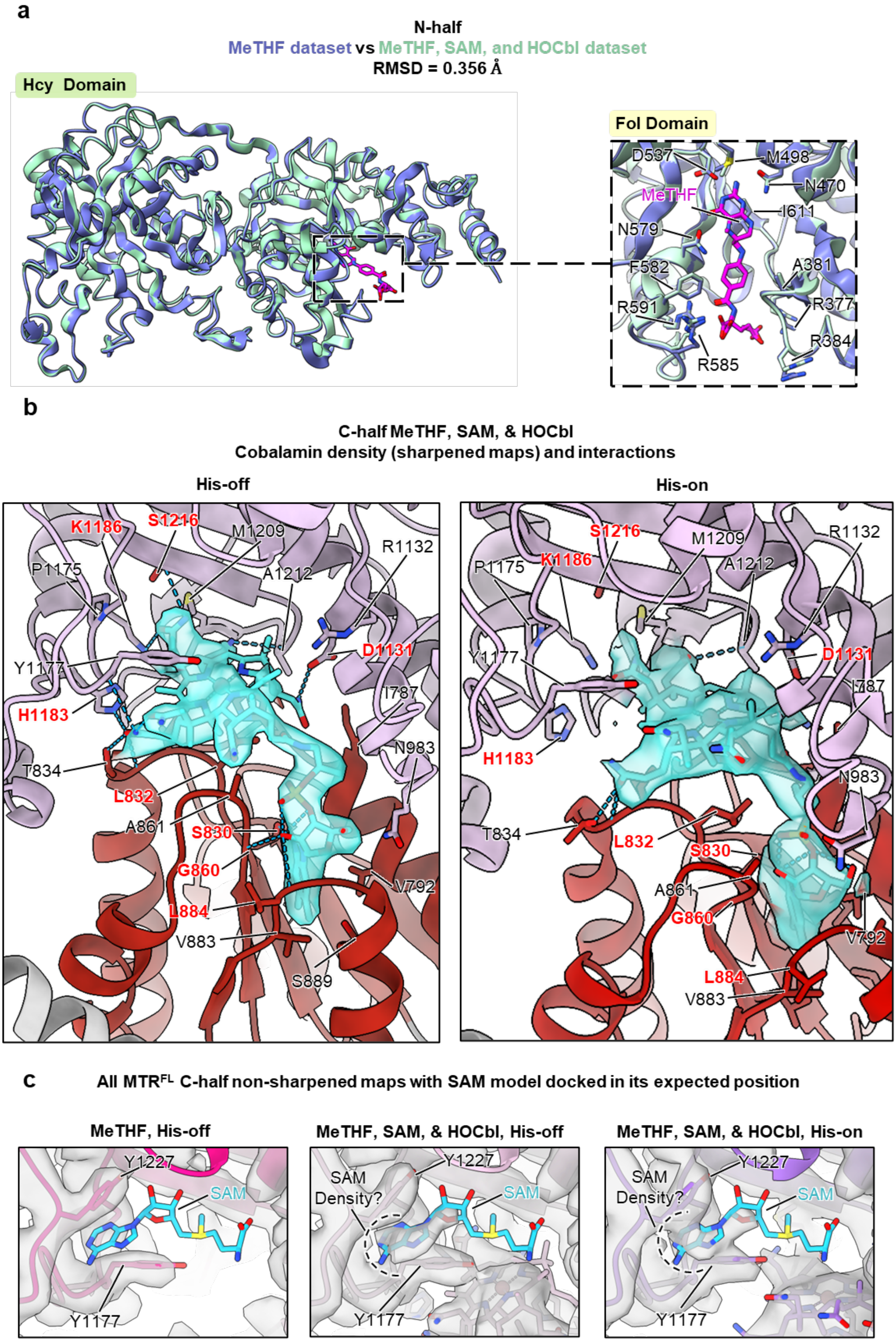
a
N-half
MeTHF dataset vs MeTHF, SAM, and HOCbl dataset
RMSD = 0.356 Å
Hcy Domain
Fol Domain
D537
M498
N470
MeTHF
I611
N579
F582
A381
R591
R377
R384
R585
b
C-half MeTHF, SAM, & HOCbl
Cobalamin density (sharpened maps) and interactions
His-off
His-on
K1186
S1216
M1209
R1132
P1175
A1212
Y1177
D1131
I787
H1183
N983
T834
L832
A861
S830
G860
V792
L884
V883
S889
c
All MTR^FL C-half non-sharpened maps with SAM model docked in its expected position
MeTHF, His-off
MeTHF, SAM, & HOCbl, His-off
MeTHF, SAM, & HOCbl, His-on
Y1227
SAM
SAM Density?
Y1177

**Supplementary Fig. 6 | Structural comparison of ligand binding in MeTHF-only and MeTHF, SAM, and HOCbl datasets.** **a**, Structure alignment of the N-half of $MTR^{FL}$ with MeTHF bound in the folate domain (inset) from MeTHF-only and MeTHF, SAM, and HOCbl dataset. Both structures show identical positioning of MeTHF and conserved interactions with surrounding residues. **b**, Comparison of residues in the cobalamin-binding pocket region of the C-half of $MTR^{FL}$ in the His-on and His-off states. In the His-on state, residues S830, T834, N983, and A1212 interact with HOCbl, while in the His-off state, interactions involve S830, L832, T834, G860, L884, D1131, H1183, K1186, A1212, and S1216. Residues that alter their interaction with the bound cobalamin between His-off and His-on states are highlighted in red. **c**, Non-sharpened C-half cryo-EM density maps of the SAM-binding pocket from the MeTHF-only dataset (left) and from the His-off and His-on reconstructions from the MeTHF, SAM, and HOCbl dataset (middle and right, respectively). The MeTHF-only map shows no SAM density, while a weak density for this ligand is visible in both Cbl bound His-off and His-on maps.

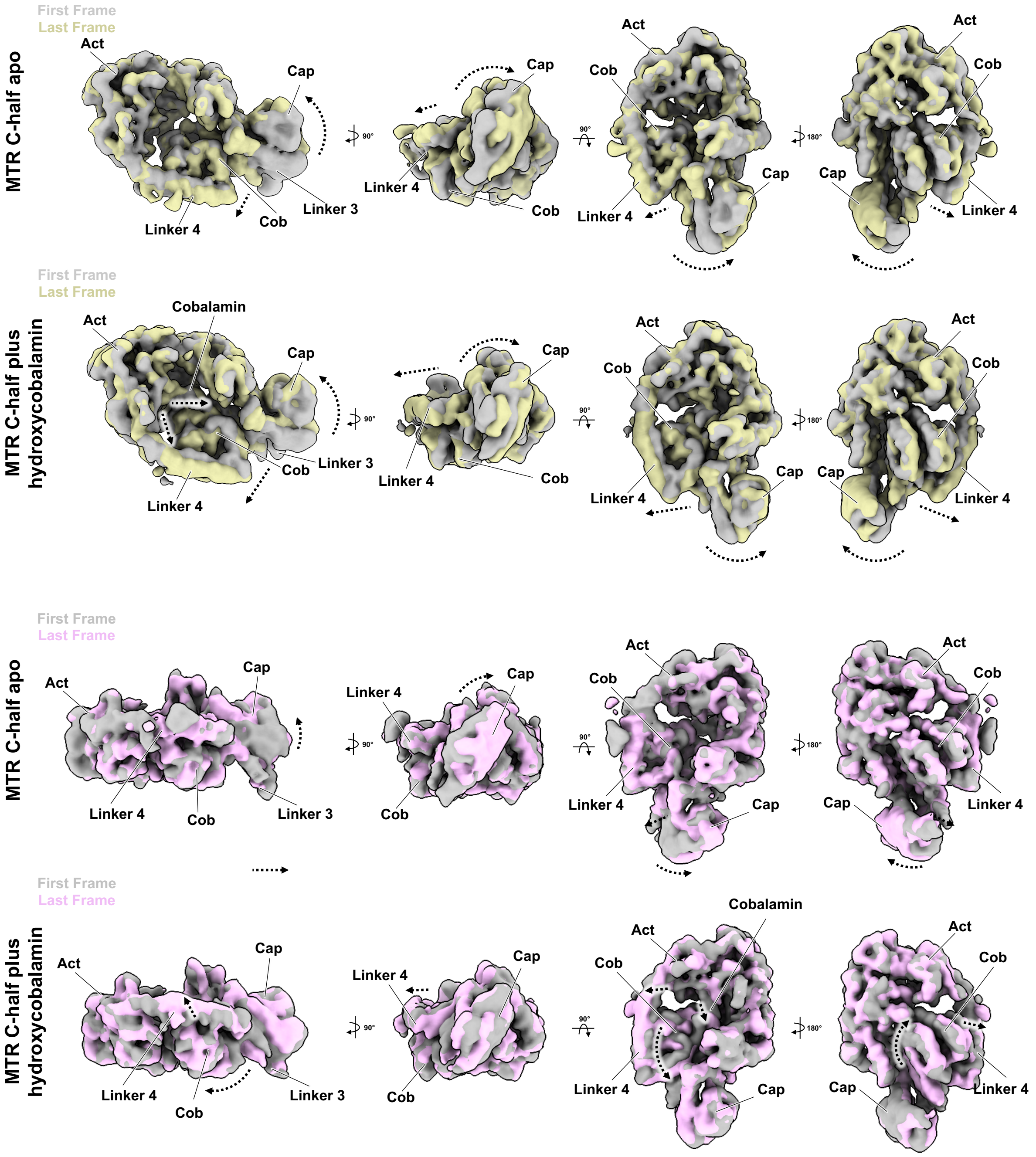


**Supplementary Fig. 7 | 3D variability analysis of the C-half maps of MTR$^{FL}$ demonstrating the flexible Cap domain along with Cob domain movements dependent on the presence of bound cobalamin**. **a and c,** 3D variability analysis of apo MTR C-halfviewed from different orientations, revealing motion of the Cob and Cap domains and adjacent regions. **b and d,** Corresponding analysis of HOCbl-bound MTR C-half showing the flexibility upon co-factor binding.

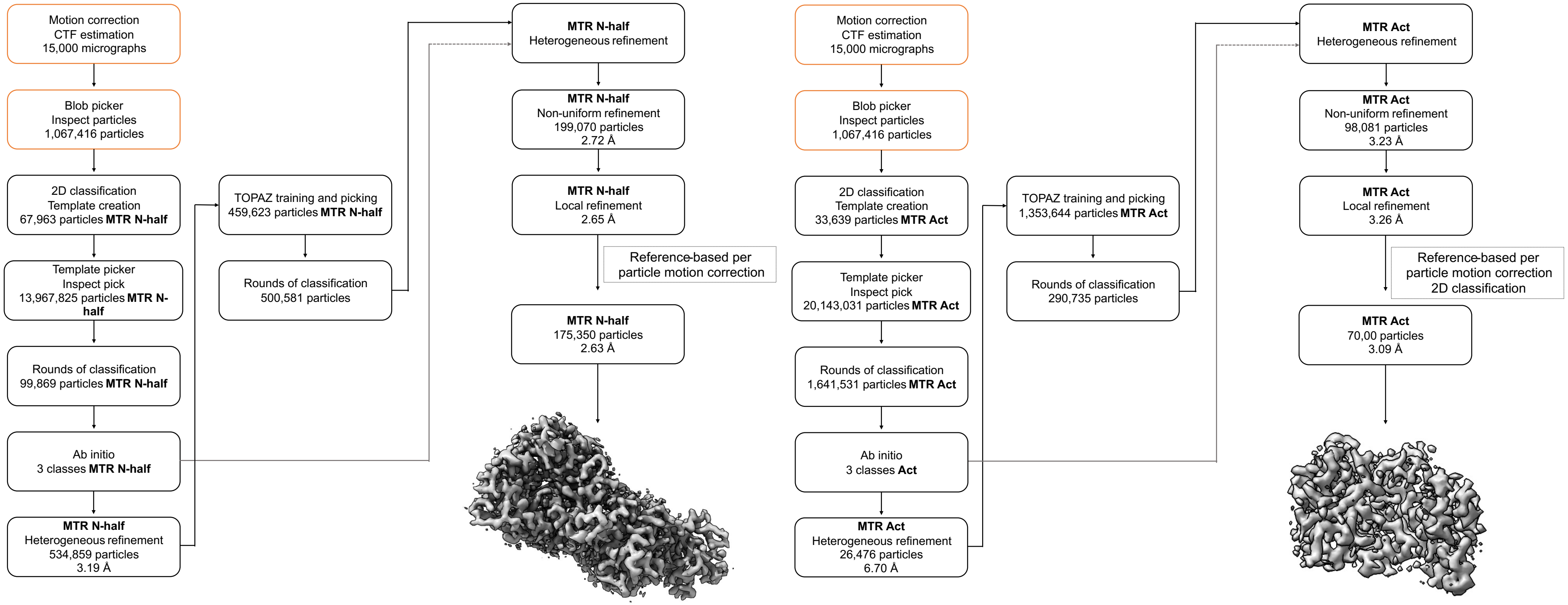


**Supplementary Fig. 8 | Cryo-EM image processing workflow of MTR$^{FL}$ plus MeCbl.** Classes for the MTR N-half and Act domain were processed separately after blob picking (orange highlight) producing the two separate maps.

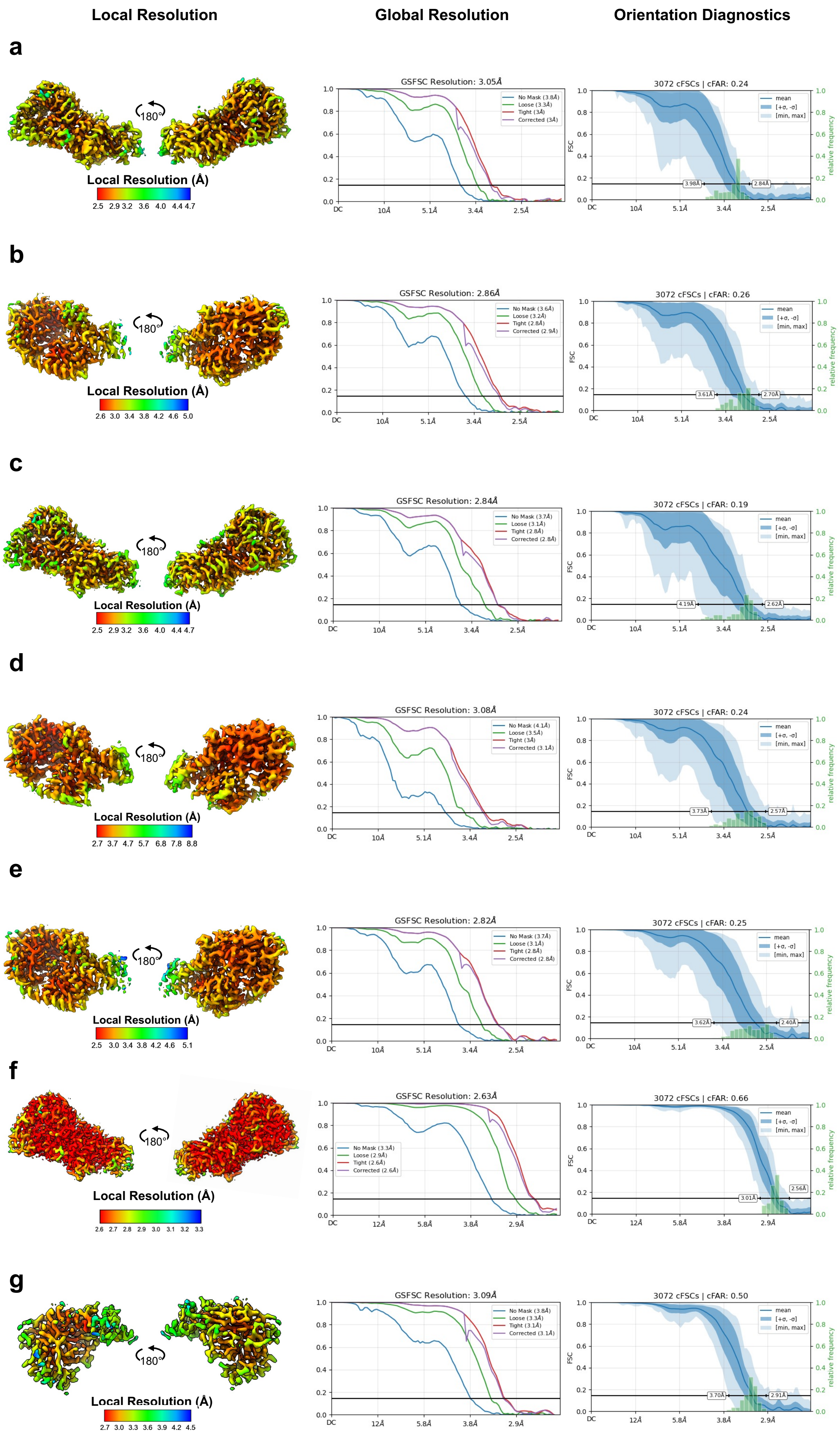
Local Resolution
Global Resolution
Orientation Diagnostics
a
Local Resolution (Å)
GSFSC Resolution: 3.05Å
3072 cFSCs | cFAR: 0.24
b
Local Resolution (Å)
GSFSC Resolution: 2.86Å
3072 cFSCs | cFAR: 0.26
c
Local Resolution (Å)
GSFSC Resolution: 2.84Å
3072 cFSCs | cFAR: 0.19
d
Local Resolution (Å)
GSFSC Resolution: 3.08Å
3072 cFSCs | cFAR: 0.24
e
Local Resolution (Å)
GSFSC Resolution: 2.82Å
3072 cFSCs | cFAR: 0.25
f
Local Resolution (Å)
GSFSC Resolution: 2.63Å
3072 cFSCs | cFAR: 0.66
g
Local Resolution (Å)
GSFSC Resolution: 3.09Å
3072 cFSCs | cFAR: 0.50
180°

**Supplementary Fig. 9 | Cryo-EM map quality metrics for all reported cryo-EM structures in this study.** For each map the local resolution (left panel), the global resolution FSC curves (middle panel), and orientation diagnostic plots (right panel) are shown. **a**, N-half structure from $MTR^{FL}$ plus MeTHF. **b,** C-half structure from $MTR^{FL}$ plus MeTHF. **c**, N-half structure from $MTR^{FL}$ plus MeTHF, SAM, and HOCbl. **d**, C-half His-on structure from $MTR^{FL}$ plus MeTHF, SAM, and HOCbl. **e**, C-half His-off structure from $MTR^{FL}$ plus MeTHF, SAM, and HOCbl. **f**, $MTR^{N\text{-}Half}$ structure from $MTR^{FL}$ plus MeCbl. **g**, Act domain structure from $MTR^{FL}$ plus MeCbl.

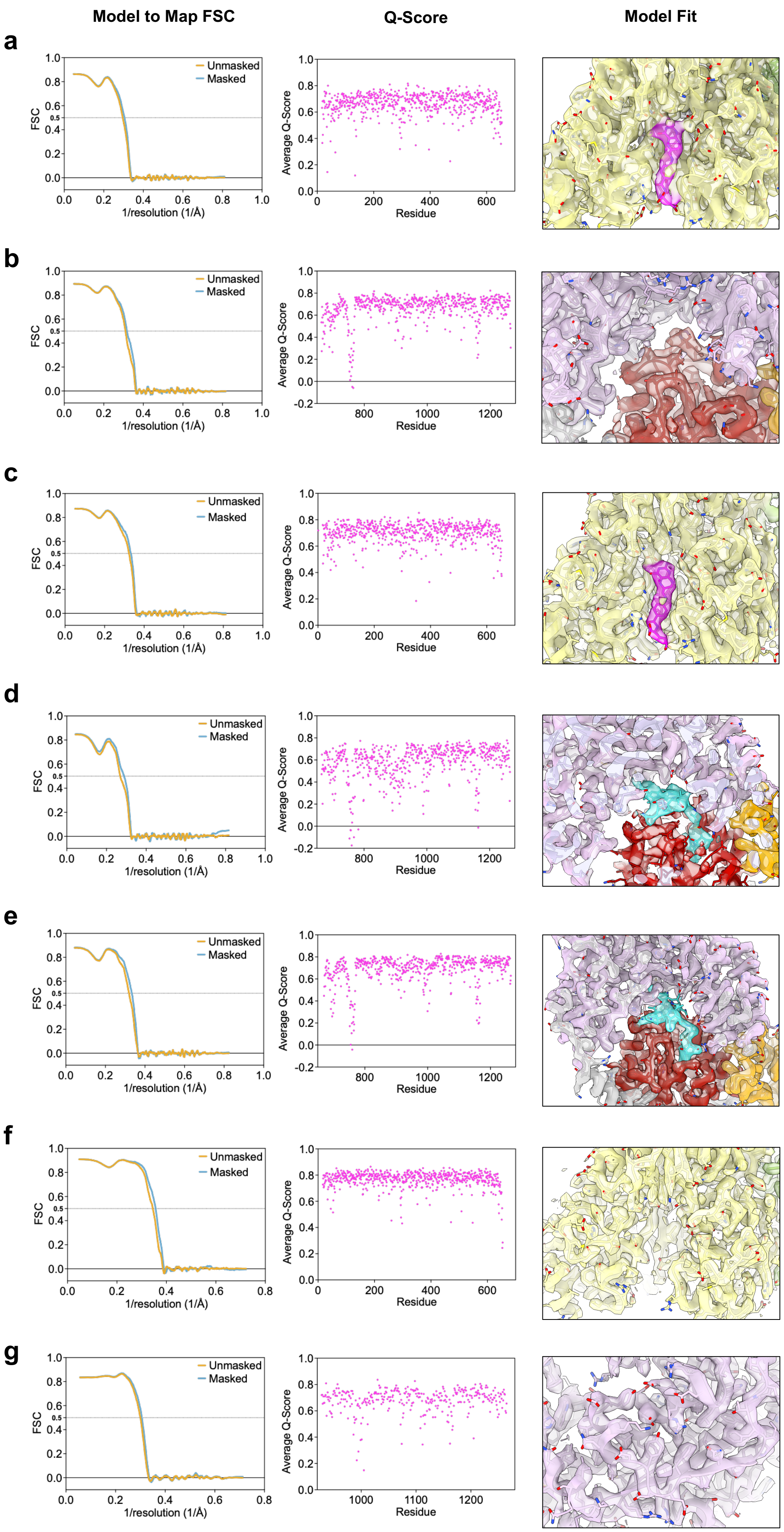

Model to Map FSC
Q-Score
Model Fit
a
b
c
d
e
f
g
Unmasked
Masked
FSC
1/resolution (1/Å)
Average Q-Score
Residue

**Supplementary Fig. 10 | Model fit quality metrics for all reported cryo-EM structures in this study.** For each structure the model to map FSC curves (left panel), per residue average Q-score (middle panel), and model fit quality (right panel) are shown. **a**, N-half structure from $MTR^{FL}$ plus MeTHF. **b,** C-half structure from $MTR^{FL}$ plus MeTHF. **c**, N-half structure from $MTR^{FL}$ plus MeTHF, SAM, and HOCbl. **d**, C-half His-on structure from $MTR^{FL}$ plus MeTHF, SAM, and HOCbl. **e**, C-half His-off structure from $MTR^{FL}$ plus MeTHF, SAM, and HOCbl. **f**, N-half structure from $MTR^{FL}$ plus MeCbl. **g**, Act domain structure from $MTR^{FL}$ plus MeCbl.

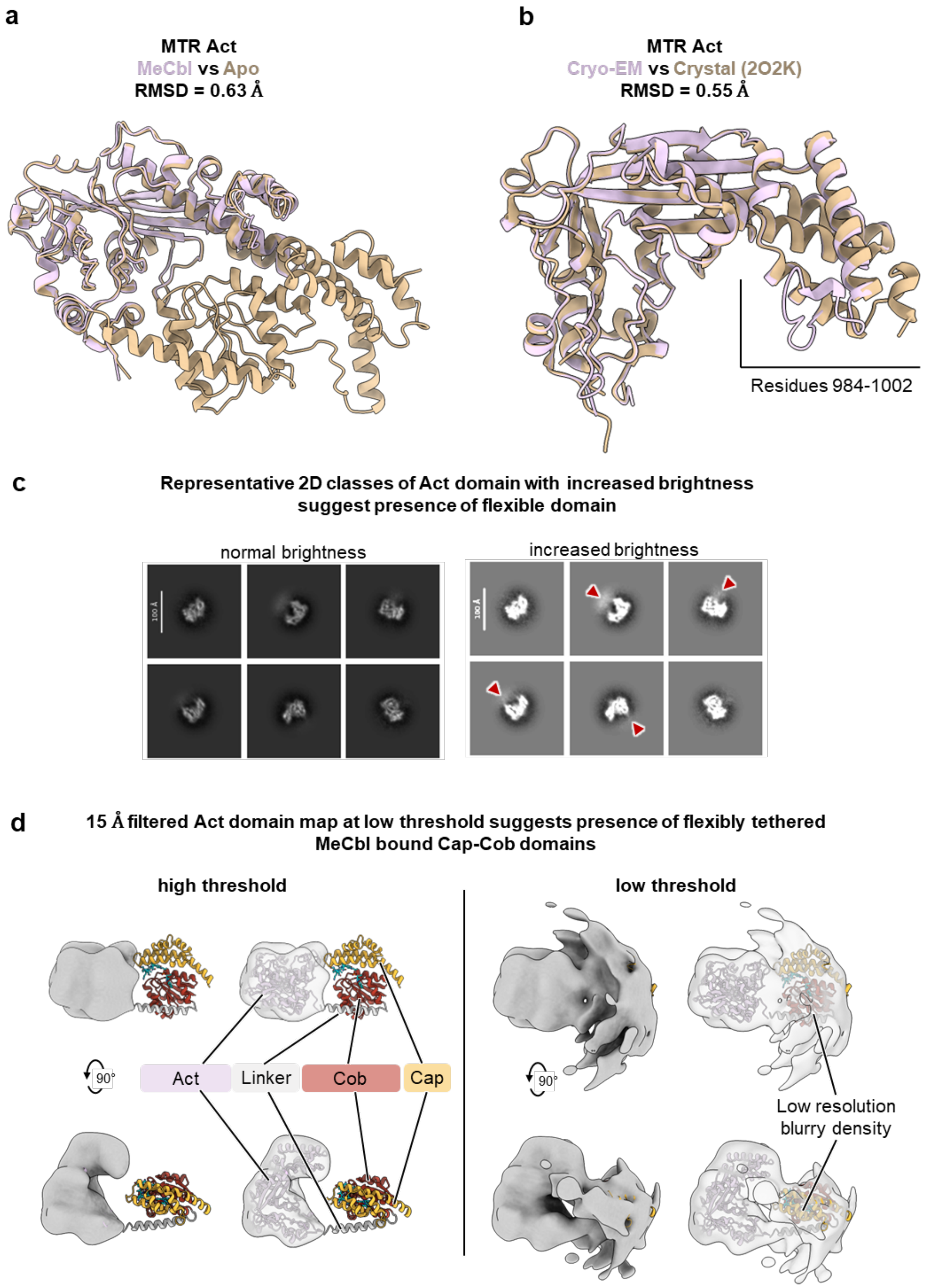


**Supplementary Fig. 11 | Structural analysis of the Act domain from MeCbl dataset. a**, Structure alignment of the Act domain structure determined from the MeCbl dataset against the entire C-half structure without bound Cbl. **b**, Structure alignment of the Act domain structure determined from the MeCbl dataset with the corresponding crystal structure (PDB 2O2K). **c,** Representative 2D classes averages from the MeCbl dataset used for the Act domain reconstruction. The red mark regions with increased blur, suggesting the presence of

Cap and Cob domains. **d,** Left, possible position of the Cap and Cob domains relative to the Act domain volume, shown at high threshold. Right, the Act domain displayed at low threshold reveals additional low-resolution density, suggesting the presence of the other two domains.

**AF3: MTR$^{FL}$ plus MMACHC$^{FL}$** ipTM = 0.18 pTM = 0.49

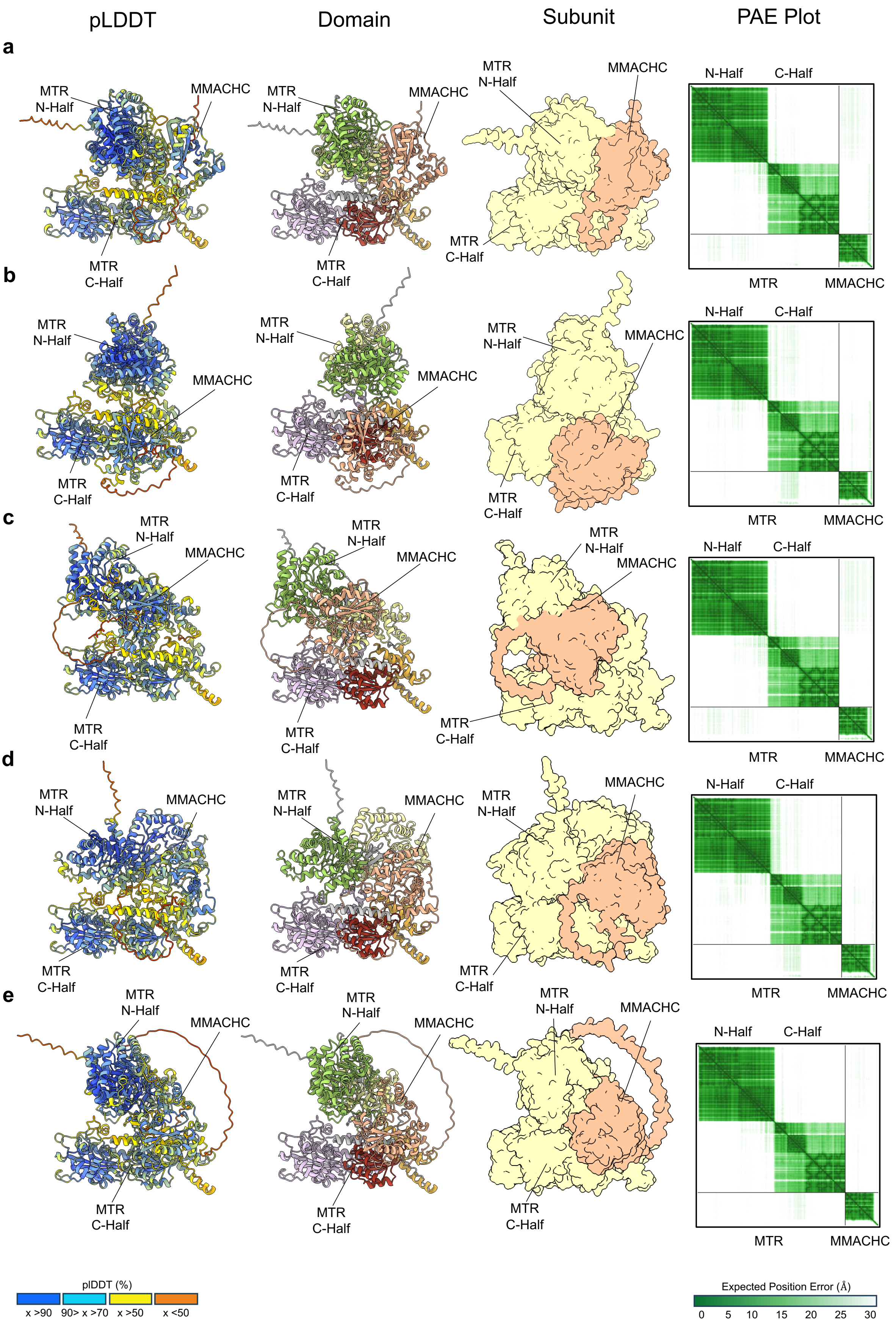

**Supplementary Fig. 12 | AF3 prediction models of $MTR^{FL}$ and $MMACHC^{FL}$.** Predicted complexes are shown coloured by per-residue confidence scores (pLDDT, far left), by proteins domains (middle left), and by subunit (middle right). The corresponding predicted aligned error (PAE) plot for each model are shown to the far right, indicating the inter-domain confidence and the overall accuracy of the models. In all five prediction models, MMACHC binds in different regions of $MTR^{FL}$, suggesting that these proteins do not form a stable complex. The low ipTM score (0.18) further supports that these proteins may not form a complex. **a to e,** are the five predicted complex models generated using the sequence of $MTR^{FL}$ and $MMACHC^{FL}$ representative of one AF3 run.

**AF3: MTR$^{FL}$ plus MMADHC$^{FL}$** ipTM = 0.55 pTM = 0.52

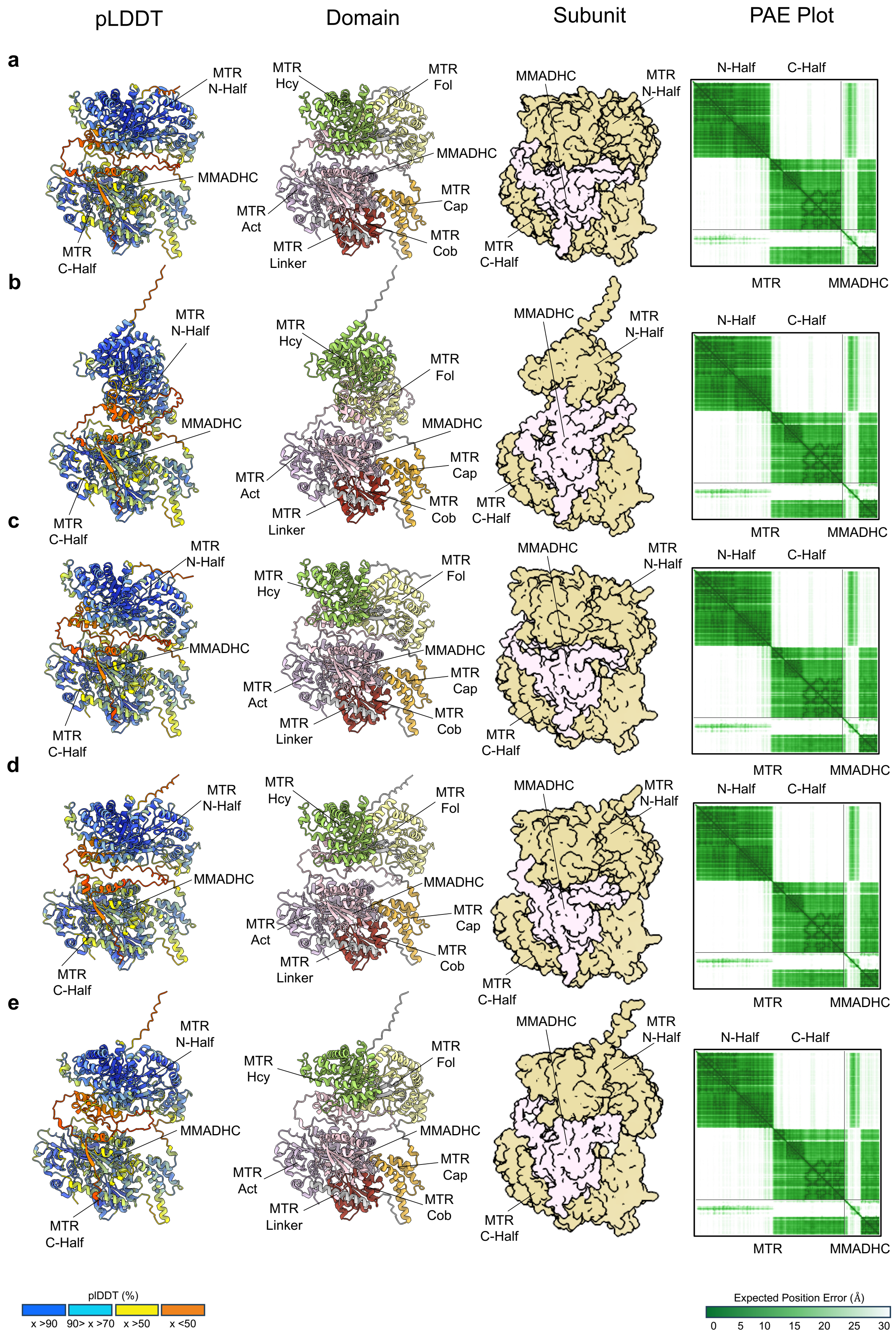

**Supplementary Fig. 13 | AF3 prediction models of MTR$^{FL}$ and MMADHC$^{FL}$.** Predicted complexes are shown coloured by per-residue confidence scores (pLDDT, far left), by proteins domains (middle left), and by subunit (middle right). The corresponding predicted aligned error (PAE) plot for each model are shown to the far right, indicating the inter-domain confidence and the overall accuracy of the models. In all five prediction models, MMADHC binds near the cobalamin-binding pocket on MTR$^{C\text{-}Half}$ in a consistent orientation, suggesting these proteins may form a stable complex. The moderate ipTM score (0.55) further supports that these proteins may form a complex. **a to e,** are the five predicted complex models generated using the sequence of MTR$^{FL}$ and MMADHC$^{FL}$ representative of one AF3 run.

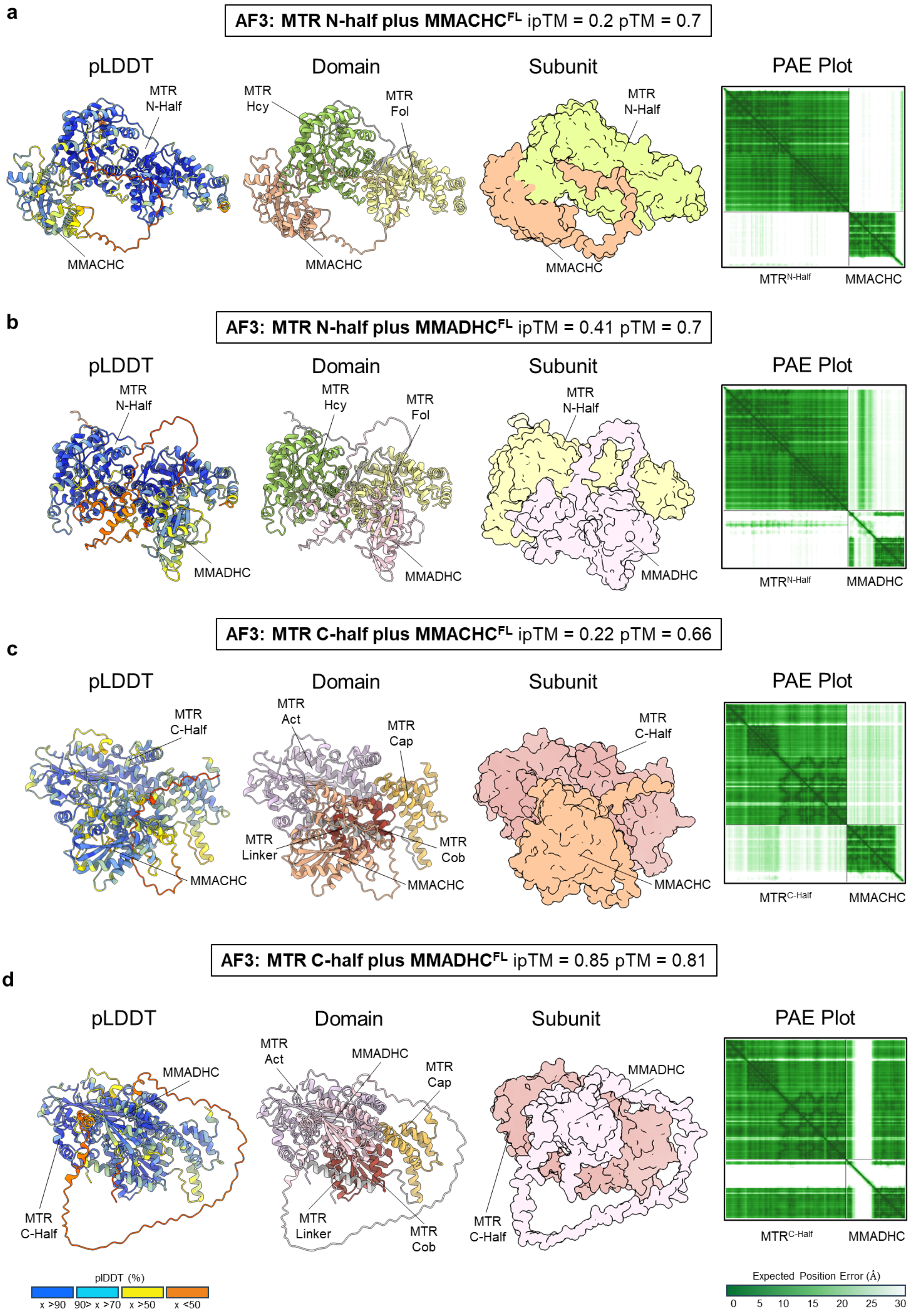

a
AF3: MTR N-half plus MMACHC$^{FL}$ ipTM = 0.2 pTM = 0.7
pLDDT
Domain
Subunit
PAE Plot
MTR N-Half
MMACHC
MTR Hcy
MTR Fol
MTR$^{N-Half}$
b
AF3: MTR N-half plus MMADHC$^{FL}$ ipTM = 0.41 pTM = 0.7
MMADHC
c
AF3: MTR C-half plus MMACHC$^{FL}$ ipTM = 0.22 pTM = 0.66
MTR C-Half
MTR Act
MTR Cap
MTR Linker
MTR Cob
MTR$^{C-Half}$
d
AF3: MTR C-half plus MMADHC$^{FL}$ ipTM = 0.85 pTM = 0.81
plDDT (%)
x >90
90> x >70
x >50
x <50
Expected Position Error (Å)
0
5
10
15
20
25
30

**Supplementary Fig. 14 | AF3 prediction models of MTR N-half and MTR C-half with MMACHC$^{FL}$ or MMADHC$^{FL}$.** Predicted complexes are shown coloured by per-residue confidence scores (pLDDT, far left), by proteins domains (middle left), and by subunit (middle right). The corresponding predicted aligned error (PAE) plot for each model are shown to the far right, indicating the inter-domain confidence and the overall accuracy of the models. Shown structures are representative of the five predicted models from one AF3 run. **a,** Predicted model of MTR N-half and MMACHC$^{FL}$ showing a low ipTM score (0.2), suggesting no stable interaction. **b,** Predicted model of MTR N-half and MMADHC$^{FL}$ showing a low ipTM score (0.41), suggesting no stable interaction. **c,** Predicted model of MTR C-half and MMACHC$^{FL}$ showing a low ipTM score (0.22), suggesting no stable interaction. **d,** Predicted of MTR C-half and MMADHC$^{FL}$ showing a high ipTM score (0.85), suggesting that these proteins might form a stable complex.

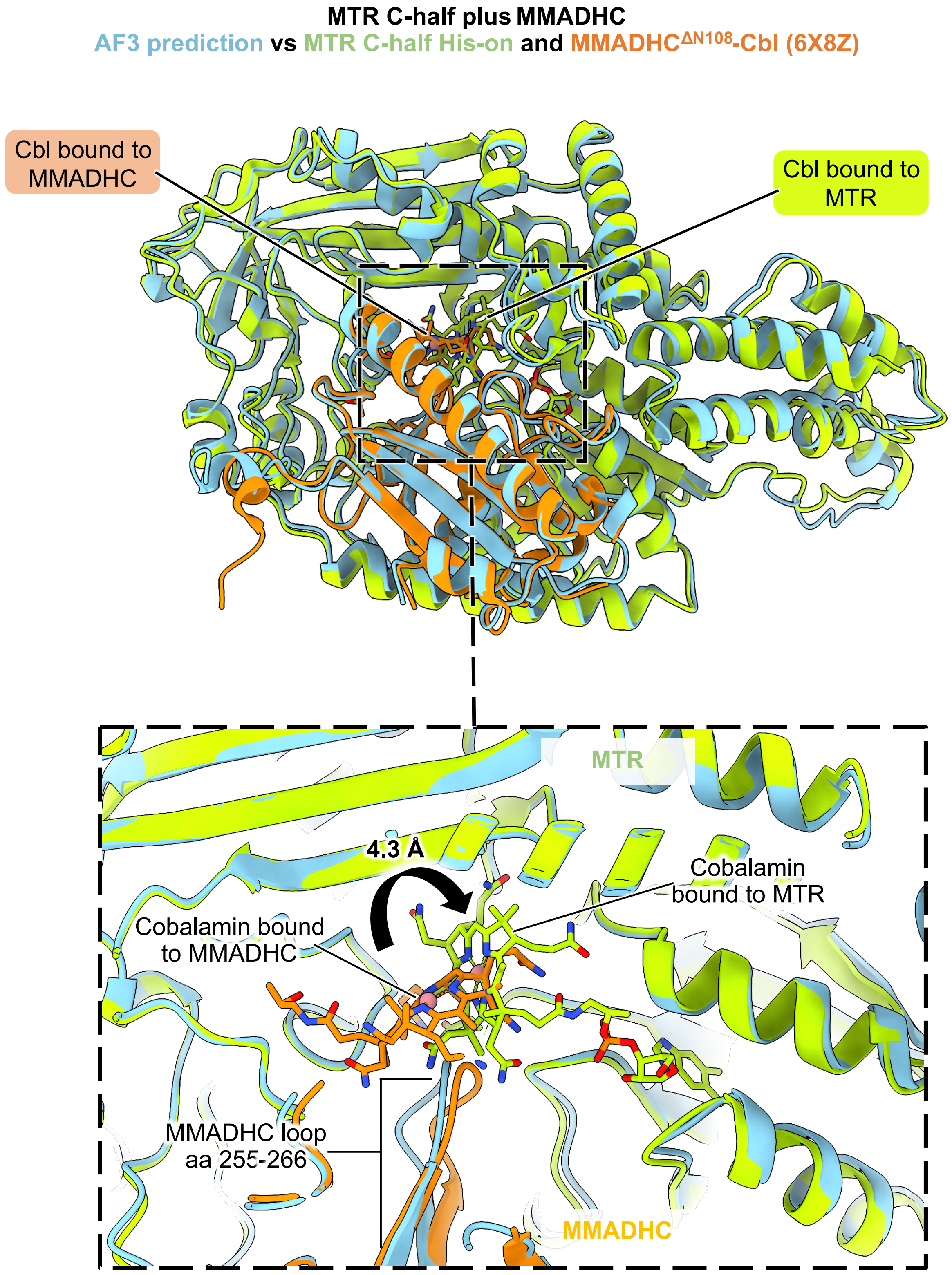


**Supplementary Fig. 15 | Structural alignment of the predicted MTR$^{FL}$+MMADHC$^{FL}$ complex with experimental Cbl-bound structures.** Superimposition of the AF3 predicted MTR$^{FL}$-MMADHC$^{FL}$ complex against our cryo-EM structure of MTR C-half His-off and crystal structure of the Cbl-bound MMADHC$^{\Delta N108}$ (6X8Z) show the relative positioning of Cbl in both proteins. The alignment indicates that Cbl may move 4.3 Å towards its binding pocket in the MTR Cob domain to be offloaded (inset).

**a**

**Replicate pulldown of MTR$^{FL}$ against GST-MMACHC and GST-MMADHC$^{\Delta N}$**

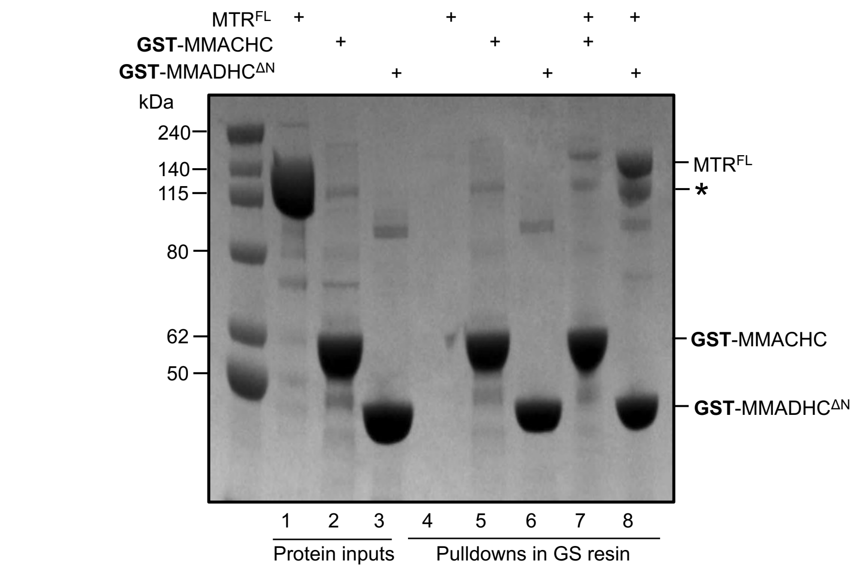


**b**

**Replicate pulldown of MTR$^{N\text{-}Half}$ and MTR$^{C\text{-}Half}$ against GST-MMACHC and GST-MMADHC$^{\Delta N}$**

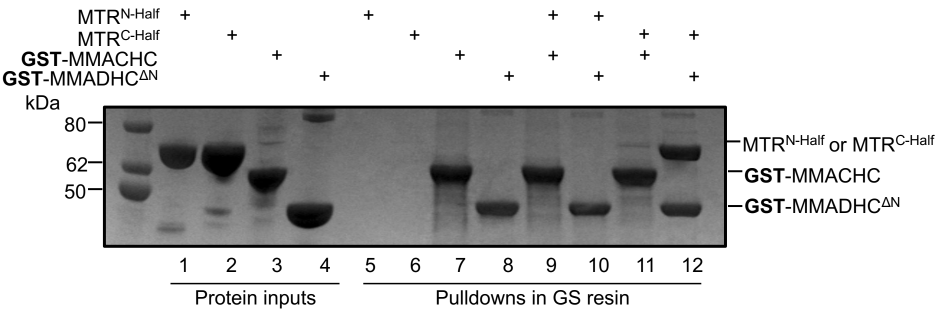


**c**

**Replicate runs of cobalamin binding of MTR$^{FL}$ and loading by MMADHC$^{\Delta N}$**

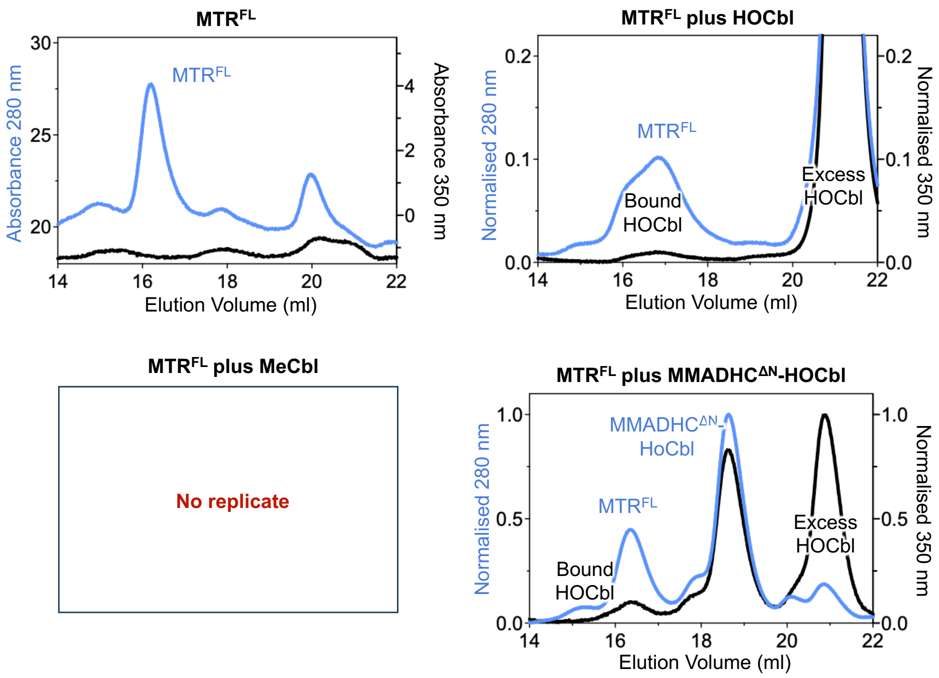

**Supplementary Fig. 16 | Replicates of affinity-based pulldown and Cbl loading experiments. a,** Replicate GS resin affinity pulldown of $MTR^{FL}$ against GST-MMACHC and GST-$MMADHC^{\Delta N}$. Lanes 1–3 represent the protein inputs used for the assay. Lanes 4–6 represent the elution fractions of each individual protein incubated with the resin to verify their ability to bind the resin. Lanes 7 and 8 represent the pulldown of $MTR^{FL}$ with GST-MMACHC and the pulldown of $MTR^{FL}$ with GST-MMADHC, respectively. The asterisk indicates a degradation product of $MTR^{FL}$. **b,** Replicate GS resin affinity pulldown of $MTR^{N\text{-}Half}$ and $MTR^{C\text{-}Half}$ against GST-MMACHC and GST-$MMADHC^{\Delta N}$. Lanes 1–4 represent the protein inputs used for the assay. Lanes 5–8 represent the elution fractions of each individual protein incubated with the resin to verify their ability to bind the resin. Lanes 9 and 10 represent the pulldown of $MTR^{N\text{-}Half}$ with GST-MMACHC and GST-MMADHC, respectively. Lanes 11 and 12 represent the pulldown of $MTR^{C\text{-}Half}$ with GST-MMACHC and GST-MMADHC, respectively. **c,** Replicate runs of size-exclusion chromatography of cobalamin loading assay showing $MTR^{FL}$, $MTR^{FL}$ plus HOCbl, and $MTR^{FL}$ plus HOCbl-loaded $MMADHC^{\Delta N}$ monitored at 280 nm (blue) and 350 nm (black for HOCbl). Binding of MeCbl by $MTR^{FL}$ was only done once and is presented in the main figure.

.

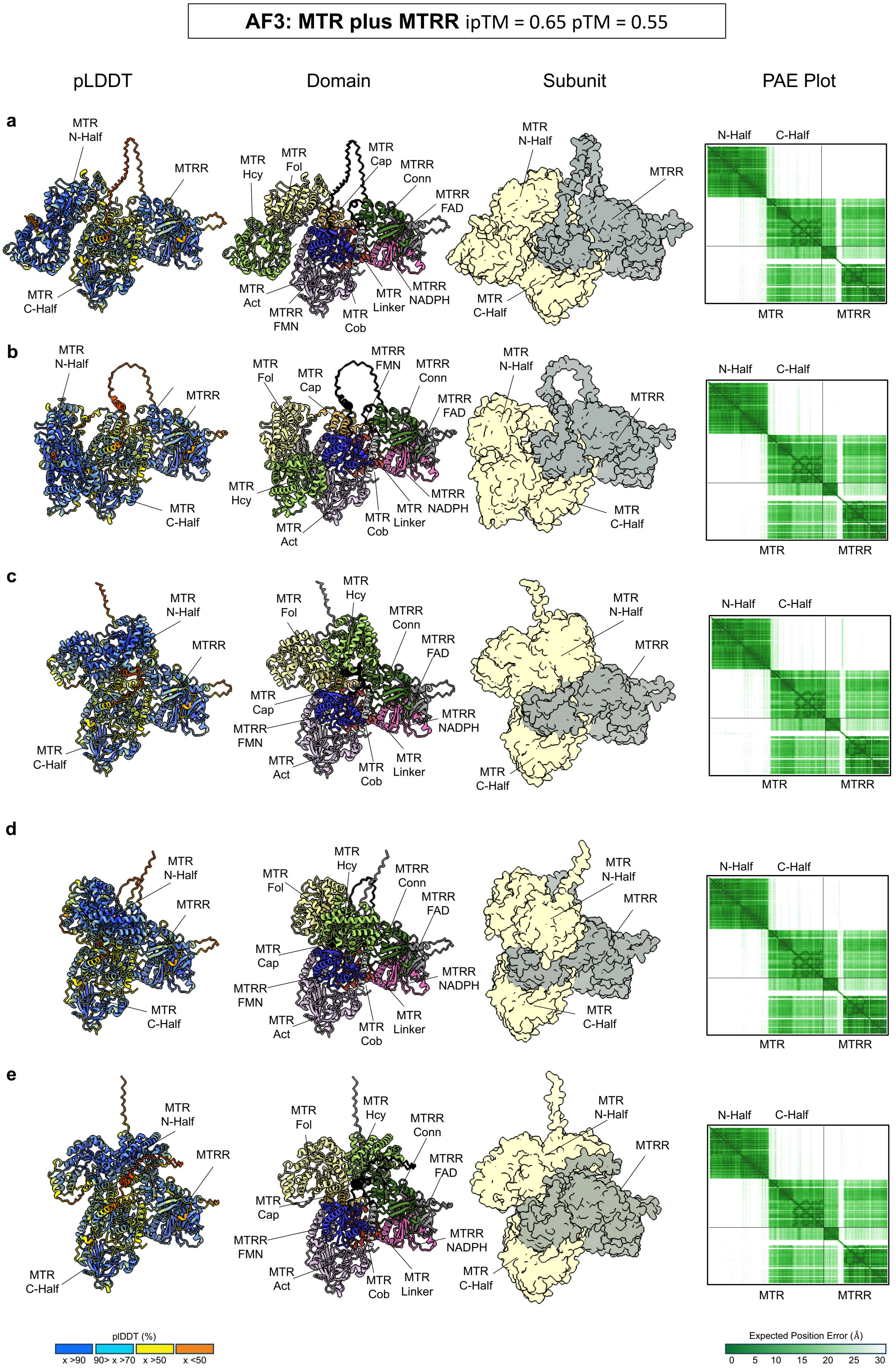
AF3: MTR plus MTRR ipTM = 0.65 pTM = 0.55
pLDDT
Domain
Subunit
PAE Plot
a
b
c
d
e
MTR N-Half
MTR C-Half
MTRR
MTR Hcy
MTR Fol
MTR Cap
MTRR Conn
MTRR FAD
MTR Act
MTRR FMN
MTR Cob
MTR Linker
MTRR NADPH
N-Half
C-Half
MTR
MTRR
pIDDT (%)
x >90
90> x >70
x >50
x <50
Expected Position Error (Å)
0
5
10
15
20
25
30

**Supplementary Fig. 17 | AF3 prediction models of $MTR^{FL}$ and $MTRR^{FL}$.** Predicted complexes are shown coloured by per-residue confidence scores (pLDDT, far left), by proteins domains (middle left), and by subunit (middle right). The corresponding predicted aligned error (PAE) plot for each model are shown to the far right, indicating the inter-domain confidence and the overall accuracy of the models. In all five prediction models, $MTRR^{FL}$ binds to $MTR^{FL}$ in a consistent orientation, suggesting these proteins may form a stable complex. The moderate ipTM score (0.65) further supports that these proteins may form a complex. **a to e,** are the five predicted complex models generated using the sequence of $MTR^{FL}$ and $MTRR^{FL}$ representative of one AF3 run.

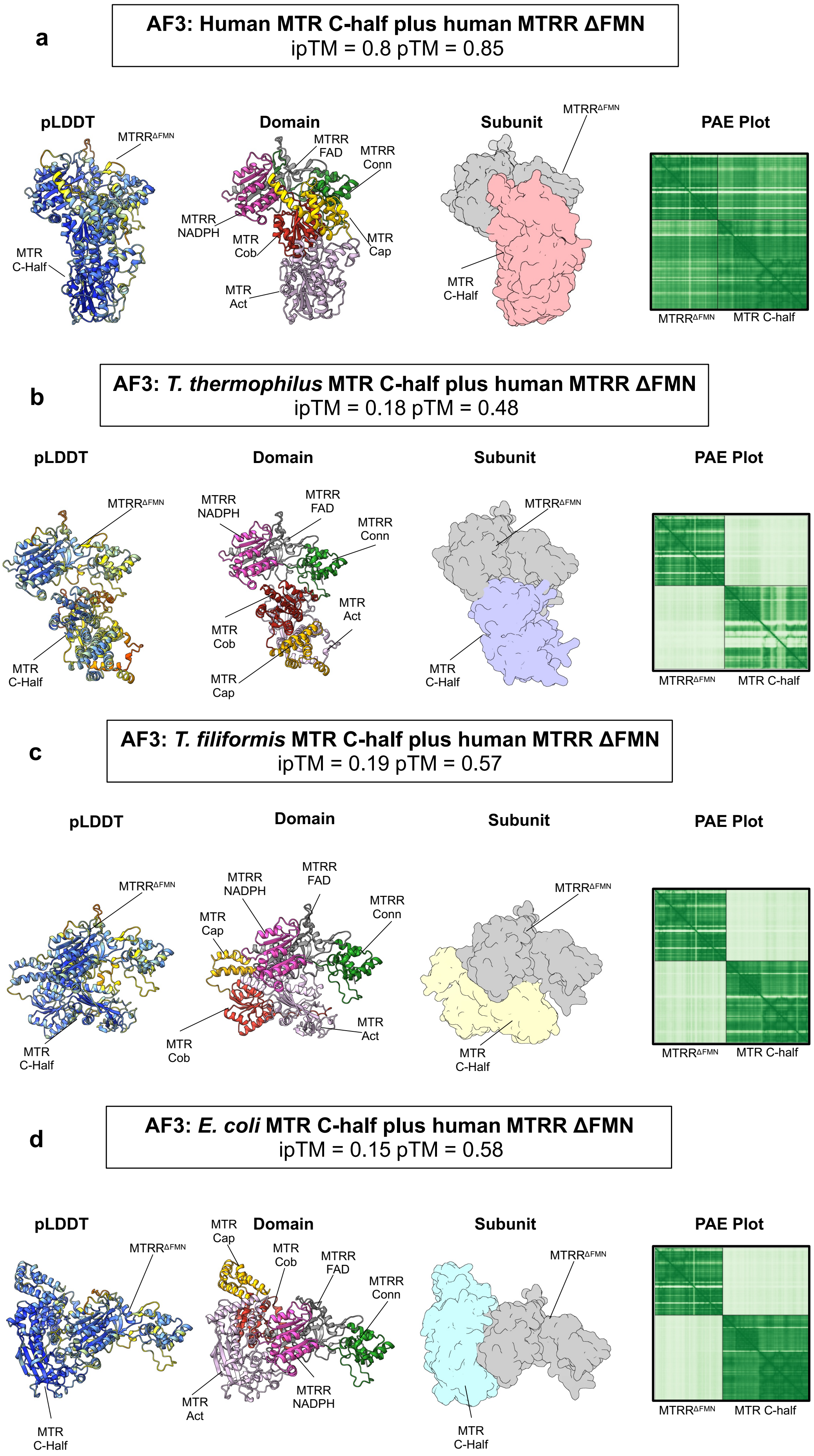
a
AF3: Human MTR C-half plus human MTRR ΔFMN
ipTM = 0.8 pTM = 0.85
pLDDT
Domain
Subunit
PAE Plot
MTRR$^{\Delta FMN}$
MTR C-Half
MTRR FAD
MTRR Conn
MTRR NADPH
MTR Cob
MTR Cap
MTR Act
MTRR$^{\Delta FMN}$
MTR C-half
b
AF3: *T. thermophilus* MTR C-half plus human MTRR ΔFMN
ipTM = 0.18 pTM = 0.48
pLDDT
Domain
Subunit
PAE Plot
MTRR$^{\Delta FMN}$
MTR C-Half
MTRR NADPH
MTRR FAD
MTRR Conn
MTR Act
MTR Cob
MTR Cap
MTRR$^{\Delta FMN}$
MTR C-half
c
AF3: *T. filiformis* MTR C-half plus human MTRR ΔFMN
ipTM = 0.19 pTM = 0.57
pLDDT
Domain
Subunit
PAE Plot
MTRR$^{\Delta FMN}$
MTR C-Half
MTRR NADPH
MTRR FAD
MTR Cap
MTRR Conn
MTR Act
MTR Cob
MTRR$^{\Delta FMN}$
MTR C-half
d
AF3: *E. coli* MTR C-half plus human MTRR ΔFMN
ipTM = 0.15 pTM = 0.58
pLDDT
Domain
Subunit
PAE Plot
MTRR$^{\Delta FMN}$
MTR C-Half
MTR Cap
MTR Cob
MTRR FAD
MTRR Conn
MTRR NADPH
MTR Act
MTRR$^{\Delta FMN}$
MTR C-half

**Supplementary Fig. 18 | AF3 prediction models of different bacterial MTR C-half and human MTRR MTR$^{\Delta FMN}$.** Predicted complexes are shown coloured by per-residue confidence scores (pLDDT, far left), by proteins domains (middle left), and by subunit (middle right). The corresponding predicted aligned error (PAE) plot for each model are shown to the far right, indicating the inter-domain confidence and the overall accuracy of the models. Shown structures are representative of the five predicted models from one AF3 run. **a,** Predicted model of human MTR C-half and human MTRR$^{\Delta FMN}$ showing a high ipTM score (0.8), suggesting stable interaction.**b,** Predicted model of *Thermus thermophilus* MTR C-half (UniProt Q9RA53) and human MTRR$^{\Delta FMN}$ (UniProt Q9UBK8) showing a low ipTM score (0.18), suggesting no stable interaction. **c,** Predicted model of *Thermus filiformis* MTR C-half (UniProt A0A0A2XCD7) and human MTRR$^{\Delta FMN}$ showing a low ipTM score (0.19), suggesting no stable interaction. **d,** Predicted model of *Escherichia coli* MTR C-half (UniProt P13009) and human MTRR$^{\Delta FMN}$ showing a low ipTM score (0.15), suggesting no stable interaction.

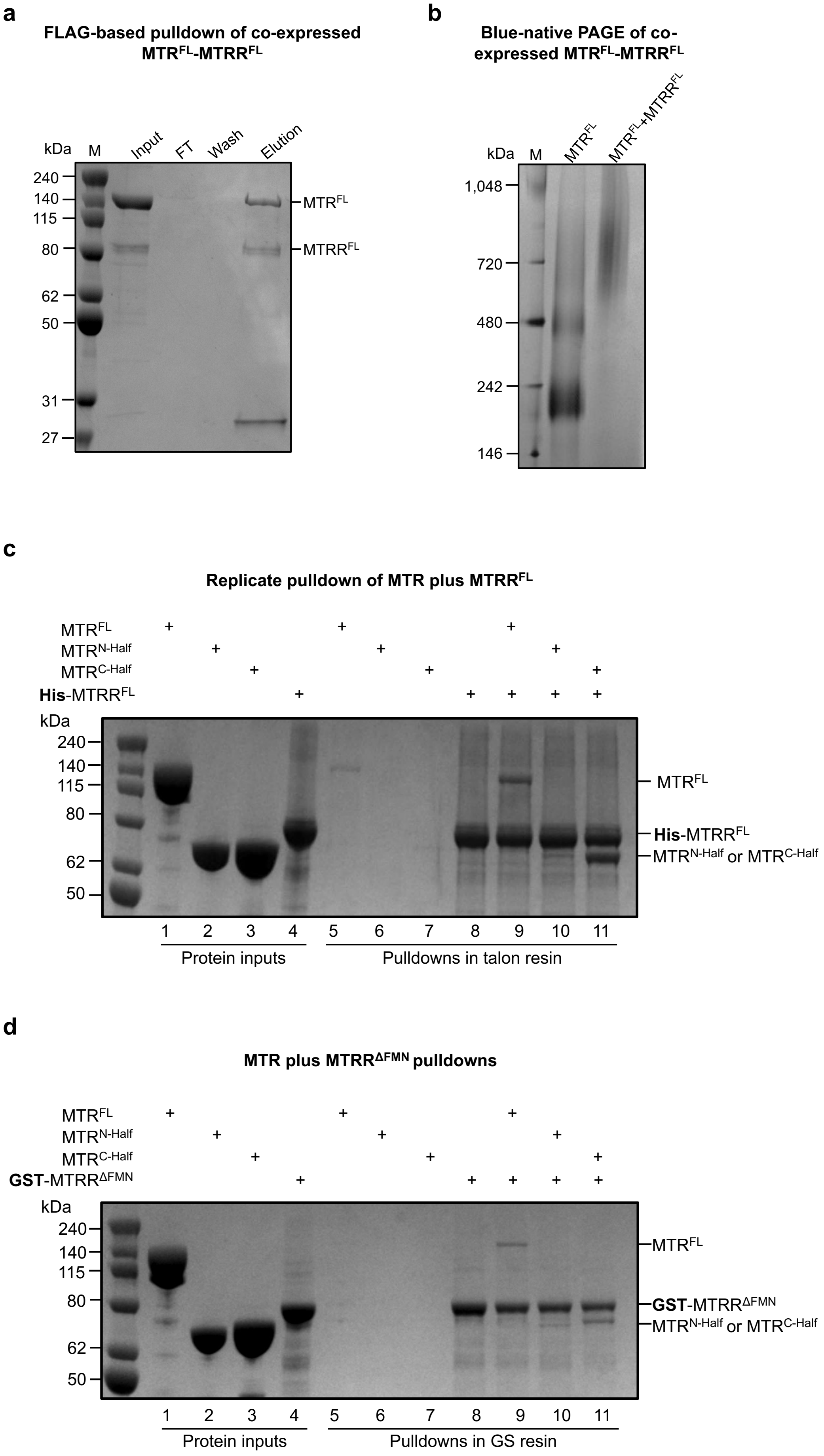
a
FLAG-based pulldown of co-expressed MTRFL-MTRRFL
kDa M Input FT Wash Elution
240
140
115
80
62
50
31
27
MTRFL
MTRRFL
b
Blue-native PAGE of co-expressed MTRFL-MTRRFL
kDa M MTRFL MTRFL+MTRRFL
1,048
720
480
242
146
c
Replicate pulldown of MTR plus MTRRFL
MTRFL
MTRN-Half
MTRC-Half
His-MTRRFL
kDa
240
140
115
80
62
50
MTRFL
His-MTRRFL
MTRN-Half or MTRC-Half
1 2 3 4 5 6 7 8 9 10 11
Protein inputs
Pulldowns in talon resin
d
MTR plus MTRRΔFMN pulldowns
MTRFL
MTRN-Half
MTRC-Half
GST-MTRRΔFMN
kDa
240
140
115
80
62
50
MTRFL
GST-MTRRΔFMN
MTRN-Half or MTRC-Half
1 2 3 4 5 6 7 8 9 10 11
Protein inputs
Pulldowns in GS resin

**Supplementary Fig. 19 | Test of complex formation of co-expressed $MTR^{FL}$ and $MTRR^{FL}$.** Co-expressed $MTR^{FL}$ and $MTRR^{FL}$ were purified and analysed to confirm complex formation. **a,** FLAG-based pulldown of the co-expressed proteins. The SDS-Page shows the molecular weight marker (M), sample input, flow-through (FT), wash, and elution fractions. Both $MTR^{FL}$ (~140 kDa) and $MTRR^{FL}$ (~80 kDa) are visible in the input and elution lanes, confirming the interaction. **b,** Blue-native PAGE of the co-expressed proteins. The $MTR^{FL}$-$MTRR^{FL}$ complex migrates at a higher apparent molecular weight than isolated $MTR^{FL}$, suggesting a stable complex formation. **c,** Replicate talon resin affinity pulldown of $MTR^{FL}$, $MTR^{N\text{-}Half}$, and $MTR^{C\text{-}Half}$ against His-$MTRR^{FL}$. Lanes 1–4 represent the protein inputs used for the assay. Lanes 5–8 represent the elution fractions of each individual protein incubated with the resin to verify their ability to bind the resin. Lanes 9-11 represent the separate pulldowns of $MTR^{FL}$, $MTR^{N\text{-}Half}$, and $MTR^{C\text{-}Half}$ against His-$MTRR^{FL}$ respectively. **d,** Replicate GS resin affinity pulldown of $MTR^{FL}$, $MTR^{N\text{-}Half}$, and $MTR^{C\text{-}Half}$ against GST-$MTRR^{\Delta FMN}$. Lanes 1–4 represent the protein inputs used for the assay. Lanes 5–8 represent the elution fractions of each individual protein incubated with the resin to verify their ability to bind the resin. Lanes 9-11 represent the separate pulldowns of $MTR^{FL}$, $MTR^{N\text{-}Half}$, and $MTR^{C\text{-}Half}$ against GST-$MTRR^{\Delta FMN}$ respectively.

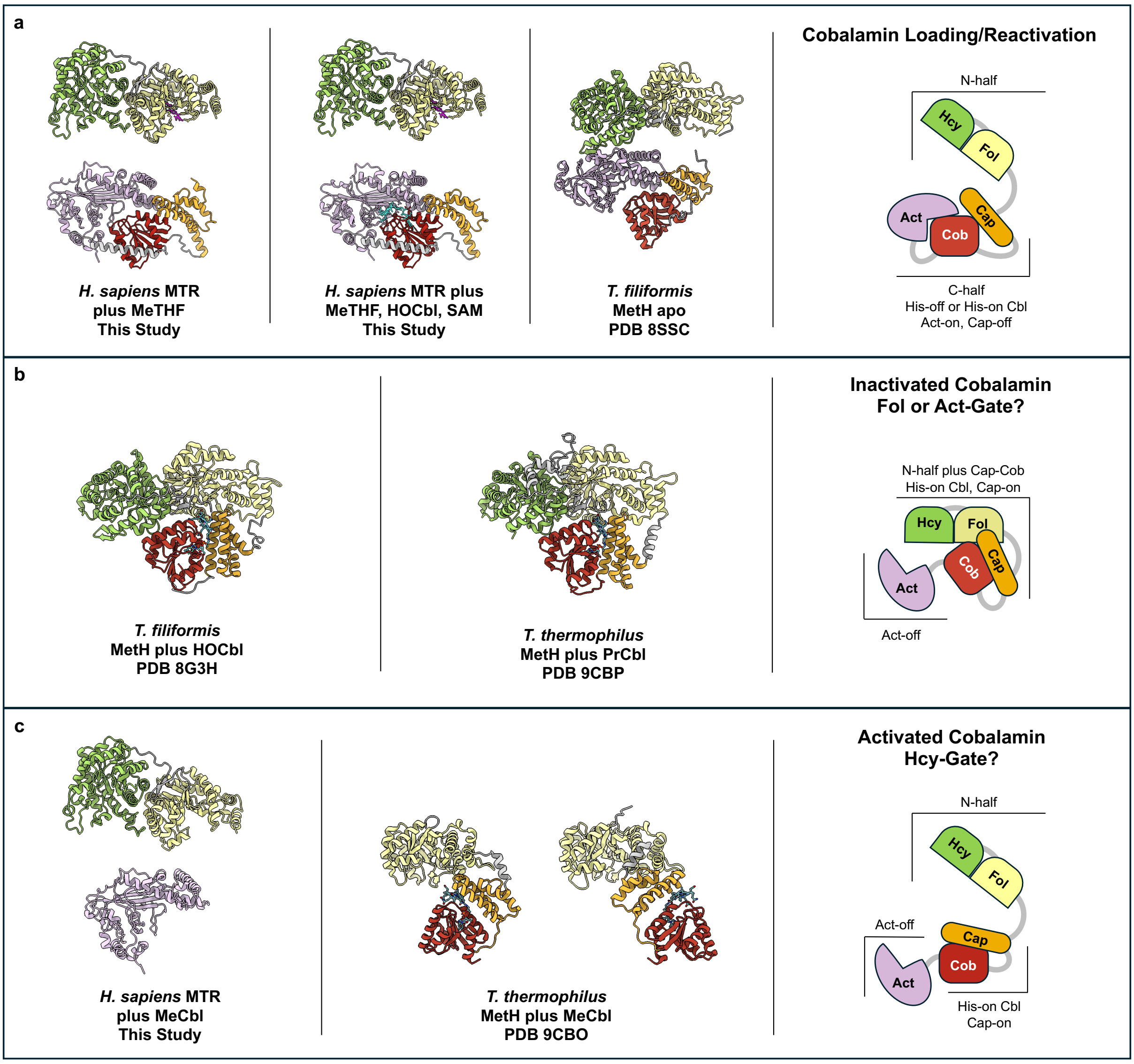


**Supplementary Fig. 20 | Comparison of human MTR structures determined in this study against recent structures of thermophilic MeTH. a,** Our structures of human MTR with MeTHF or MeTHF and HOCbl are similar to the crystal structure of full-length *T. thermophilus* (PDB 8SSC). These structures have an independent N-half and C-half modules as expected for MTR in a reactivation state. **b,** The structures of the *T. filiformis* bound to HOCbl (8G3H) and *T. thermphilus* bound to PrCbl (PDB 9CBP) are of the same conformational state of a Cap-On Act-off tetradomain arrangement. Both structures have inactive versions of Cbl. As this state has no bound folate these structures are likely representative of MTR before binding MeTHF to start a new catalytic cycle, Fol-gate, or before transforming into a reactivation conformation, Act-gate. **c,** Our structures of human MTR bound to MeCbl suggests high flexibility of the Cap-Cob module as only the Act domain was resolved of the C-half. Two structures of a tridomain *T. thermphilus* construct of the Fol-Cap-Cob domains bound to MeCbl (PDB 9CBO) also suggests a high degree of flexibility of the Cap-Cob module relative to the Fol domain. As MeCbl is the active version of Cbl used by MTR these structures are likely an active state before interacting with the Hcy domain, Hcy-gate.

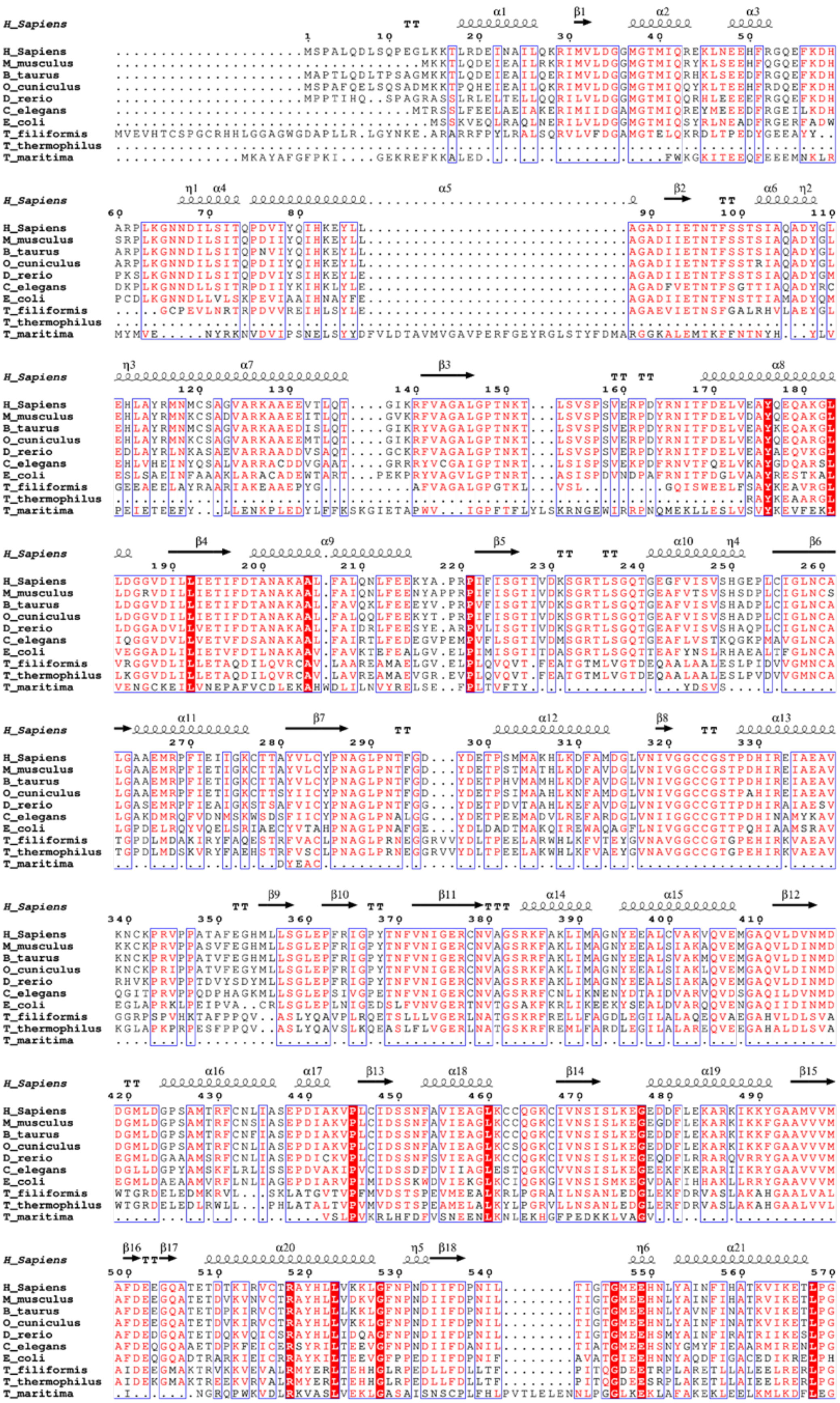
H_Sapiens
M_musculus
B_taurus
O_cuniculus
D_rerio
C_elegans
E_coli
T_filiformis
T_thermophilus
T_maritima

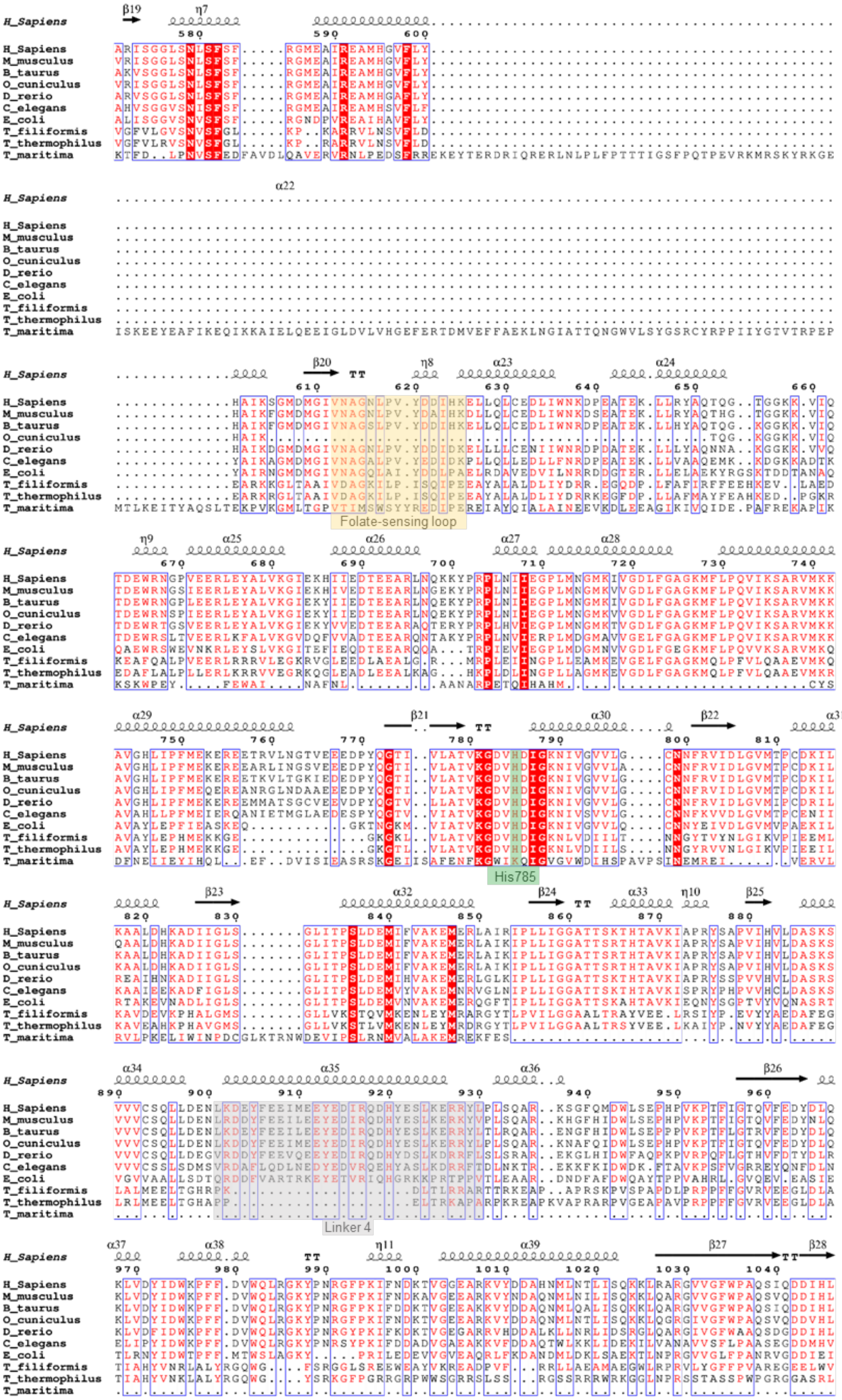
Folate-sensing loop
His785
Linker 4

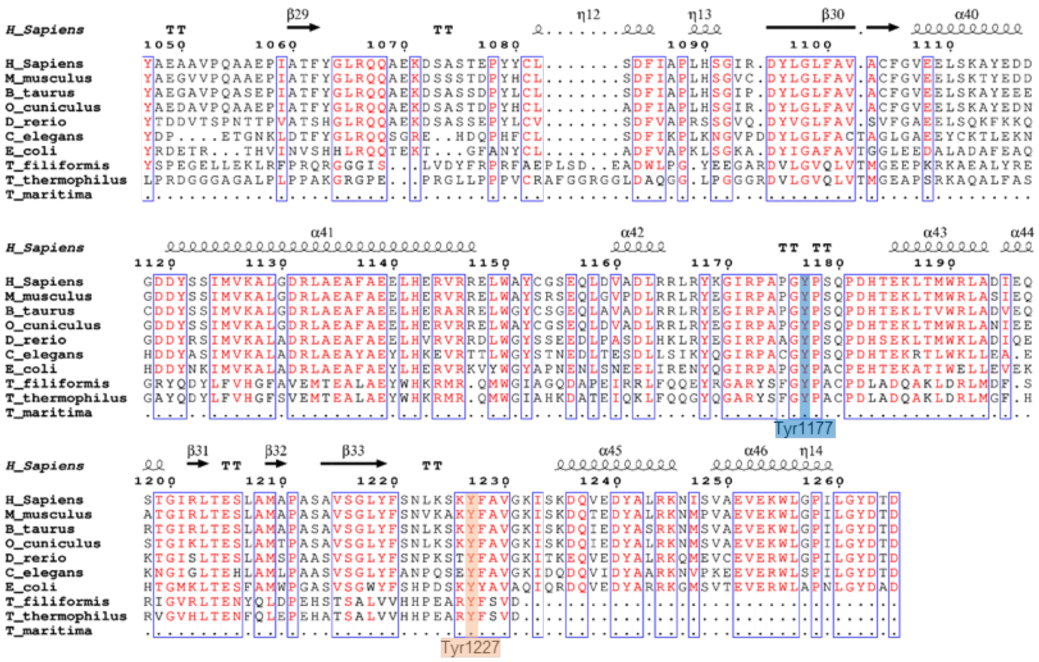

**Supplementary Fig. 21 | MTR$^{FL}$ sequence alignment.** Illustration of the secondary structure of human MTR$^{FL}$ and amino acid sequence alignment of MTR$^{FL}$ from *Mus musculus* (UniProt Q9970), *Bos taurus* (UniProt Q4JIJ3), *Oryctolagus cuniculus* (UniProt G1SXW3), *Danio rerio* (UniProt Q8JIY6), *Caenorhabditis elegans* (UniProt Q09582), *Escherichia coli* (UniProt P13009), *Thermus filiformis* (UniProt A0A0A2XCD7), *Thermus thermophilus* (UniProt Q9RA53), and *Thermotoga maritima* (UniProt Q9X112). α-helices and βsheets are shown as coils and arrows, respectively. Highlighted are the folate-sensing loop (Val612–Glu625) in yellow, histidine 785 in green, Linker 4 (Leu901–Leu930) in grey, tyrosine 1177 in blue, and tyrosine 1227 in orange. Highlighted in red are the residues conserved across all analyzed species.

### Supplementary Table 1 | Cryo-EM data collection, refinement, and validation statistics-Gatan K3 data

| | **Human Methionine Synthase With Methyltetrahydrofolate, N-Half From Full-Length** (EMDB-55190) (PDB 9SSP) | **Human Methionine Synthase With Methyltetrahydrofolate, C-Half From Full-Length** (EMDB-55191) (PDB 9SSQ) | **Human Methionine Synthase With Methyltetrahydrofolate, Hydroxocobalamin, and SAM, N-Half From Full-Length** (EMDB-55192) (PDB 9SSR) | **Human Methionine Synthase With Methyltetrahydrofolate, Hydroxocobalamin, and SAM, C-Half His-ON From Full-Length** (EMDB-55193) (PDB 9SSS) | **Human Methionine Synthase With Methyltetrahydrofolate, Hydroxocobalamin, and SAM, C-Half His-OFF From Full-Length** (EMDB-55194) (PDB 9SST) |
|---|---|---|---|---|---|
| **Data collection and processing** | | | | | |
| Magnification | 165,000 | | | 165,000 | |
| Voltage (kV) | 300 | | | 300 | |
| Electron exposure ($e^{-}/Å^2$) | 65.20 | | | 65.20 | |
| Defocus range (μm) | -0.6 to -2.0 | | | -0.6 to -2.0 | |
| Pixel size (Å) | 0.508 | | | 0.508 | |
| Symmetry imposed | C1 | C1 | C1 | C1 | C1 |
| Initial particle images (no.) | 1,638,370 | 1,638,370 | 2,195,581 | 2,195,581 | 2,195,581 |
| Final particle images (no.) | 51,788 | 100,343 | 76,292 | 25,594 | 97,889 |
| Map resolution (Å) | 3.1 | 2.9 | 2.8 | 3.1 | 2.8 |
| FSC threshold | 0.143 | 0.143 | 0.143 | 0.143 | 0.143 |
| Map resolution range (Å) | 2.5-4.7 | 2.6-5.0 | 2.5-4.7 | 2.7-8.8 | 2.5-5.1 |
| **Refinement** | | | | | |
| Initial model used | AlphaFold | AlphaFold | AlphaFold | AlphaFold | AlphaFold |
| Model resolution (Å) | 3.3 | 3.1 | 3.0 | 3.5 | 3.0 |
| FSC threshold | 0.5 | 0.5 | 0.5 | 0.5 | 0.5 |
| Map sharpening *B* factor ($Å^2$) | -106.6 | Phenix auto-sharpen | -92.8 | -80.8 | -95.2 |
| Model composition | | | | | |
| Nonhydrogen atoms | 4,925 | 4,739 | 4925 | 4876 | 4830 |
| Protein residues | 638 | 596 | 638 | 601 | 596 |
| Ligands | 1 (THH) | 0 | 1 (THH) | 1 (B12) | 1 (B12) |
| *B* factors ($Å^2$) | | | | | |
| Protein | 47.24 | 63.08 | 48.21 | 77.19 | 59.61 |
| Ligand | 44.71 | N/A | 41.20 | 66.64 | 48.34 |
| R.m.s. deviations | | | | | |
| Bond lengths (Å) | 0.003 | 0.004 | 0.003 | 0.003 | 0.002 |
| Bond angles (°) | 0.590 | 0.960 | 0.600 | 0.649 | 0.508 |
| Validation | | | | | |
| MolProbity score | 1.59 | 1.39 | 1.77 | 1.50 | 1.48 |
| Clashscore | 6.61 | 6.12 | 7.32 | 8.53 | 4.97 |
| Poor rotamers (%) | 0.38 | 1.18 | 1.52 | 0.20 | 1.97 |
| Ramachandran plot | | | | | |
| Favored (%) | 96.54 | 98.99 | 96.54 | 97.83 | 98.82 |
| Allowed (%) | 3.46 | 1.01 | 3.46 | 2.17 | 1.18 |
| Disallowed (%) | 0.00 | 0.00 | 0.00 | 0.00 | 0.00 |

### Supplementary Table 2 | Cryo-EM data collection, refinement, and validation statistics-Falcon 4i data

| | **Human Methionine Synthase With Methylcobalamin, N-Half From Full-Length** (EMDB-55195) (PDB 9SSU) | **Human Methionine Synthase With Methylcobalamin, Activation Domain From Full-Length** (EMDB-55196) (PDB 9SSV) |
|---|---|---|
| **Data collection and processing** | | |
| Magnification | 215,000 | |
| Voltage (kV) | 300 | |
| Electron exposure ($e^{-}$/Å$^2$) | 70.00 | |
| Defocus range (μm) | -0.6 to -2.0 | |
| Pixel size (Å) | 0.576 | |
| Symmetry imposed | C1 | C1 |
| Initial particle images (no.) | 1,067,416 | 1,067,416 |
| Final particle images (no.) | 175,350 | 70,000 |
| Map resolution (Å) | 2.6 | 3.1 |
| FSC threshold | 0.143 | 0.143 |
| Map resolution range (Å) | 2.6-3.3 | 2.7-4.5 |
| **Refinement** | | |
| Initial model used | AlphaFold | AlphaFold |
| Model resolution (Å) | 2.8 | 3.3 |
| FSC threshold | 0.5 | 0.5 |
| Map sharpening *B* factor (Å$^2$) | -107.6 | -120.5 |
| Model composition | | |
| Nonhydrogen atoms | 4,893 | 2716 |
| Protein residues | 638 | 338 |
| Ligands | 0 | 0 |
| *B* factors (Å$^2$) | | |
| Protein | 35.33 | 67.24 |
| Ligand | N/A | N/A |
| R.m.s. deviations | | |
| Bond lengths (Å) | 0.003 | 0003 |
| Bond angles (°) | 0.575 | 0.586 |
| Validation | | |
| MolProbity score | 1.42 | 1.13 |
| Clashscore | 4.81 | 3.36 |
| Poor rotamers (%) | 0.95 | 0.00 |
| Ramachandran plot | | |
| Favored (%) | 97.01 | 98.51 |
| Allowed (%) | 2.99 | 1.49 |
| Disallowed (%) | 0.00 | 0.00 |
